# Theoretical and practical considerations when using retroelement insertions to estimate species trees in the anomaly zone

**DOI:** 10.1101/2020.09.29.319038

**Authors:** Erin K. Molloy, John Gatesy, Mark S. Springer

## Abstract

A potential shortcoming of concatenation methods for species tree estimation is their failure to account for incomplete lineage sorting. Coalescent methods address this problem but make various assumptions that, if violated, can result in worse performance than concatenation. Given the challenges of analyzing DNA sequences with both concatenation and coalescent methods, retroelement insertions (RIs) have emerged as powerful phylogenomic markers for species tree estimation. Here, we show that two recently proposed quartet-based methods, SDPquartets and ASTRAL BP, are statistically consistent estimators of the unrooted species tree topology under the coalescent when RIs follow a neutral infinite-sites model of mutation and the expected number of new RIs per generation is constant across the species tree. The accuracy of these (and other) methods for inferring species trees from RIs has yet to be assessed on simulated data sets, where the true species tree topology is known. Therefore, we evaluated eight methods given RIs simulated from four model species trees, all of which have short branches and at least three of which are in the anomaly zone. In our simulation study, ASTRAL BP and SDPquartets always recovered the correct species tree topology when given a sufficiently large number of RIs, as predicted. A distance-based method (ASTRID BP) and Dollo parsimony also performed well in recovering the species tree topology. In contrast, unordered, polymorphism, and Camin-Sokal parsimony typically fail to recover the correct species tree topology in anomaly zone situations with more than four ingroup taxa. Of the methods studied, only ASTRAL BP automatically estimates internal branch lengths (in coalescent units) and support values (i.e. local posterior probabilities). We examined the accuracy of branch length estimation, finding that estimated lengths were accurate for short branches but upwardly biased otherwise. This led us to derive the maximum likelihood (branch length) estimate for when RIs are given as input instead of binary gene trees; this corrected formula produced accurate estimates of branch lengths in our simulation study, provided that a sufficiently large number of RIs were given as input. Lastly, we evaluated the impact of data quantity on species tree estimation by repeating the above experiments with input sizes varying from 100 to 100 000 parsimony-informative RIs. We found that, when given just 1 000 parsimony-informative RIs as input, ASTRAL BP successfully reconstructed major clades (i.e clades separated by branches > 0.3 CUs) with high support and identified rapid radiations (i.e. shorter connected branches), although not their precise branching order. The local posterior probability was effective for controlling false positive branches in these scenarios.

---

Concatenation methods continue to be widely used for estimating species trees from phylogenomic data sets. However, a potential pitfall of these methods is that they do not account for gene tree heterogeneity due to incomplete lineage sorting (ILS). Many recent studies have focused on scenarios where the species tree has consecutive short branches that put it in the anomaly zone (AZ), meaning that the most probable gene tree topology differs from the species tree topology (Degnan and Rosenberg, 2006, 2009). In such instances, concatenation can fail to recover the true species tree topology (Kubatko and Degnan, 2007) and can even be positively misleading (Roch and Steel, 2015). These results have motivated numerous authors to propose species tree estimation methods that either explicitly or implicitly account for ILS, as modeled by the multispecies coalescent (MSC) (Pamilo and Nei, 1988; Rannala and Yang, 2003). The three main approaches are (1) co-estimation methods, such as *BEAST (Heled and Drummond, 2010), that co-estimate gene trees and species trees, (2) summary methods, such as MP-EST (Liu et al., 2010) and ASTRAL (Mirarab and Warnow, 2015), that estimate species trees from (estimated) gene trees, and (3) site-based methods that estimate the species tree directly from site patterns. Many of these methods have been proven to be statistically consistent under the MSC model given their assumptions (Nute et al., 2018; Roch et al., 2019; Islam et al., 2020).

Assumptions of the standard MSC model include neutral evolution, gene tree heterogeneity that results exclusively from ILS, free recombination between loci, and no intralocus recombination (note: we refer to loci as “c-genes,” adopting the language of Doyle, 1997; Springer and Gatesy, 2016). Importantly, methods make use of these assumptions to varying degrees. For example, ASTRAL makes the assumption that the most frequent quartet (in the input set of unrooted gene trees) agrees with the unrooted species tree; this assumption holds for gene trees evolving within a species tree under the MSC model. To estimate branch lengths, ASTRAL further assumes that the frequency of quartet topologies is a good approximation for the probability that a four-taxon unrooted gene tree agrees with the four-taxon unrooted species tree, under the MSC model.

If the assumptions made by ASTRAL (or other methods) are violated, there is no guarantee that they will perform well. For summary methods, the problem of gene tree reconstruction error is especially troublesome and has been documented for numerous phylogenomic data sets (Mirarab and Warnow, 2015; Simmons and Gatesy, 2015; Springer and Gatesy, 2016, 2017, 2018; Gatesy et al., 2017, 2019; Shen et al., 2017). Indeed, Roch et al. (2019) recently showed that summary methods, including ASTRAL, can be statistically inconsistent when the number of c-genes tends toward infinity but the number of sites per c-gene is fixed.

In contrast, some site-based methods, such as SVDquartets (Chifman and Kubatko, 2014), are statistically consistent when the number of sites per c-gene is fixed at length one (Chifman and Kubatko, 2015; Wascher and Kubatko, 2020). While these methods are expected to be more robust than summary methods to problems stemming from intralocus recombination and gene tree reconstruction error, they could still be negatively impacted by model violations. For example, the current proofs of statistical consistency for SVDquartets (Chifman and Kubatko, 2015; Wascher and Kubatko, 2020) assume sites evolve down gene trees under standard models of molecular sequence evolution whose assumptions (e.g. stationarity, reversibility, and homogeneity) may be violated in practice (Naser-Khdour et al., 2019).

Given the challenges of analyzing DNA sequences with both concatenation and coalescent methods, retroelement insertions (RIs) are a promising phylogenomic marker. RIs are a broad class of “copy-and-paste” transposable elements that have been utilized for species tree estimation throughout the last two decades (e.g. Nikaido et al., 1999; Suh et al., 2015a; Lammers et al., 2019). Prior to species tree estimation, the absence (0) or presence (1) of each RI is determined for each species, producing a 0/1 character matrix. Unlike DNA sequences, homoplasy is almost unknown for RIs (Shedlock et al., 2000, 2004; Ray et al., 2006; Doronina et al., 2019), so conflicting patterns that look like homoplasy may be attributed to ILS; this is referred to as “hemiplasy” by Avise and Robinson (2008).

Shedlock (2006) suggested analyses of RIs within a population genetic framework as a promising direction for future research. This line of inquiry has been realized in recent years; for example, to detect hybridization, Kuritzin et al. (2016) modeled RIs under the MSC model with mutations occurring under a neutral infinite-sites model (henceforth we refer to this as the MSC+infinite-sites model). The infinite-sites model is appropriate, considering that a RI at a specific genomic position is a rare event (so multiple hits are unlikely), and the precise excision of a RI is also a rare event (so back mutations are unlikely) (Shedlock and Okada, 2000; van de Lagemaat et al., 2005; Doronina et al., 2019); we discuss this issue further in the discussion section. Furthermore, RIs generally occur in introns and intergenic regions of the genome (Chuong et al., 2017), which may be safe havens from selection. Indeed, Pollard et al. (2010) showed (using the statistical tests implemented within phyloP) that these regions have a much smaller percentage of sites that are under selective constraints than do protein-coding regions (i.e., introns = 2.2%, intergenic regions = 6.8%, 1st codon positions = 65.5%, 2nd codon positions = 70.8%, 3rd codon positions = 24.6%). Together these results led Springer et al. (2020) to conjecture that RIs come closer than protein-coding DNA sequences to satisfying the neutral evolution assumption.

Other studies have modeled RIs under MSC+infinite-sites model. Doronina et al. (2017) proposed a likelihood-based approach for estimating four-taxon species networks, and Mendes and Hahn (2017) provided related theoretical results, suggesting quartet-based species tree estimation as a promising direction of future research. Recently, two quartet-based methods, ASTRAL BP and SDPquartets, were introduced by Springer et al. (2020), who subsequently applied these methods to RI data sets published for Placentalia (Nishihara et al., 2009), Laurasiatheria (Doronina et al., 2017), Balaenopteroidea (Lammers et al., 2019), and Palaeognathae (Cloutier et al., 2019; Sackton et al., 2019). However, SDPquartets and ASTRAL BP have not yet been evaluated on simulated data sets where the true species tree is known. To our knowledge, the same is true of parsimony variants commonly applied to RI data sets (see Nikaido et al., 1999; Suh et al., 2015a; Lammers et al., 2019).

In this study, we evaluate methods for estimating species trees from RIs empirically as well as theoretically. First, we show that ASTRAL BP and SDPquartets are statistically consistent estimators of the unrooted species tree topology under the MSC+infinite-sites model, when the expected number of RIs per generation is constant across the species tree. Second, we benchmark these methods (as well as six other methods) on RIs simulated under the MSC+infinite-sites model from four model species trees, at least three of which are in the AZ. Half of the tested methods apply variants of parsimony (specifically unordered, Camin-Sokal, Dollo, or polymorphism parsimony) to the RIs directly. In contrast, ASTRAL BP and SDPquartets effectively estimate species trees by restricting RIs to subsets of four taxa. ASTRAL BP, in particular, encodes each RI as an unrooted tree with a single bipartition, so that the dynamic programming algorithm implemented in the efficient summary method ASTRAL (Mirarab and Warnow, 2015) can be utilized. The summary methods ASTRID (Vachaspati and Warnow, 2015) and Minimizing Deep Coalescences (Maddison and Knowles, 2006; Yu et al., 2011) can be run in similar fashion; we refer to these approaches as ASTRID BP and MDC BP, respectively.

Among the eight methods tested, ASTRAL BP is the only method that automatically estimates internal branch lengths in coalescent units (CUs) and support values, specifically the local posterior probability (PP) (Sayyari and Mirarab, 2016). As previously mentioned, the branch length estimation technique used by ASTRAL assumes the frequency with which quartets appear in the input data set is a good estimate of the probability of quartets under the MSC. This assumption does not hold for RI data sets; therefore, we modify the branch length estimation technique so that it can be used when RIs are given as input instead of gene trees.

In our simulation study, the quartet-based methods, ASTRAL BP and SDPquartets, always returned the true species tree topology when the number of parsimony-informative RIs was very large, as suggested by the theory. In this scenario, the branch lengths on the ASTRAL BP tree were accurately estimated if using the formula that we derived for RIs; if using the formula derived for gene trees, the estimated branch lengths were upwardly biased when they were greater than 0.3 CUs in the true species tree. Two other methods (ASTRID BP and Dollo parsimony) achieved similarly good topological accuracy to the quartet-based methods; in contrast, the other variants of parsimony tested (MDC BP as well as unordered, polymorphism, and Camin-Sokal parsimony) typically failed to recover the correct species tree topology in AZ situations with more than four ingroup taxa.

By repeating these experiments with 100 to 100 000 parsimony-informative RIs, we evaluated the impact of data quantity on species tree estimation. Notably, when given just 1 000 parsimony-informative RIs as input, ASTRAL BP successfully reconstructed major clades (i.e clades separated by branches > 0.3 CUs) with high support and identified rapid radiations (i.e. shorter incident branches), although not their precise branching order. The local posterior probability was effective for controlling false positive branches in these scenarios. Lastly, we compare our observations on simulated data sets to the biological data set assembled by Cloutier et al. (2019) and discuss directions for future work.

## 1 Methods

### 1.1 Model Species Trees

We simulated RIs from four model species trees, at least three of which are in the AZ (Fig. 1). Whether a model species tree is in the AZ can be determined by finding a gene tree with higher probability (under the MSC) than the gene tree that is topologically equivalent to the true species tree. To find such a tree, we examined the clade with the two shortest consecutive branches in the model species tree (denoted *𝒞*\*) and then formed alternative gene trees with symmetric resolutions of this clade. Gene tree probabilities were computed using both PhyloNet version 3.8.2 (Than et al., 2008; Yu et al., 2014; Wen et al., 2018) and PRANC (Kim et al., 2019); for details, see Section 1.1 in the Supplement, available on Github (https://github.com/ekmolloy/retrosim-study/blob/main/retrosim-supplement.pdf).

**Figure 1:**
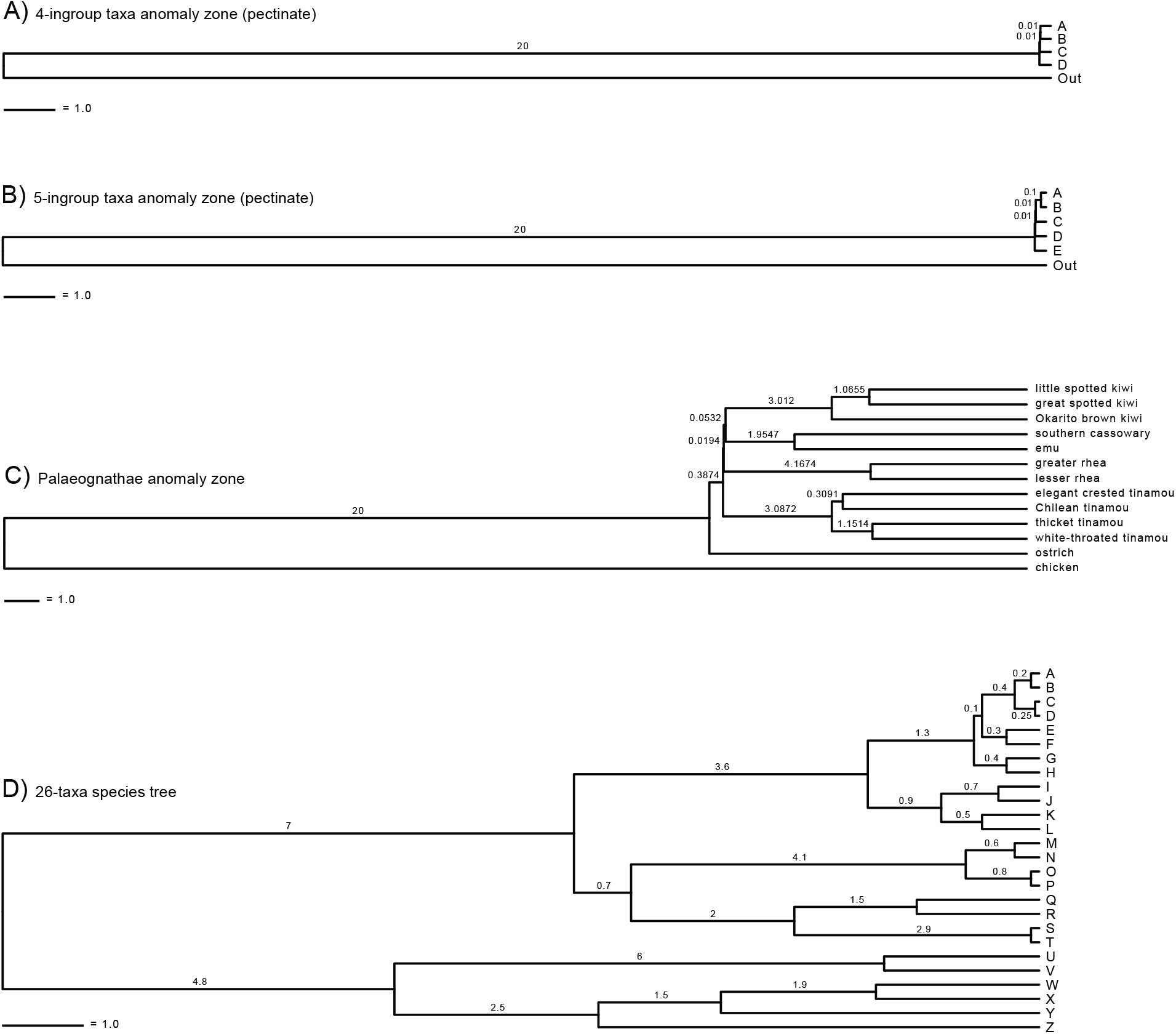
Retroelement insertion (RI) data sets were simulated from four model species trees: (A) 4-ingrouptaxa-AZ, (B) 5-ingroup-taxa-AZ, (C) Palaeognathae-GT-AZ, and (D) 26-taxa. Note that Palaeognathae-GT-AZ model species tree is based on the species tree estimated with ASTRAL given 20 850 estimated gene trees as input (Cloutier et al., 2019). Branch lengths are in coalescent units (CUs).

#### 4-ingroup-taxa-AZ

The first model species tree (Fig. 1A) is pectinate with four ingroup taxa (*A,B,C,D*) and an outgroup (*Out*). The two shallowest internal branches each have length 0.01 CUs (note: this model species tree was been studied by Mendes and Hahn, 2017). A four-taxon pectinate tree is the simplest case for anomalous rooted gene trees (Degnan and Rosenberg, 2006), and a sufficient condition for this species tree to be in the AZ occurs when both internal branches in the ingroup clade are less than 0.1568 CUs (Degnan and Rosenberg, 2006). This condition is met. Furthermore, we find that gene trees with clades *𝒞*\* = (*D*, (*C*, (*B, A*))) and *𝒞* = ((*D, C*), (*B, A*)) have probabilities 0.0600 and 0.1133, respectively.

#### 4-ingroup-taxa-AZ

The second model species tree (Fig. 1B) is a pectinate tree with five ingroup taxa (*A,B,C,D,E*) and an outgroup (*Out*). The shallowest internal branch has length 0.1 CUs, and the other two internal branches in the ingroup clade each have length 0.01 CUs (note: this model species tree was studied by Rosenberg and Tao, 2008 and Degnan and Rosenberg, 2009). Gene trees with clades *𝒞*\* = (*E*, (*D*, (*C*, (*B, A*)))), *𝒞*_1_ = ((*E, D*), (*C*, (*B, A*))), and *𝒞*_2_ = (*E*, ((*D, C*), (*B, A*))) have probabilities 0.0113, 0.0265, and 0.0161, respectively; therefore, this model species tree is in the AZ.

#### Palaeognathae-GT-AZ

The third model species tree (Fig. 1C) is based on the species tree estimated by Cloutier et al. (2019) by running ASTRAL on gene trees estimated from 20 850 loci (12 676 CNEEs, 5 016 introns, 3 158 UCEs). In order to create a model species tree, we rooted this estimated tree at the outgroup (chicken) and set the lengths of terminal branches so that the resulting tree was ultrametric. There are two short consecutive internal branches: a branch with length 0.0532 CUs separating emu, cassowary, and kiwi from the other taxa, and a branch with length 0.0194 CUs separating emu, cassowary, kiwi, and rhea from the other taxa. Gene trees with clades *𝒞* * = ((((*emu, cassowary*), *kiwi*), *rhea*), *tinamou*) and *𝒞* = (((*emu, cassowary*), *kiwi*), (*rhea, tinamou*)) have probabilities 0.006 and 0.009, respectively; therefore, this model species tree is in the AZ.

Again, note that this model species tree, referred to as Palaeognathae-GT-AZ, is based on Cloutier et al.’s (2019) ASTRAL analysis of *gene trees*. Cloutier et al. (2019) also assembled a data set of *RIs* on which Springer et al. (2020) estimated a species tree using ASTRAL BP; we refer to the result as the Palaeognathae-RI tree. The Palaeognathae-RI tree has the same topology as the Palaeognathae-GT-AZ tree; however, it is *not* in the AZ. These two trees, Palaeognathae-GT-AZ and Palaeognathae-RI, should not be confused with each other, as they were estimated from different types of data (gene trees versus RIs).

#### 26-taxa

The fourth model species tree (Fig. 1D) has 26 taxa and includes a wider range of internal branch lengths for evaluating the accuracy of branch length estimation with ASTRAL BP. This model tree contains two short consecutive internal branches: a branch with length 0.3 CUs separating *E* and *F* from the other taxa, and a branch with length 0.1 CUs separating *A,B,C, D, E*, and *F* from the other taxa. Gene trees with clades *𝒞*\* = (((*AB, CD*), *EF*), *GH*) and *𝒞* = ((*AB, CD*), (*EF, GH*)) have probabilities 0.000143 and 0.000130, respectively. While this model species tree is not necessarily in the AZ, it is still a challenging model condition.

### 1.2 Data sets

#### Simulated data sets

RIs were simulated with the *ms* program (Hudson, 2002), which enables coalescent simulations with 0/1 mutations following a neutral infinite-sites model. Our simulations further assume free recombination among loci, no intralocus recombination, neutrality, no missing data, constant effective population size, and a uniform rate of RIs per unit length of the species tree. We simulated 25 replicate data sets from each of the four model species trees (Fig. 1) with one segregating site for each locus (note: species tree branch lengths were halved prior to simulating data, because the *ms* program uses a currency of 4*N* generations per unit for species tree and gene tree branch lengths, whereas a CU is 2*N* generations for a population of diploid individuals). Given that we were interested in whether different species tree estimation methods converge on the correct species tree, we simulated 25 replicate data sets that were sufficiently large so that they could be pruned to contain exactly 100 000 parsimony-informative RIs (note: we say that a RI is parsimony-informative if it contains at least two 0’s and at least two 1’s). The *ms* commands are provided in Section 1.2 of the Supplement.

### 1.3 Species Tree Estimation Methods

We estimated species trees from RIs using eight different methods: four variants of parsimony, two quartet-based methods, a distance-based method, and a method based on minimizing deep coalescences. The first six were selected because they had been used to infer species trees from RIs in prior studies; the last two were selected as candidate methods for species tree inference from 0/1 characters because they are popular summary methods that accept unrooted and non-binary gene trees as input. Details, including software commands, can be found in Section 1.3 of the Supplement.

#### Parsimony methods for 0/1 characters

Four variants of parsimony (unordered, Camin-Sokal, Dollo, and polymorphism) were used to estimate species trees directly from RI site patterns; we provide a brief overview of these methods, referring the interested reader to Felsenstein (1983) for further discussion.

Unordered parsimony (Fitch, 1971) applies equal weights to “forward” changes (0 to 1) and “backward” changes (1 to 0). The other variants of parsimony evaluated here treat these changes differently, assuming 0 is the ancestral state and 1 is the derived state. Camin-Sokal parsimony (Camin and Sokal, 1965) allows forward changes only (so homplasy is explained by parallel evolution); in contrast, Dollo parsimony (Farris, 1977) allows at most one forward change followed by any number of backward changes (so homoplasy is explained by reversals). Polymorphism parsimony (Felsenstein, 1979) allows at most one forward change after which the derived allele may co-exist with the ancestral allele (so homoplasy is explained via polymorphism). For a given site pattern, each node in the tree can be labeled polymorphic (meaning the two states co-exist) or monomorphic (implying that one of the two states has been fixed or that the node is ancestral to the origin of the derived allele). Polymorphism parsimony seeks to minimize the extent of polymorphism across the tree (Felsenstein, 1983), whereas the other three variants seek to minimize the total number of character state changes.

Analyses with unordered, Dollo, and Camin-Sokal parsimony were performed using PAUP* v4a168 (Swofford, 2002). We used branch-and-bound searches for all data sets with the exception of the 26-taxon data set, in which case we employed heuristic searches (specifically taxon addition followed by tree-bisection and reconnection moves) for 100 random orders of taxa. In either case, PAUP* returns all trees with the same parsimony score, and we used the strict consensus of the returned trees to evaluate species tree topological accuracy. Analyses with polymorphism parsimony were performed using the dollop program in PHYLIP v3.695 (Felsenstein, 1989) with the jumble option set to 50 (note: this was the only program run manually in this study).

#### Quartet-based methods

SDPquartets (Springer et al., 2020) is a quartet-based method that was developed for low-homoplasy 0/1 characters such as RIs. In the first step of SDPquartets, an unordered parsimony analysis is performed for RIs, restricted to all subsets of four species. In the second step, the optimal trees on four taxa are assembled into a species tree on the full set of taxa using Matrix Representation with Parsimony (MRP) (Ragan, 1992). We used branch-and-bound searches when running the unordered parsimony analysis of the MRP matrix to ensure recovery of all most parsimonious trees. For our analyses with SDPquartets, we used a custom Perl script (https://github.com/dbsloan/SDPquartets) that directs PAUP* (Swofford, 2002) to perform both steps of the species tree estimation pipeline.

Because the SDPquartets pipeline enumerates all subsets of four species, it will not be feasible to run this approach on data sets with very large numbers of species. To improve computationally efficiency, we encode RIs as unrooted trees with a single bipartition (BP) and then apply ASTRAL, which implements a dynamic programming algorithm. This pipeline, referred to as ASTRAL BP, was proposed by Springer et al. (2020). For our analyses with ASTRAL BP, we used a custom Python script (https://github.com/ekmolloy/retrosim-study/blob/main/tools/run_astral_bp.py) and ASTRAL version 5.7.5 (i.e. ASTRAL-III as described by Zhang et al., 2018).

ASTRAL BP automatically annotates the internal branches of the species tree with branch lengths in CUs and branch support (local PP) using the algorithms introduced by Sayyari and Mirarab (2016). Specifically, Sayyari and Mirarab (2016) derived the maximum a posteriori (MAP) estimate and maximum likelihood estimate (MLE) of the branch length for unrooted gene trees, denoted MAP-GT and MLE-GT, respectively. In this study, we derived the MLE of the branch length for RIs, denoted MLE-RI, and then implemented this calculation within our custom Python script (mentioned above).

#### Other “BP” methods

Because we encoded RIs as unrooted trees with a single bipartition to run ASTRAL BP, we applied other (gene tree) summary methods, specifically ASTRID v2.2.1 (Vachaspati and Warnow, 2015) and minimizing deep coalescence (Maddison and Knowles, 2006), to the same data sets. These methods are referred to as ASTRID BP and MDC BP, respectively.

ASTRID is a distance-based method that operates by computing the number of internodes between all pairs of taxa in an unrooted gene tree, averaging this value across all input gene trees, and then running FastME (Lefort et al., 2015) on the resulting dissimilarity matrix. In the context of RI data sets, this approach is equivalent to running FastME given uncorrected p-distances as input.

MDC is a parsimony-based approach that traditionally infers a species tree from a set of rooted gene trees; however, there is also a version of MDC for unrooted and non-binary gene trees (Yu et al., 2011) implemented within PhyloNet v3.8.2 (Than et al., 2008; Wen et al., 2018). We allowed MDC to return an unresolved species tree (option: “-ur”), which is recommended when the input gene trees are unresolved.

### 1.4 Method Evaluation

Species trees were estimated on data sets with 100 000 parsimony-informative RIs. Then, to evaluate the impact of data set size, methods were run on data sets from two model species trees (Palaeognathae-GT-AZ and 26-taxa) restricted to the first 100, 500, 1 000, 5 000, 10 000, and 50 000 RIs.

#### Analyses of species tree topological accuracy

For data sets with 100 000 RIs, we report specific differences between the estimated and true species tree topology across all replicate data sets. For data sets with varying numbers of RIs, we report species tree error as the false negative rate (i.e. the number of internal branches in the true species tree that are missing from the estimated tree, divided by the number of internal branches in the true species tree). We also explored the utility of the branch support metric computed by ASTRAL BP by collapsing branches with local PP below a given threshold and reporting the FN rate as well as the false positive rate (i.e. the number of internal branches in the estimated species tree that are missing from the true tree, divided by the number of internal branches in the true tree).

#### Analyses of specific branches in model species trees

Species tree error is typically reported globally (i.e. across all branches in the tree); however, we also explored performance for some specific branches in the Palaeognathae-GT-AZ and the 26-taxa model species trees. We looked at the proportion of replicates (out of 25) recovering the branch, the accuracy of the estimated branch length (MLE-RI only), and the effective number (*EN*) of RIs around the branch (i.e. the number of RIs with topological information for resolving a branch; this quantity is approximated as discussed by Sayyari and Mirarab, 2016). Branch length accuracy was measured as the absolute error (i.e. the absolute value of the difference between the true and estimated branch length) as well as the percent error (i.e. the absolute error divided by the length of the true branch, multiplied by 100).

#### Analyses of Palaeognathae trees

Recall that the Palaeognathae-GT-AZ tree (which was estimated by running ASTRAL on 20 850 gene trees) was in the AZ, whereas the Palaeognathae-RI tree (which was estimated by running ASTRAL BP on 4 301 parsimony-informative RIs) was not. To determine whether it would be possible to detect whether a species tree was in the AZ from a relatively small number of RIs, we evaluated the performance of ASTRAL BP on data sets simulated from the Palaeognathae-GT-AZ tree with 1 000 and 5 000 RIs. For each estimated species tree, we evaluated whether it (1) resolved major clades (i.e clades separated by branches *>* 0.3 CUs) correctly and with high support, (2) contained short branches indicative of a rapid radiation (i.e. shorter connected branches), (3) was in the AZ, and (4) recovered low support values for any incorrect branch. This was achieved through several analyses performed separately for replicate data sets with 1 000 or 5 000 RIs. First, for each internal branch in the Palaeognathae-GT-AZ model species tree, we counted the number of replicates (out of 25) recovering the branch and computed the average (*±* standard deviation) length and local PP for the branch. We also computed the average length and local PP for any incorrect branch. Second, we identified all species tree topologies recovered across the 25 replicates; these topologies were visualized with average branch lengths and support values. By looking at the visualizations, we were able to inspect whether there was a rapid radiation in each estimated tree and to manually construct alternative topologies to test whether the estimated species tree was in the anomaly zone (note: this test was performed using PhyloNet as discussed above).

## 2 Theoretical Results

We begin this section by showing that SDPquartets is statistically consistent; the result for ASTRAL BP is in the Appendix.

There are six parsimony-informative site patterns for taxon set {*A, B, C, D*}; patterns 1100 and 0011 both correspond to quartet *AB*|*CD*, patterns 1010 and 0101 both correspond to quartet *AC*|*BD*, and patterns 1001 and 0110 both correspond to quartet *AD*|*BC*. Suppose that site patterns encode the absence (0) or presence (1) of a RI and that RIs are generated under the MSC+infinite-sites model, parameterized by a rooted species tree topology on {*A, B, C, D*}, with each branch annotated by the time in generations, the effective population size, and the probability of a new insertion for each allele in the population (note: the latter two parameters must also be specified for the population above the root). Doronina et al. (2017) derived the expected number of RIs displaying each of the six patterns listed above when RIs are generated from four-taxon species networks under the MSC+infinite-sites model (note: they used the diffusion approximation to the neutral Wright-Fisher coalescent model; see Fisher, 1922; Wright, 1931; Kimura, 1955a,b). Using their derivations, we computed the probability that a RI has pattern 1100 conditioned on it having one of the six parsimony-informative patterns listed above; this probability is denoted *P* (1100). After repeating this calculation for the other five parsimony-informative site patterns, we showed that

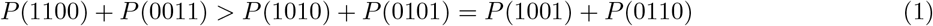

for a model species tree with either a pectinate topology (((*A, B*), *C*), *D*) or a balanced topology ((*A, B*), (*C, D*)), provided that the length (in CUs) of the internal branch is positive. Put simply, the probability that a parsimony-informative RI corresponds to the model species tree topology, denoted *P*_*RI*_ (*AB*|*CD*) = *P* (1100)+ *P* (0011), is strictly greater than the probability that it corresponds to one of the two alternative quartet topologies, denoted *P*_*RI*_ (*AC*|*BD*) = *P* (1010) + *P* (0101) and *P*_*RI*_ (*AD*|*BC*) = *P* (1001) + *P* (0110). This is Theorem 1 in the Supplemental Text (also see Mendes and Hahn, 2017). We use this result to show that SDPquartets and ASTRAL BP are statistically consistent.

### Theorem 1.

Suppose that RIs are generated under the MSC+infinite-sites model (as approximated by Doronina et al., 2017) and that the expected number of new RIs per generation is constant across the species tree. Then, SDPquartets is statistically consistent estimator of the unrooted species tree topology.

**Proof Sketch.** Let *T* * be the true species tree on taxon set *S*, and suppose that |*S*| = 4. For each of the three possible quartet topologies on taxon set *S*, denoted *t*_1_, *t*_2_, *t*_3_, the unordered parsimony score is computed as

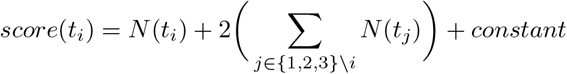

where *N* (*t*_*k*_) is the number of RIs that correspond to quartet topology *t*_*k*_ for *k* = {1, 2, 3} and *constant* is the number of RIs that contain both 0’s and 1’s but are not parsimony-informative (and thus do not correspond to any of the three quartet topologies). The quartet topology with the lowest score is selected. Let *n* = *N* (*t*_1_) + *N* (*t*_2_) + *N* (*t*_3_) be the number of RIs that are parsimony-informative. As the total number of RIs goes to infinity, *n* also goes to infinity (i.e. *n* is not bounded). Then, by Theorem 1 in the Supplemental Text (which says the most probable quartet agrees with the species tree and the two alternative quartets have equal probability), unordered parsimony selects the true species tree *T* * with high probability.

Now suppose that |*S*| > 4. In this case, SDPquartets operates in two phases. In the first phase, SDPquartets restricts the RIs to each subset of four taxa, applies unordered parsimony to the restricted RIs, and then adds the quartet topology with the lowest score to a set 𝒬. As the number of RIs goes to infinity, with high probability, every quartet *q* ∈ 𝒬 will be topologically equivalent to the true species tree *T* *; in this scenario *T* * is the unique compatibility supertree for 𝒬 (see Sections 3.2.1, 7.2, and 7.5 in Warnow, 2017 for details). In the second phase, SDPquartets runs the supertree method Matrix Representation with Parsimony (Ragan, 1992) on 𝒬. Any optimal solution to MRP is a (refined) compatibility supertree for 𝒬, provided that a compatibility supertree for 𝒬 exists (Theorem 7.8 in Warnow, 2017). Therefore, as the number of RIs goes to infinity, the optimal solution to MRP (given 𝒬 as input) will be *T* * with high probability.

MRP is an NP-hard optimization problem; however, branch-and-bound algorithms (Hendy and Penny, 1982), which are guaranteed to find the optimal solution, can be utilized whenever the number of taxa is sufficiently small. Therefore, SDPquartets (using a branch-and-bound algorithm for MRP) will return the true species tree *T* * with high probability, as the number of RIs goes to infinity.

The proof of statistical consistency of ASTRAL BP is closely related to the proof of statistical consistency for ASTRAL (Theorem 2 in Mirarab et al., 2014), so we provide the proof in the Appendix (Theorem 2).

### 2.1 Branch Length and Support Estimation

The gene tree summary method ASTRAL not only estimates the unrooted species tree topology but also the lengths of the internal branches in CUs (Sayyari and Mirarab, 2016). For an estimated unrooted species tree *T* on taxon set {*A, B, C, D*}, the estimation of the internal branch length is based on quartet frequencies, that is, the number *z*_1_ of unrooted gene trees, out of *n*, that display the same topology as *T*. Assuming the branch in question exists in the true unrooted species tree *T* *, the MLE of its length, referred to as MLE-GT, is 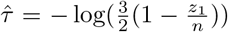 (Theorem 2 in Sayyari and Mirarab, 2016). This follows from their statistical framework (reviewed in the Appendix) and from the probability of unrooted gene trees under the MSC:

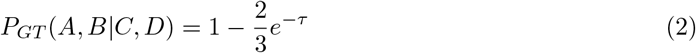

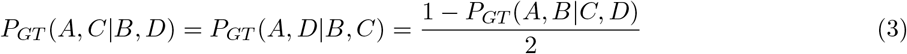

where *τ* is the length (in CUs) of the internal branch inducing *A, B*|*C, D* in the unrooted model species tree (Section 4.1 in Allman et al., 2011). As *A, B* |*C, D* agrees with the species tree, we refer to it as the “dominant quartet”; we refer to *A, C* |*B, D* and *A, D* |*B, C* as the “alternative quartets.”

The statistical framework proposed by Sayyari and Mirarab (2016) can be applied to RIs (Appendix); however, the formula for the probability of the dominant quartet is more complicated and depends on whether the model species tree is pectinate or balanced (see Section 2 of the Supplemental Text for details). When internal branches of the model species tree are short enough so that the small angle approximation *e*^−*τ*^ = 1 − *τ* can be applied, the probability of the dominant and alternative quartets simplifies to

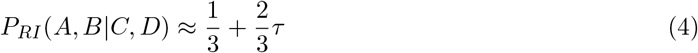

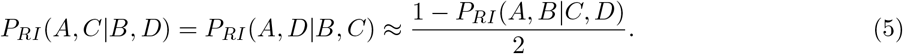

for both the pectinate and balanced species tree. Applying the small angle approximation to the formula for unrooted gene trees (Equation 2) also yields Equation 4; therefore, provided that the internal branches are sufficiently short (*<* 0.3 CUs), the MLE-GT formula can be used to estimate lengths, even when RIs are given as input (Figure 2).

**Figure 2:**
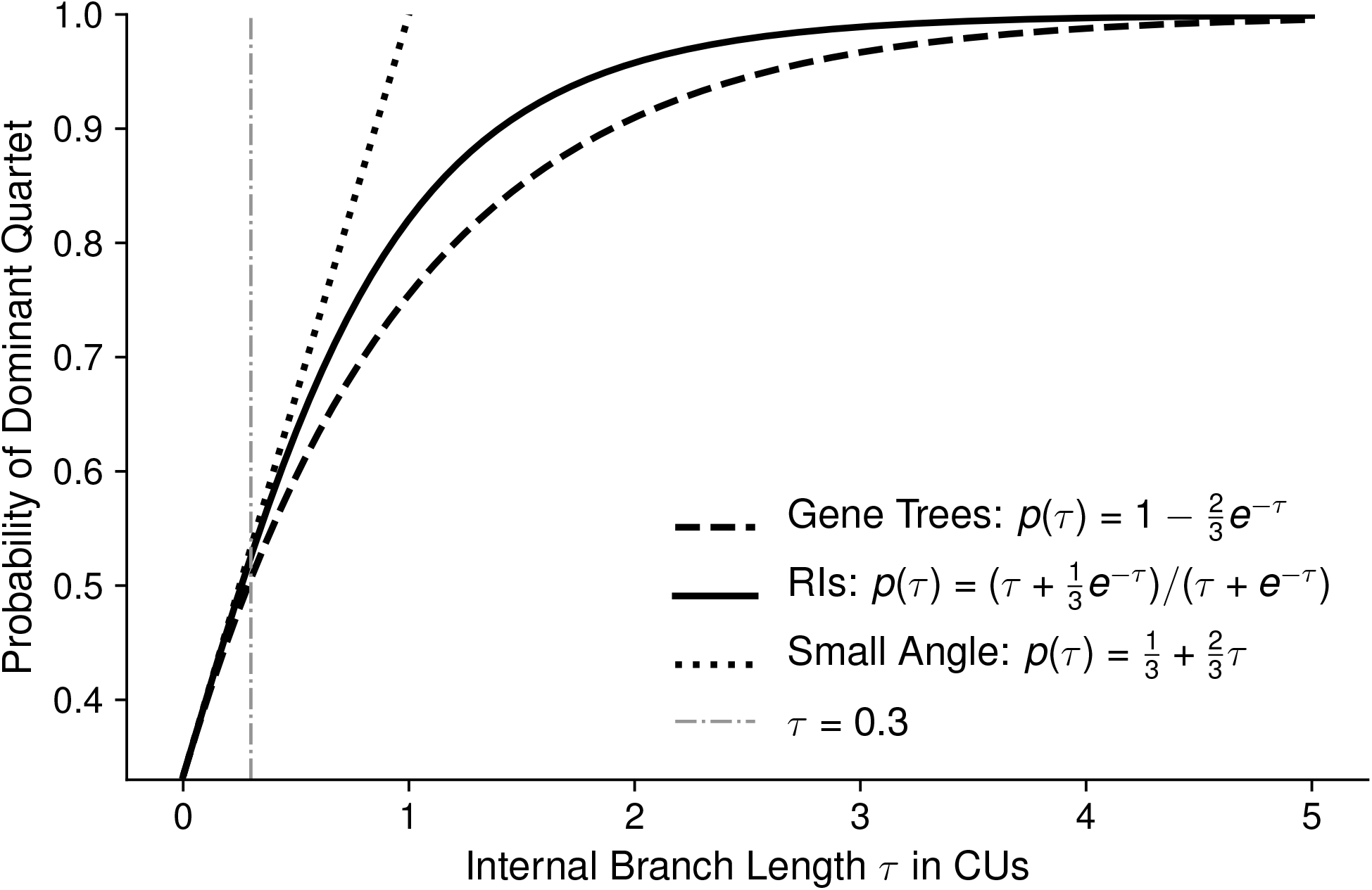
Relationship between the internal branch lengths and the probability of the dominant quartet (i.e. the quartet that agrees with the species tree). The formula for gene trees is under the MSC model (Allman et al., 2011). The formula for retroelement insertions (RIs) is under the MSC+infinite-sites model, assuming that the expected number of new RIs per generation is constant across the species tree. When the internal branch length *τ* is sufficiently short so that we can use the small angle approximation *e*^−*τ*^ = 1 −*τ*, both the equation for gene trees and the equation for RIs simplifies to the “small angle” equation above.

If we assume that the expected number of new RIs per generation is constant across the species tree, the probability of the dominant quartet simplifies to

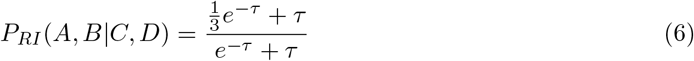

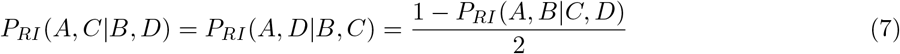

for both the pectinate and balanced species tree (see Appendix for details). Then, using the statistical framework proposed by Sayyari and Mirarab (2016), we show that the MLE of the branch length, referred to as MLE-RI, is

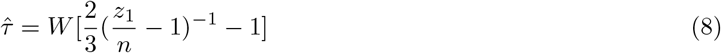

where W is Lambert’s W (Theorem 3 in the Appendix). The MLE does not exist 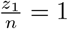 (i.e. there is no conflict); in this case, we set the estimated branch length 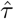 to by default.

Lastly, ASTRAL provides a measure of branch support: the local PP. This calculation requires us to place a prior on the probability of the dominant quartet, which could be viewed as placing a prior on the branch lengths in the species tree. The local PP calculation implemented within ASTRAL places a prior on branch lengths that corresponds to the species tree being generated under a Yule process with birth rate *λ* (Lemma 2 in Sayyari and Mirarab, 2016). For 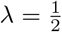 (the default in ASTRAL), this is equivalent to placing a uniform prior on the probability of the dominant quartet. When RIs are given as input instead of gene trees, the interpretation (regarding the generation of the species tree under the Yule process) does not hold; see Appendix for details. Nevertheless, it is reasonable to place a uniform prior on the probability of the dominant quartet in the absence of other information.

## 3 Simulation Study

We now describe the results of benchmarking ASTRAL BP, SDPquartets, and six other methods on simulated data sets.

### 3.1 4-ingroup-taxa-AZ

For the 4-ingroup-taxa-AZ model species tree, all methods (with the exception of Camin-Sokal parsimony) returned the correct species tree topology for all 25 replicate data sets, each with 100 000 parsimony-informative RIs. Camin-Sokal parsimony always recovered an incorrect position for taxon *C*, placing it as the sibling of *D* instead of as the sibling of the clade (*A, B*) (Fig. 3A). Considering the probabilities of rooted gene trees under the MSC, the (incorrect) topology returned by Camin-Sokal is more probable than the true species tree topology (see Methods section for details).

**Figure 3:**
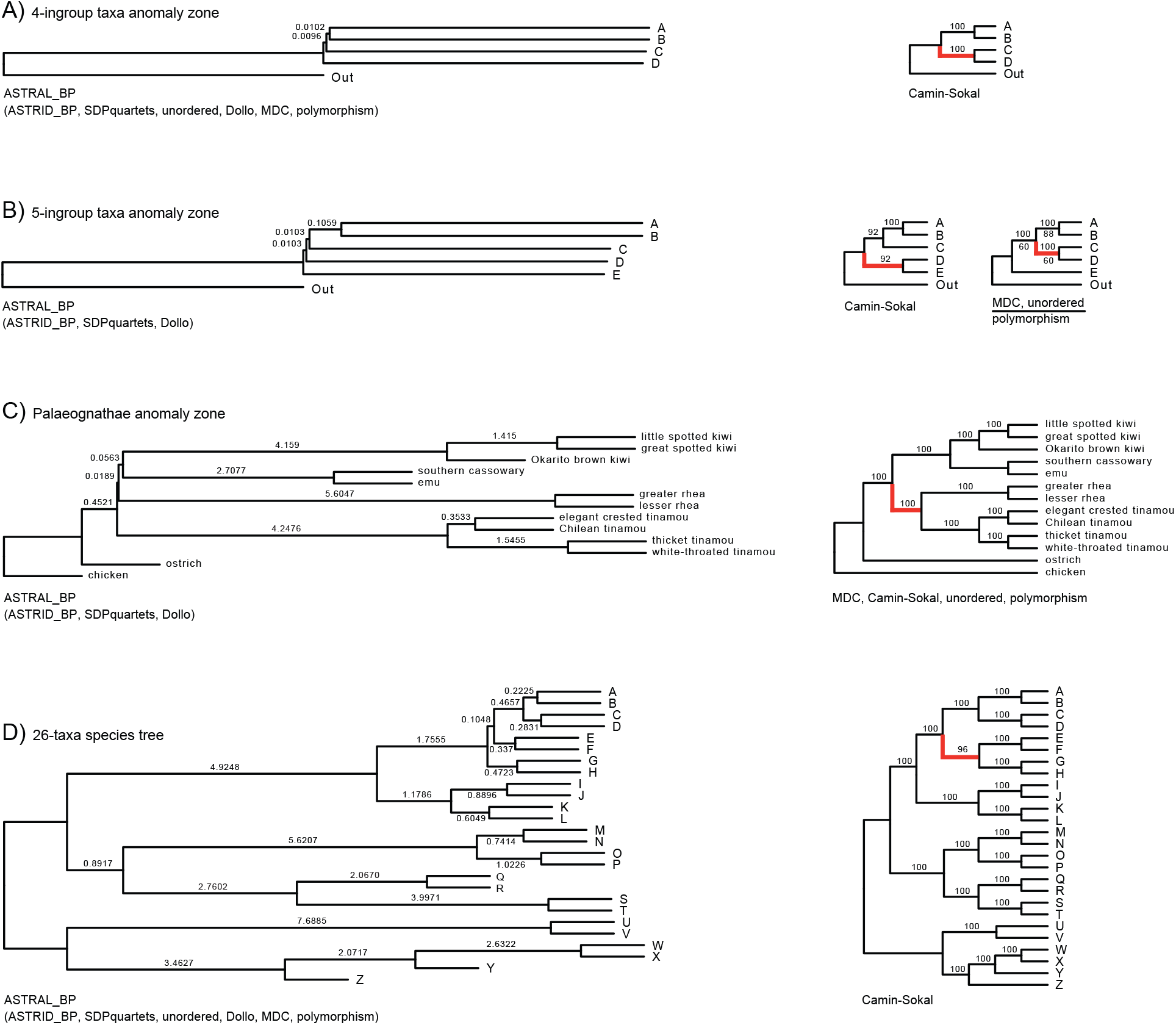
Summary of results for eight different phylogeny reconstruction methods (ASTRAL BP, ASTRID BP, SDPquartets, Camin-Sokal parsimony, Dollo parsimony, MDC BP, polymorphism parsimony, and unordered parsimony) that were employed to estimate species trees for 25 replicate data sets simulated from four model species trees: (A) 4-ingroup-taxon-AZ, (B) 5-ingroup-taxon-AZ, (C) Palaeognathae-GT-AZ, and (D) 26-taxa. ASTRAL BP always recovered the correct species tree, which is shown on the left side, with branches annotated with the default branch lengths (in coalescent units) averaged across the 25 replicate data sets. The default branch lengths use the maximum a posteriori (MAP) formula derived for gene trees; see Table S1 in the Supplement for maximum likelihood (ML) branch lengths using the formula derived for gene trees and retroelement insertions (RIs). The other methods that recovered the correct species tree for all 25 replicates are shown in parentheses on the left side; there is one exception: Dollo parsimony recovered the correct tree for Palaeognathae-GT-AZ in 22 of 25 simulations but is still shown in parentheses on the left side. For methods that recovered incorrect species trees, their majority-rule consensus species tree (across the 25 replicates) is shown on the right side; red branches are those that conflict with the true species tree. On each internal branch, the numbers above and below indicate the percentage of replicates for which the branch was reconstructed by the methods above and below the lines, respectively.

The MAP-GT branch lengths, averaged across the 25 ASTRAL BP trees, had values of 0.0096 and 0.0102 CUs for branches *x* (0.01 CUs) and *y* (0.01 CUs), respectively (Fig. 3A). The absolute differences between the true and estimated (MAP-GT) branch lengths were small, with mean error of 0.0021 (21%) and 0.0025 (25%), respectively. The mean branch lengths were the same regardless of the approach (MAP-GT, MLE-GT, or MLE-RI) used for estimation (Table S1 in Supplement).

### 3.2 5-ingroup-taxa-AZ

In contrast to the previous model condition, only four of the eight methods (ASTRAL BP, ASTRID BP, SDPquartets, Dollo parsimony) recovered the correct species tree topology when given 100 000 RIs simulated from 5-ingroup-taxa-AZ model species tree. The other methods (Camin-Sokal parsimony, polymorphism parsimony, unordered parsimony, and MDC BP) always recovered incorrect species trees that were not fully pectinate. For these methods, we computed the majority consensus trees across the 25 replicates (Fig. 3B). All of the resulting topologies have higher probability than the model species tree topology, when considering the probabilities of rooted gene trees under the MSC model (see Methods section for details).

For branch *x* (0.01 CUs), branch *y* (0.01 CUs), and branch *z* (0.1 CUs), ASTRAL BP recovered MAP-GT branch lengths with mean values of 0.0103, 0.0103, and 0.1059 CUs, respectively (Fig. 3B). The mean error for these branches was small: 0.0018 (18%), 0.0018 (18%), and 0.0060 (6%), respectively, and the estimated branch lengths computed with different approaches (MAP-GT, MLE-GT, MLE-RI) were quite similar on average (Table S1 in Supplement).

### 3.3 26-taxa

#### Results for 100 000 RIs

For the 26-taxa model species tree, all methods except Camin-Sokal parsimony estimated the correct species tree for all 25 replicate data sets (Fig. 3D) when given 100 000 parsimony-informative RIs. Camin-Sokal parsimony recovered an incorrect phylogeny for 24 out of 25 replicates; this topology had clade (*E, F*) misplaced as the sibling of clade (*G, H*). Considering the probabilities of rooted gene trees under the MSC, the alternative topology typically recovered by Camin-Sokal was only slightly less probable than the true species tree topology (see Methods section for details).

The conditioning of the ML branch length estimation problem worsens as the branch length increases (Table S2 in the Supplement), so we considered differences between approaches (MAP-GT, MLE-GT, MLE-RI) for branch lengths less than 4.25 CUs. When the the true branch lengths were longer than 1 CU and shorter than 4.25 CUs, the MLE-RI branch lengths had 1–3% error on average; in contrast, both the MAP-GT and MLE-GT branch lengths had over 30% error on average and were upwardly biased for all 25 replicates. As the true branch lengths decreased below 1 CU, the error in the MAP-GT and MLE-GT branch lengths decreased. There was little difference between the MAP (or MLE) branch lengths derived for gene trees compared to the MLE branch lengths derived for RIs when the true branch length was shorter than 0.3 CUs (Fig. 4A–C; Table S1 in Supplement).

**Figure 4:**
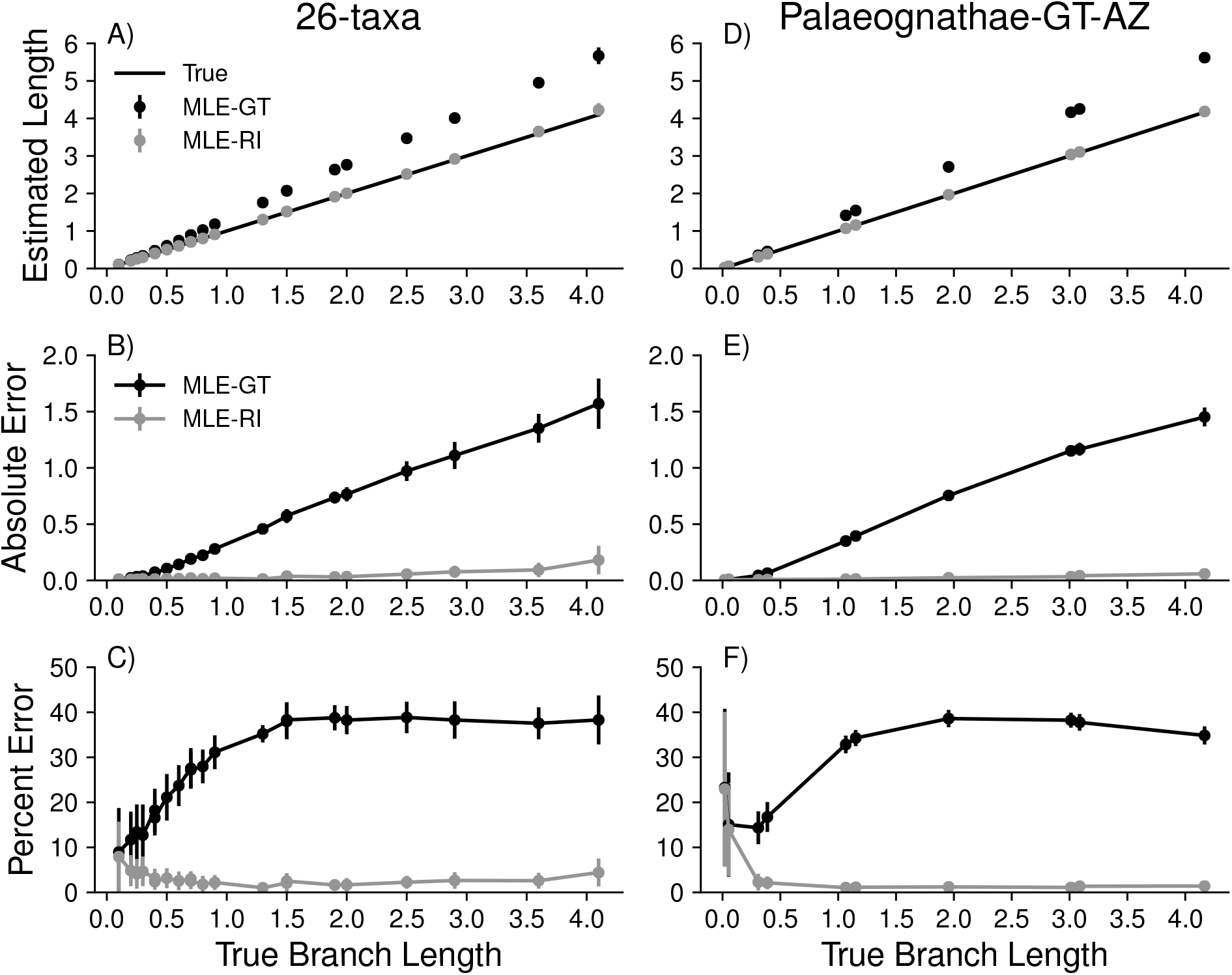
Branch length estimation using ASTRAL BP for two of the four simulated data sets. Results for 26-taxa and Palaeognathae-GT-AZ model trees are shown in subfigures A–C and subfigures D–F, respectively. Subfigures (A) and (D) show the true species tree branch lengths (*x*-axis) plotted against either the true branch lengths, the lengths computed using the MLE derived for gene trees, or lengths computed using the MLE derived for RIs (Equation 8). Subfigures (B) and (E) show the absolute error: *abs*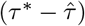, where 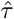 is the estimated branch length and *τ* * is the true branch length. When the true branch length *τ* * is greater than 0.25 CUs, the branch length estimates computed using the MAP-GT and MLE-GT equations are greater than the true branch length for all 25 replicates (Table S1 in the Supplement). Subfigures (C) and (F) show percent error: 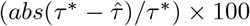. All values are averaged over 25 replicate data sets; dots are means, and bars are standard deviations. Note that only true branch lengths of less than 5 coalescent units are shown, as branch length estimation is ill-conditioned for longer branch lengths (Table S1 in Supplement).

#### Results for varying numbers of RIs

As the number of parsimony-informative RIs increased from 100 to 100 000, species tree error decreased for the six methods studied, converging to the results described above for 100 000 RIs. Namely, five methods (ASTRAL BP, ASTRID BP, Dollo parsimony, MDC BP, and unordered parsimony) converged to the true species tree topology, achieving a mean false negative error rate of zero when given 100 000 RIs (Fig. 5A). In contrast, Camin-Sokal parsimony converged to a tree with one incorrect branch for all but one of the 25 replicates with 100 000 RIs, achieving a mean false negative error rate 0.0417.

**Figure 5:**
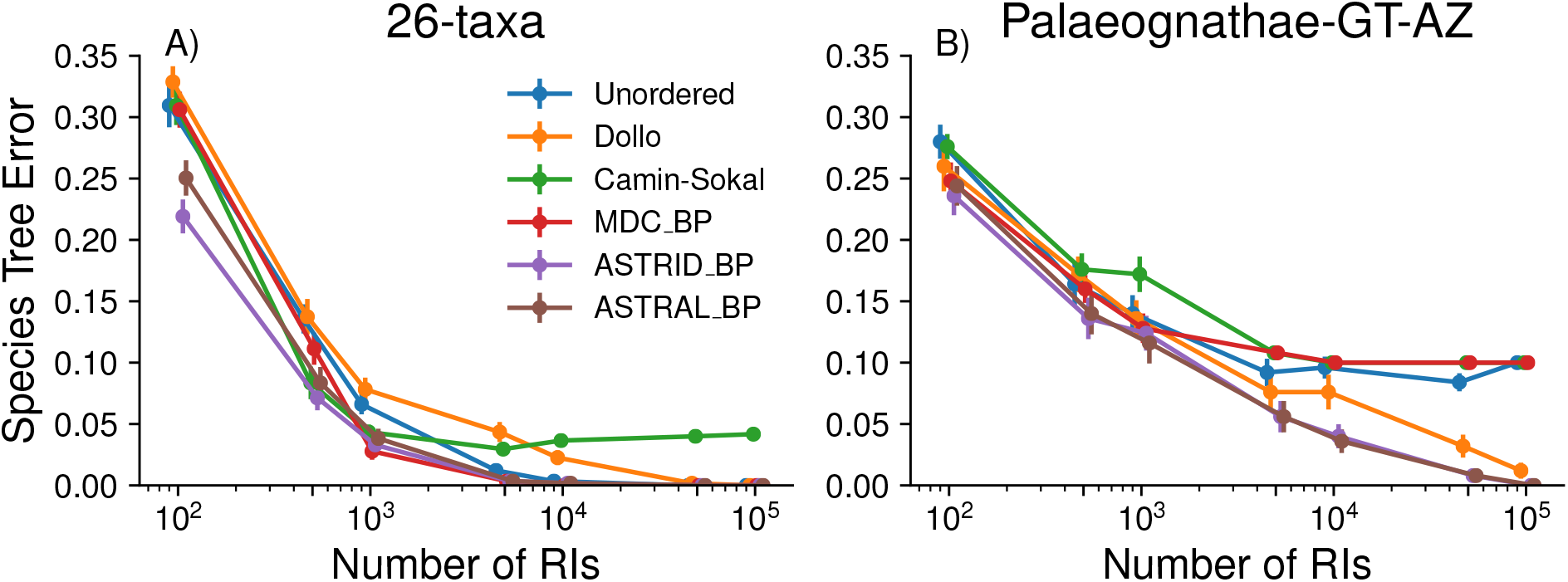
Impact of varying the number of retroelement insertions (RIs) on species tree estimation methods (indicated by color in the legend). The results for the 26-taxa and Palaeognathae-GT-AZ data sets are shown on the left and right, respectively. Species tree error (*y* − *axis*) is measured as the false negative rate (i.e. the number of internal branches in the true species tree that are missing from the estimated tree, divided by the number of internal branches in the true species tree). These values are shown across 25 replicate data sets; dots are means, and bars are standard errors (note: the dots are jittered). The number of parsimony-informative RIs (*x*-axis) in these simulated data sets varies from 100 to 100 000.

Examining five branch lengths with varying lengths (0.1, 0.2, 0.5, 1.3, and 2 CUs) in the true species tree, we found that whether or not a branch was recovered in the ASTRAL BP tree depended on its length in the model species tree. For example, on data sets with 1 000 parsimony-informative RIs, ASTRAL BP recovered the branches with lengths 0.1 and 0.2 CUs in 56% and 84% of replicates, respectively; in contrast, branches with lengths 0.5, 0.13, and 2 CUs were recovered in 100% of replicates (Fig. 6A). Overall, the fraction of replicates in which branches were recovered increased with the number of parsimony-informative RIs, but this increase was faster for longer branches. Similar to the results for topological accuracy, the accuracy of branch length estimation (MLE-RI) improved with increasing numbers of RIs, and the decrease in percent error was faster for longer branch lengths (Fig. 6B).

**Figure 6:**
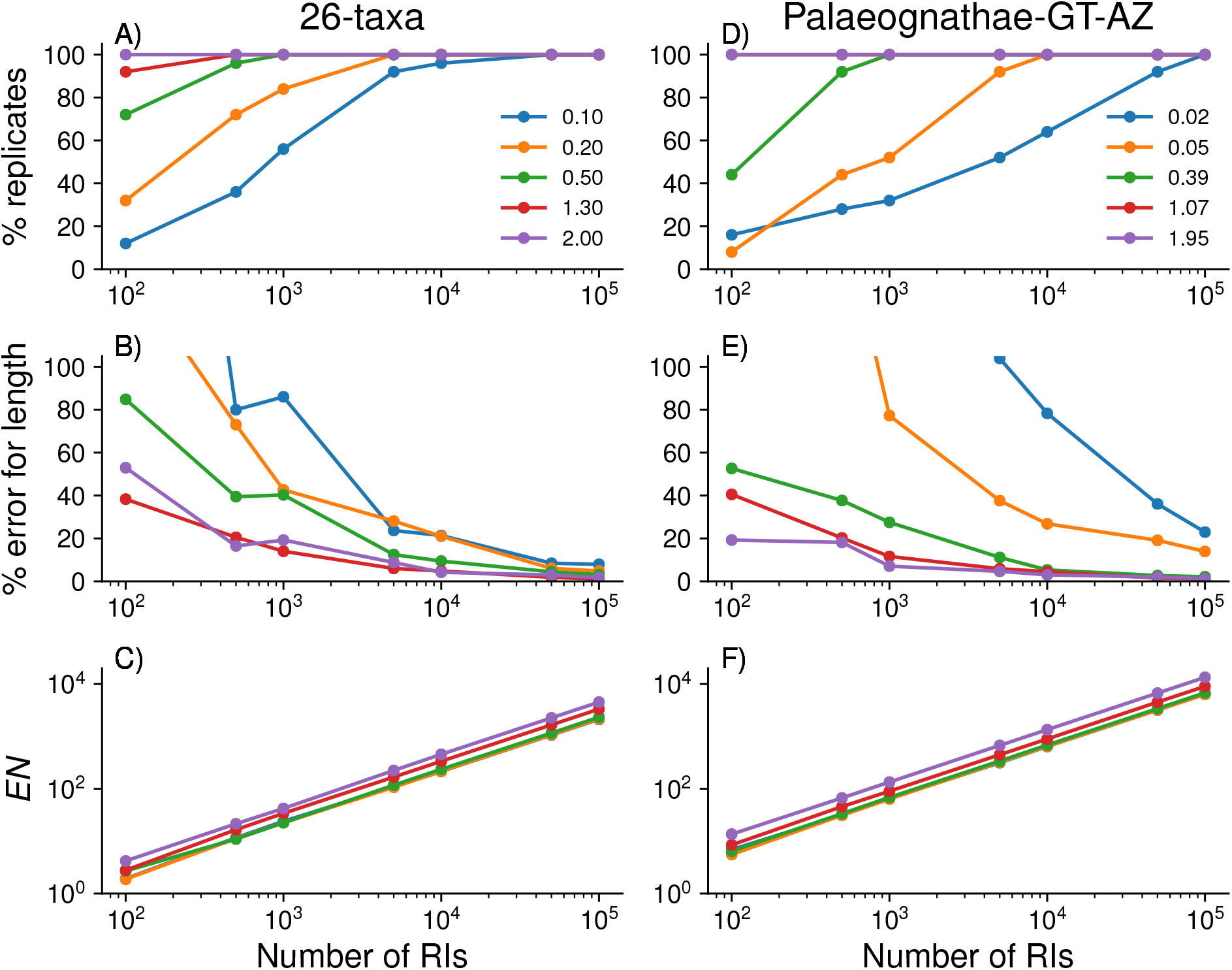
Impact of varying the number of retroelement insertions (RIs) for quantities estimated for specific branches in the model species tree. Subplots A–C shows results for the 26-taxon-Sim data set, and subplots D–F shows results for the Palaeognathae-GT-AZ data set. Subplots A and D shows the effective number (*EN*) around the branch (i.e. the approximate number of RIs containing information for resolving the branch). Subplots B and E shows the percent of replicates in which the branch was recovered by AS-TRAL BP. Subplots C and F show the percent error for the estimated branch length (MLE-RI). All values are shown averaged over replicates for which the branches (indicated by its length in the true species tree) were recovered by ASTRAL BP. The number of parsimony-informative RIs (*x*-axis) in these simulated data sets varies from 100 to 100 000.

It is worth noting that the number of parsimony-informative RIs in the input data set is not equivalent to the number of RIs that are informative for resolving a specific branch; we refer to the latter as the effective number (*EN*) of RIs. ASTRAL BP estimates this value for all internal branches, and we found that *EN* depends on the branch lengths in the model species tree (Fig. 6C). For simulated data sets with 1 000, 10 000, and 100 000 parsimony-informative RIs, the branch with length 0.2 CUs had mean *EN* of 23, 216, and 2 161, respectively; in contrast, the branch with length 2.0 CUs had mean *EN* of 42, 455, and 4 510 for those same data set sizes. Put simply, the branch with 2.0 CUs had ∼ 2x more data than the branch with length 0.2 CUs.

Lastly, we evaluated the utility of ASTRAL BP’s support values (i.e. local PP) for controlling false positive branches. When there were at least 1 000 RIs, collapsing branches with local PP less than 0.8, resulted in the mean false positive rate of 0.004. In this scenario, the mean false negative rate was 0.12 (∼ 3 branches) on average. As the number of RIs increased, both error rates decreased (Fig. S1A–B in Supplement).

### 3.4 Palaeognathae-GT-AZ

#### Results for 100 000 RIs

Only three methods (ASTRAL BP, ASTRID BP, and SDPquartets) recovered the correct species tree for all 25 replicate data sets (Fig. 3C) with 100 000 RIs simulated from the Palaeognathae-AZ model tree. The model species tree has the pectinate topology: ((((*emu, cassowary*), *kiwi*), *rhea*), *tinamou*). Dollo parsimony recovered the correct tree for 22 of 25 replicate data sets and in the other three instances reconstructed an incorrect tree that swapped the position of rhea and tinamou. The other four methods (Camin-Sokal parsimony, MDC BP, polymorphism parsimony, unordered parsimony) always recovered an incorrect species tree that placed rhea as the sibling of tinamou (Fig. 3C). This incorrect topology has greater probability than the model species tree topology, when considering the probabilities of rooted gene trees under the MSC (see Methods for details).

The trends for branch length estimation with ASTRAL BP were similar to the 26-taxa model species tree (Figure 4D–F; Table S1 in Supplement). When the true branch lengths were shorter than 0.3 CUs, there was little difference between the three branch length estimation techniques (MAP-GT, MLE-GT, and MLE-RI). When the true branch lengths were between 0.3 and 1 CUs, the MLE-RI estimate had 2% error on average, whereas the MAP-GT and MLE-GT estimates were upwardly biased for all 25 replicates, with 14–17% error on average. When the true branch lengths increased (i.e. they were longer than 1 CU), the percent error in the MAP-GT and MLE-GT estimates increased to over 30% on average; in contrast, the percent error in the MLE-RI estimate decreased to 1% on average.

#### Results for varying numbers of RIs

The results for varying numbers of RIs were consistent with the trends observed for the 26-taxa model species tree (Figs. 5B; 6D–F; S1C–D in Supplement).

## 4 ASTRAL BP Analysis for Palaeognathae

As previously discussed, the Palaeognathae-GT-AZ model species tree was based on the tree estimated by Cloutier et al. (2019), who ran ASTRAL on 29 850 estimated gene trees. The two shortest branches in the Palaeognathae-GT-AZ tree are consecutive: branch *x* separates emu, cassowary, and kiwi from the remainder of taxa, and branch *y* separates emu, cassowary, kiwi, and rhea from the remainder of taxa. Branches *x* and *y* have (MLE-GT) lengths of 0.02 and 0.05 CUs, respectively (Fig. 1C), putting the Palaeognathae-GT-AZ tree in the AZ (see Methods section for details). This is in contrast to the tree estimated by Springer et al. (2020), who ran ASTRAL BP on a data set assembled by Cloutier et al. (2019) containing 4 301 parsimony-informative RIs. This tree (referred to as Palaeognathae-RI) has the same topology as the Palaeognathae-GT-AZ tree but is *not* in the anomaly zone (Figure S2 in the Supplement). Specifically, branches *x* and *y* have (MLE-RI) lengths 0.87 and 0.26 CUs, respectively (note: Springer et al., 2020 used the MAP-GT formula to compute branch lengths, which in part motivated our study on branch length estimation and the derivation of the MLE-RI formula).

Because of the differences between Palaeognathae-GT-AZ and Palaeognathae-RI species trees, we explored the effectiveness of ASTRAL BP on data sets simulated from Palaeognathae-GT-AZ model tree with just 1 000 or 5 000 parsimony-informative RIs. The correct species tree topology was recovered in 7/25 and 13/25 replicates for data sets with 1,000 and 5,000 RIs, repectively (Figs. S3 and S8 in the Supplement). For all data sets, false positive branches (i.e. branches in the estimated species trees that are not in the model species trees) determined the branching order of four clades: emu+cassowary, kiwi, rhea, and tinamou. The resolution of these four clades results in either a *pectinate* or a *symmetric* topology. A pectinate tree was returned by ASTRAL BP in 20/25 replicates with 1,000 RIs; at least 17 of these trees were in the anomaly zone. For data sets with 5,000 RIs, a pectinate tree was returned by ASTRAL BP in 17/25 replicates; all 17 of these trees were in the anomaly zone.

Regardless of the tree topology returned, the two internal branches separating these four clades were quite short (Figs. S4–S7 and S9–S17 in the Supplement). For any incorrect branch, the average (*±* standard deviation) length was 0.06 *±* 0.03 and 0.03 *±* 0.02 CUs for data sets with 1 000 and 5 000 RIs, respectively. For the correct branches (*x* and *y*), the mean branch lengths were 0.09 *±* 0.05 and 0.07 *±* 0.05 CUs, respectively, for data sets with 1 000 RIs. For data sets with 5 000 RIs, branch *x* had mean length 0.06 *±* 0.02 CUs, and branch *y* had mean length 0.04 *±* 0.02 CUs.

Importantly, the two shortest internal branches had lower support values (local PP) in all estimated species trees, indicating uncertainty regarding the specific branching order. For any incorrect internal branch, the average (*±* standard deviation) local PP of any incorrect branch was 0.52 *±* 0.10 (range: 0.38 to 0.71) and 0.56 *±* 0.13 (range: 0.37 to 0.83) for data sets with 1,000 and 5,000 RIs, respectively. In contrast, for both data sets sizes, the other eight internal branches were recovered in 25/25 replicates with high support (Tables S4 in Supplement).

These analyses suggest that ASTRAL BP should be useful for identifying major clades (i.e clades separated by branches *>* 0.3 CUs) and rapid radiations (i.e. shorter connected branches), even when data sets have only a few thousand RIs, as was the case for the Palaeognathae RI data set analyzed by Springer et al., 2020. In the next section, we discuss some potential explanations for differences between the Palaeognathae-GT-AZ tree and the Palaeognathae-RI tree.

## 5 Discussion

In our simulation study, we evaluated eight methods for estimating species trees from RIs on four model species trees, at least three of which are in the AZ. Both the 4-ingroup-taxa-AZ and 5-ingroup-taxa-AZ model species trees are pectinate, with two consecutive short branches with lengths of 0.01 CUs. Patel et al. (2013) suggested 400 000 years of evolution along a branch is a reasonable approximation for a coalescent unit in vertebrates. If we use this approximation, then branch lengths of 0.01 CUs are equivalent to just 4,000 years and highlight the challenging conditions that we evaluated methods on.

For model species trees in the AZ, the two quartet-based methods (ASTRAL BP and SDPquartets) and the distance-based method (ASTRID BP) always recovered the correct species tree topology when given 100 000 RIs. Dollo parsimony also performed well, only failing to recover the correct species tree topology in 3 of 25 replicates simulated from the Palaeognathae-GT-AZ model species tree. MDC BP, polymorphism parsimony, and unordered parsimony generally performed well for the 26-taxa model condition (which is unlikely to be in the AZ based on our analyses) as well as the simplest anomaly zone situation with four ingroup taxa (Fig. 1A).

The 4-ingroup-taxa-AZ model species tree has been considered in previous studies, including those by Mendes and Hahn (2017) and Than and Rosenberg (2011). Our results for unordered parsimony are consistent with the work of Mendes and Hahn (2017), who also found that unordered parsimony recovered the correct species tree topology on data simulated from this same model species tree, although they simulated mutations down each gene tree under the Jukes-Cantor model rather than the infinite-sites model. These experimental results highlight the point made by Mendes and Hahn (2017) that a model species being in the AZ does not guarantee that concatenation-based approaches (e.g. unordered parsimony) will perform poorly. Indeed, Mendes and Hahn (2017) hypothesized that unordered parsimony should return the correct species tree for this model tree, even though the most probable gene tree differs from the species tree, because the anomalous gene trees have very short internal branches, on average, relative to internal branches on gene trees that agree with the species tree. Put simply, the net effect of these branch length differences is that the most common (democratic) site patterns will still support the correct species tree.

Our results for MDC BP differ from work by Than and Rosenberg (2011), who showed that for a pectinate species tree with four ingroup taxa, the MDC criterion is statistically inconsistent if branch *x* = *y <* 0.2215 CUs. This theoretical result assumes rooted, binary gene trees as input, rather than RIs encoded as unrooted trees with a single bipartition; therefore, it is not directly applicable to our simulation study. It is reasonable that MDC would achieve good performance on data sets simulated from the four-ingroup-taxa-AZ model species tree when we consider how the MDC method, implemented within PhyloNet, operates when given a set of unrooted and unresolved gene trees 𝒢= *g*_1_, *g*_2_, … *g*_*k*_. In this scenario, the solution to the MDC problem is a rooted, binary species tree that minimizes the number of deep coalescences across all possible sets 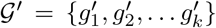 of rooted, binary gene trees with the property that 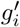 can be obtained from *g*_*i*_ by refining polytomies and adding a root (Yu et al., 2011). PhyloNet implements a dynamic programming algorithm (Yu et al., 2011) that finds the optimal species tree within a constrained version of tree space—the space is constrained using bipartitions in the set 𝒢, in a fashion similar to ASTRAL (Mirarab et al., 2014). For the data sets of 100 000 RIs simulated from the four-ingroup-taxa-AZ model species tree, our unordered parsimony analysis suggests that the most common site pattern (or bipartition) in the input set RIs agrees with the model species tree. In this case, all bipartitions in the model species tree topology exist in the constrained search space, and the majority of “gene trees” are bipartitions that agree with the model species tree and thus can be rooted/resolved in such a way that there are no deep coalescences.

In more complex AZ situations with greater than four ingroup taxa (i.e. the 5-ingroup-taxa-AZ and Palaeognathae-Sim model species trees), MDC BP, polymorphism parsimony, and unordered parsimony all failed—even when given 100 000 RIs as input. Camin-Sokal also failed to recover the correct topology in these cases, as well as for two easier model conditions (i.e. 4-ingroup-taxa-AZ and 26-taxa).

Overall, our results suggest that Camin-Sokal parsimony, polymorphism parsimony, and unordered parsimony are inappropriate methods for estimating species trees from RIs when the species tree is in the AZ and has more than four ingroup taxa. Dollo parsimony performs much better than the other parsimony methods in the AZ situations examined here, although it was not as accurate (or as computationally efficient) as ASTRAL BP or ASTRID BP. These results are significant because RI data sets are commonly analyzed using variants of parsimony, including Camin-Sokal (e.g. Nikaido et al., 1999; Nilsson et al., 2010; Suh et al., 2011), unordered (e.g. Gatesy et al., 2013, 2019), polymorphism (e.g. Suh et al., 2015b; Doronina et al., 2015), and Dollo (e.g. Lammers et al., 2019).

Given our theoretical results pertaining to statistical consistency (of ASTRAL BP and SDPquartets), we suggest that ASTRAL BP and SDPquartets are the most appropriate of the tested methods for inferring species trees with RIs. ASTRAL BP may be particularly useful to researchers, as it automatically estimates branch lengths (in CUs) and branch support (local PP). We derived the MLE for RIs (rather than for gene trees) and found that using this corrected formula improved the accuracy of branch length estimation in our simulation study, expect when the branch length was short enough so that all approaches (MAP-GT, MLE-GT, and MLE-RI) produced similar results, as predicted by the small angle approximation. Our simulation study also revealed that the branch support (local PP) can be used to control false positive branches successfully. We expect that ASTRAL BP should also perform well with other low-homoplasy absence/presence characters such as NUMTs and large indels that, along with RIs, are becoming increasingly easy to mine from genomic sequences (Schull et al., 2019; Churakov et al., 2020).

A major challenge to estimating species trees from RIs is the amount of data available. Unlike binary gene trees, which display a quartet topology on *all* subsets of four species, site patterns display a quartet topology on only *some* of the subsets of four species. In our simulation study, the effective number (*EN*) of RIs around short branches was lower compared to long branches, and short branches may require larger *EN* to be accurately estimated, as the probabilities of the dominant and alternative quartets will be closer. When *EN* is too low for a given branch, the quartet frequencies may fail to provide good estimates of the quartet probabilities under the MSC+infinite sites model, impacting our ability to accurately estimate the branch length and local PP.

In our analysis of the Palaeognathae-RI data set (4 301 parsimony-informative RIs), *EN* was always less than 30 for the two shortest internal branches (Table S3 in Supplement) and thus their lengths and support values should be interpreted with caution. Low *EN* for the Palaeognathae-RI tree is one explanation for the differences in branch lengths compared to the Palaeognathae-GT-AZ tree. However, in our simulation study, ASTRAL BP was effective for identifying potential rapid radiations, even when data sets had just 1 000 parsimony informative RIs simulated from Palaeognathae-GT-AZ model species tree. Notably, the *EN* for the two shortest internal branches was greater in the simulated data sets with 1 000 RIs than in the biological data set with 4 301 RIs (Table S3–S4 in the Supplement). These differences in *EN* could be due to a variety of factors, including model violations (e.g. due to the number of new RIs may not be constant across the species tree due to boom-bust activity, biases introduced when assembling RI data sets, inaccurate coding of RIs, missing data, etc.); these issues should be explored further in future work.

It is also worth noting that simulation studies have shown that high gene tree reconstruction error can result in branch lengths in the estimated ASTRAL species trees that are too short by almost an order of magnitude (Sayyari and Mirarab, 2016; Simmons and Gatesy, 2021). Gene tree reconstruction error is prevalent among phylogenomic studies and can occur because of long-branch attraction, missing data, model misspecification, homology errors, arbitrary resolution of polytomies by programs such as RAxML, and other causes (Gatesy and Springer, 2014; Springer and Gatesy, 2014, 2016).

To summarize, the Palaeognathae-GT-AZ tree is in the AZ (but the branch length estimates could be impacted by gene tree estimation error, etc.) and the Palaeognathae-RI tree is *not* in the AZ (but the branch length estimates could be impacted by low *EN*). In the future, it would be interesting to use simulations to compare species tree estimation (including branch lengths) from gene trees versus RIs. These simulations should be performed at various phylogenetic depths and with short internal branch lengths. Each gene tree has the potential to contain more phylogenetic information than its respective RI (i.e. the RI simulated down the gene tree). However, RIs could still fare well in such simulations, especially at deep divergences where the estimation of gene trees can be challenging. Unlike DNA sequences, which show increased homoplasy with depth, RIs are low-homoplasy markers in both shallow and deep phylogenetic settings when accurately coded. Although RIs become more difficult to characterize at deep divergences because indels and other mutations can erase or obscure their history, we conjecture that they will be useful for phylogenetic problems that are at least as old as the radiations of placental mammals, crocodylians, and birds that each extend to the Cretaceous (Nishihara et al., 2009; Haddrath and Baker, 2012; Suh et al., 2015a,b; Doronina et al., 2017).

Published data sets for mammalian RIs range from those with *<*100 retroelements (e.g., placental root, [Nishihara et al., 2009]) to 91 859 for eight species of baleen whales (Lammers et al., 2019). In the latter case, 24 598 of these insertions are phylogenetically-informative and occur in two to six of the balaenopteroid species. For protein-coding genes, the number of available loci is relatively fixed whether a data set includes genomes from five mammalian species or 500, because the majority of protein-coding genes are shared among these taxa. In humans, a recent estimate for the total number of protein-coding genes is 19 116 (Piovesan et al., 2019). By contrast, RIs are segregating sites as are single nucleotide mutations, albeit without the attendant homoplasy in the latter, and retroelement data sets are expected to increase in size as more taxa are added to a data set. For a taxonomically diverse genomic data set with more than 200 mammal species (e.g., Genereux et al., 2020), we are optimistic that it will soon be possible to extract hundreds of thousands or even millions of informative retroelements as improved methods become available for efficiently extracting and applying quality-control filtering steps to assemble these data sets (Churakov et al., 2020). Indeed, the number of informative markers will grow even larger if such data sets also include NUMTs and large indels (Schull et al., 2019). Lastly, combining these low-homoplasy markers with sequence-based gene trees is a valuable direction of future research for mitigating the impact of gene tree estimation error and maximizing the amount of high quality data available for species tree estimation.

## 6 Conclusions

Because accurately-coded RIs are unlikely to be impacted by homoplasy, we performed an experimental study in which RIs were simulated under the MSC+infinite-sites model, with the expected number of new RIs per generation held constant across the species tree. In this setting, two recently proposed methods, ASTRAL BP and SDPquartets, outperformed methods that have been commonly used to analyze RIs, especially when the model species tree was in the anomaly zone and had more than four ingroup taxa. Furthermore, we showed that ASTRAL BP and SDPquartets have desirable theoretical properties, that is, they are statistically consistent under the model conditions described above. We recommend that researchers apply ASTRAL BP to empirical data sets with well-vetted coding of RIs (Doronina et al., 2019) and examine the *EN* for each branch of trees estimated using ASTRAL BP to determine whether the RI data set contains sufficient phylogenetic information to resolve branches of interest. Future work should evaluate these methods in the context of model violations that occur when RIs are not perfectly encoded, the sampling of RIs is biased, the rate of new RIs is not constant across the species tree (e.g. boom-bust activity), or there is missing data. We are optimistic that, with the continued development of quartet-based methods and the accurate assembly of ultra-large RI data sets, it may be possible to resolve some of the most challenging questions in systematics.

## 8 Acknowledgements

We thank Rick Baker for technical advice, Siavash Mirarab and Arun Durvasula for discussions, and Matthew Hahn and two anonymous reviewers for detailed feedback that improved the quality of this work.

## 10 Supplementary Material

Data available on Github: https://github.com/ekmolloy/retrosim-study

## 11 Appendix

### 11.1 Statistical Consistency

We begin with the proof of statistical consistency for ASTRAL_BP.

#### Theorem 2.

Suppose that RIs are generated under the MSC+infinite-sites model (as approximated by Doronina et al., 2017) and that the expected number of new RIs per generation is constant across the species tree. Then, ASTRAL_BP is statistically consistent estimator of the unrooted species tree topology.

**Proof Sketch.** Let *T* * be the true species tree on taxon set *S*, and let *ℛ* be a set of retroelement insertions (RIs), also on taxon set *S*. Each RI *r* ∈ *ℛ* corresponds to a set of quartet topologies; this set can be created by restricting *r* to each subset *S*_*i*_ of four taxa, denoted *r*|_*S*_, for all 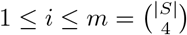. Similarly, given some binary tree *T*, also on taxon set *S*, we can restrict *T* to taxon set *S*_*i*_, denoted 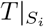, and compare the resulting topology to 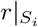. There are three possible scenarios either 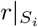 corresponds to the same quartet topology 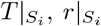, corresponds to a different quartet topology than 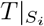, or 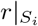does not correspond to any of the three possible quartet topologies on *S*_*i*_. For example, the site pattern 11000 for taxon set *{A, B, C, D, E}*, which can be represented as bipartition *A, B*|*C, D, E* or newick string ((*A, B*), *C, D, E*), does not display a quartet topology when it is restricted to taxon subset *{A, C, D, E}*.

Let 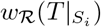 denote the number of RIs in *ℛ* that display the same quartet topology as 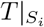, and let *n*_*i*_ be the number of RIs in *ℛ* that displays any of the three possible quartet topologies on taxon subset *S*_*i*_. For all *i* ∈ 1, 2, …, *m*, as the total number of RIs goes to infinity, *n*_*i*_ also goes to infinity (i.e. *n*_*i*_ is not bounded). Then, by Theorem 1 in the Supplemental Text (which says the most probable quartet agrees with the true species tree *T* *), for any possible tree topology *T* on taxon set *S* and for all 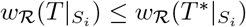 with probability going to one, as the number of RIs goes to infinity. It follows that the true species tree *T* * is the unique optimal solution to maximum quartet support supertree (MQSS) problem with high probability (recall that the MQSS problem is to find *T* such that 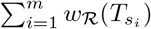 is maximized).

The MQSS problem is NP-hard (Jiang et al., 2001; Lafond and Scornavacca, 2019); however, when the solution space is constrained by a set Σ of bipartitions, it can be solved in polynomial time (Bryant and Steel, 2001; Mirarab et al., 2014). ASTRAL implements an exact algorithm for solving the bipartition-constrained version of the MQSS problem, and by default, every bipartition in the input (in this case) is added to the constraint set Σ. Because every RI (0/1) pattern occurs with non-zero probability under the MSC+infinite-sites model, the probability that every bipartition in *T* * is represented by a RI in *ℛ* goes to 1, as the number of RIs goes to infinity; therefore, ASTRAL given *ℛ* returns the true species tree with high probability.

### 11.2 Maximum Likelihood Branch Length Estimation

We now show how to compute the maximum likelihood (ML) estimate of the lengths of the internal branches for RI data sets using the framework (and notation) proposed by Sayyari and Mirarab (2016).

Consider a species tree on four taxa *{A, B, C, D}* with unrooted topology: *A, B*|*C, D*. Let *θ*_1_ denote the probability that a parsimony-informative RI corresponds to the dominant topology *t*_1_ = *A, B*|*C, D*, and let *θ*_2_ and *θ*_3_ denote the probabilities that a parsimony-informative RI corresponds to the two alternative topologies *t*_2_ = *A, C*|*B, D* and *t*_3_ = *A, D*|*B, C*, respectively. Then, *θ*_1_, which is *P*_*RI*_ (*A, B*|*C, D*) in Equation 6, is a function of the true internal branch length *τ* *, and both *θ*_2_ and *θ*_3_ are functions of *θ*_1_. Therefore, we can rewrite 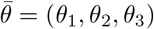 as 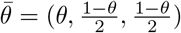, respectively.

Suppose that the RIs are generated under the MSC+infinite-sites model (as approximated by Doronina et al., 2017) and that the expected number of new RIs per generation was constant across the species tree. For RIs generated under this model, we can count the number of RIs that display one of the three quartet topologies *t*_1_, *t*_2_, and *t*_3_, denoting these frequencies *n*_1_, *n*_2_, and *n*_3_, respectively. For a given model species tree, each RIs is independent under the MSC+infinite-sites model. Therefore, we can model the RI frequencies 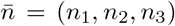 as a multinomial random variable 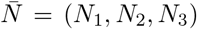 with event probabilities 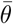 and 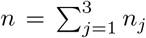 trials, as noted by Sayyari and Mirarab (2016). Alternatively, we could define some hidden random variable 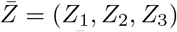 in the same way (i.e. 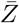 is the “true” quartet frequencies) and then model 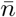 as 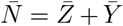, where 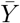 is some noise term with mean zero. Following Sayyari and Mirarab (2016), we treat the expected value of 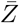 as an observed value and estimate it, denoted 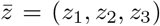, from our empirical RI data set. We discuss this procedure further at the end of the section, but for now, we simply set 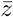 equal to 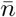 and ignore noise.

#### Theorem 3.

For a four-taxon model species tree, suppose that RIs are generated under the MSC+infinite-sites model (as approximated by Doronina et al., 2017). For some quartet topology *q*, let 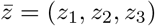 be the quartet frequencies with *z*_1_ corresponding to *q*. Given that *q* corresponds to the true unrooted species tree topology and given the modeling of 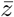 described above, the ML estimate of its internal branch length is 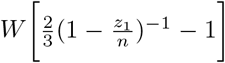, where *W* is Lambert’s function and *n* = *z*_1_ + *z*_2_ + *z*_3_ is the number of trials (i.e. parsimony-informative RIs). This holds for ^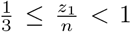^, When 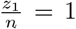, the ML estimate does not exist (note: we set the length equal to ∞ in this case), and when 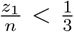, the quartet *q* does not correspond to the true unrooted species tree topology (note: we set the branch length equal to 0 in this case).

*Proof*. Let *D* ∈ [0, ∞) be a branch length. We model *D* as a random variable, so the ML estimate of the branch length is arg 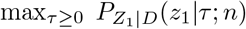, where 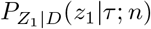 is the likelihood of *D*. Given that quartet *q* agrees with the true unrooted species tree topology, by Lemma 1 in Sayyari and Mirarab (2016) and Equation 6,

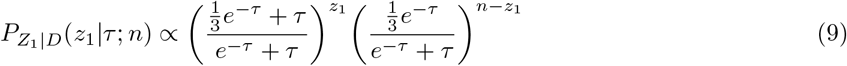

We now compute the log-likelihood function

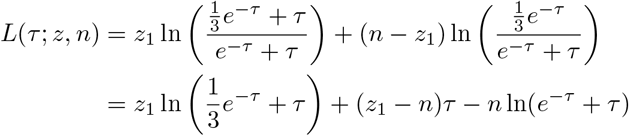

dropping the constant terms. To find the critical point, we take the first derivative of the log likelihood function

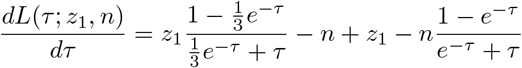

and set it equal to 0. Therefore, the critical point is given by

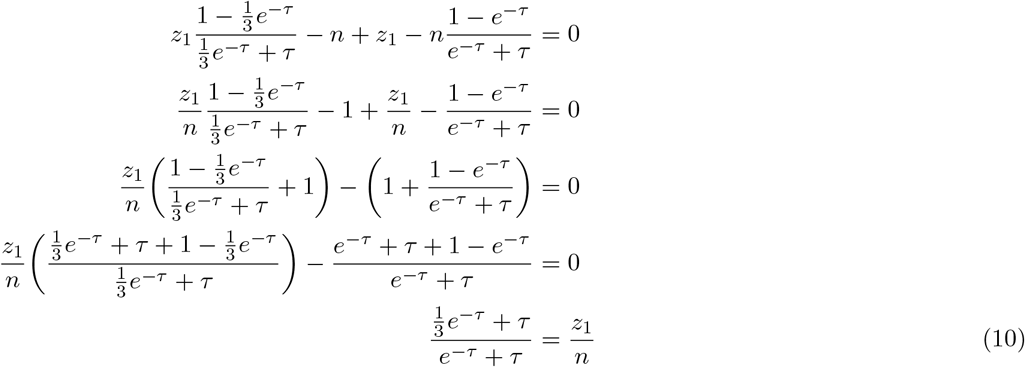

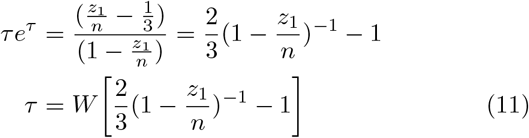

where *W* is the Lambert’s function. Note that 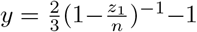 is positive when ^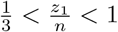^ and is undefined at 1. By Proposition 5 in Borwein and Lindstrom (2016), *W* [*y*] is positive for *y* ∈ [0, ∞) and concave on (*-*1*/e*, ∞) (also see https://www.carma.newcastle.edu.au/resources/jon/WinOpt.pdf). Therefore, the critical point is a local maximum of the likelihood function of *D* when 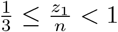. When 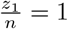 (i.e. there is no conflict), the maximum likelihood estimate does not exist; in this case, we set the estimated length to ∞ by default. Similarly, we set the estimated length to 0 when 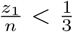, as this implies the internal branch is not in the species tree.

When there are more than four taxa in the species tree, the situation is more complicated. Consider a branch *Q* that splits two rooted subtrees *T*_*A*_ and *T*_*B*_ from two other rooted subtrees *T*_*C*_ and *T*_*D*_. Now let *𝒜, ℬ, 𝒞*, and *𝒟* be the taxon (leaf) sets of *T*_*A*_, *T*_*B*_, *T*_*C*_, and *T*_*D*_, respectively. This implies that *Q* induces *m ′* = | *𝒜*| *×* | *ℬ*| *×* |*𝒞*| *×* | *𝒟*| quartets, so we say that there are *m* quartets “around” *Q*. As noted by Sayyari and Mirarab (2016), we can consider the frequencies of these *m* quartets and use them to estimate 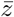.

For *k* ∈ {1, 2, …, *m*}, let 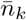 denote the frequency vector for quartet *k*, that is, the first element of 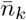, denoted *n*_1*k*_, corresponds to frequency of quartet *k* and let the second two elements of 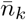, denoted *n*_2*k*_ and *n*_3*k*_, correspond to the frequencies of the two alternative topologies. One possibility is to take just one of the *m′* quartets around branch *Q* and compute the MLE of the branch length, setting *z*_1_ = *n*_1*k*_ and *n* = *n*_1*k*_ + *n*_2*k*_ + *n*_3*k*_ (where *k* is the index of the selected quartet). We could also select the quartet *k* so that the number *n* of trials is maximized. Alternatively, because 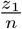 is the frequentist estimate of the probability of the dominant quartet (Equation 10), we could estimate it using the *m′* quartets as follows.

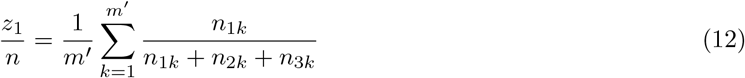

Sayyari and Mirarab (2016) provide an efficient algorithm for approximating this quantity, referred to as quartet support (“q1” when running ASTRAL with option “-t 2”). In our simulation study, we found that plugging “q1” into Equation 11 for 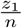 produced accurate results (Figure 4).

### 11.3 Local Posterior Probability

Sayyari and Mirarab (2016) also show how to compute the local posterior probability (PP) for branch *Q* (Therem 1 in Sayyari and Mirarab, 2016). To put a prior on *θ*, they assume that the gene trees are generated under the MSC from a model species tree generated under the Yule process with birth rate *λ*. Under this assumption, branch lengths in the species tree are exponentially distributed, so *f*_*D*_(*τ*) = 2*λe*^2*λτ*^ can be used as the prior for branch lengths (Stadler and Steel, 2012). By default, ASTRAL sets 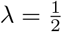, which corresponds to a uniform prior on *θ* (Lemma 1 in Sayyari and Mirarab, 2016).

We show that for RIs the prior on *θ*, denoted *f*_*θ*_, is not uniform when the species tree is generated under a Yule process with birth rate 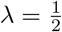. For 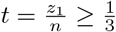 ,

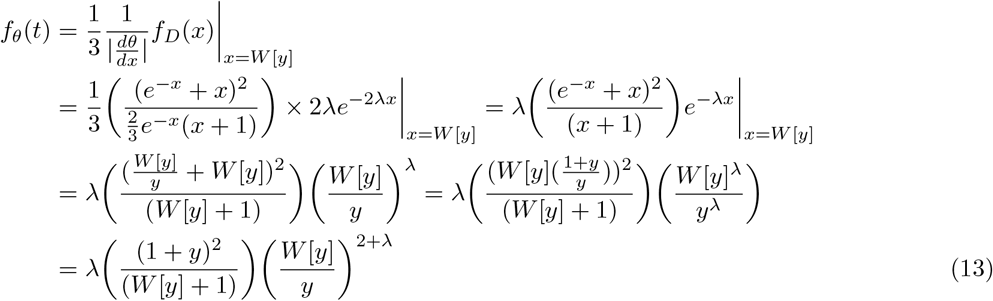

where 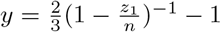. Recall that 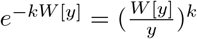 (shown as part of Proposition 6 in Borwein and Lindstrom, 2016) and Lambert’s W is positive on [0, ∞) (Proposition 5 in Borwein and Lindstrom, 2016). The prior on *θ* is not uniform on 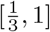 when 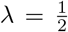 (Figure S18 in Supplement); furthermore, this prior makes it difficult to integrate

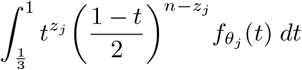

when computing local PP.

It is reasonable to have a uniform prior on *θ* in the absence of other information; therefore, one could justify utilizing the branch support returned by ASTRAL for RI data sets for four species, although we cannot keep the interpretation that the prior on branch lengths comes from species trees generated under the Yule process with birth rate *λ*. In our simulation study, the local PP computed by ASTRAL was effective at controlling the FP rate (Supplementary Figure 1). Note that when the number of species is greater than four, as was the case in our simulation study, ASTRAL uses the *m′* quartets to estimate 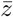 and the number of trials *n*. The latter quantity is the effective number (*EN*) of gene trees or RIs for the branch (this is “EN” when running ASTRAL with option “-t 2”). When *EN* is low, estimates of the branch length and local PP should be interpreted cautiously; this can be an issue when analyzing RI data sets.

