## Supplement for "Theoretical and practical considerations when using retroelement insertions to estimate species trees in the anomaly zone"

August 20, 2021

**3 Supplementary Results** **14**

**4 Comparing of Branch Length Estimates** **14**

**5 Conditioning of Branch Length Estimation** 15

**6 Using local PP to control false positive branches** 16

**7 ASTRAL_BP Analyses for Palaeognathae** 17

**8 Non-Uniform Prior for Local Posterior Probability** 34

**References** 35

### 1 Supplementary Methods

#### 1.1 Commands for testing whether model species trees are in anomaly zone

##### 1.1.1 26-taxa model species tree

For the 26-taxon-Sim model species tree, Camin-Sokal parsimony recovers the incorrect species tree topology, returning a balanced topology instead of a pectinate topology for clade $\{A, B, C, D, E, F, G, H\}$; specifically, Camin-Sokal parsimony returns a tree with clade $C = ((AB, CD), (EF, GH))$ instead of clade $C^* = ((((AB), (CD)), (EF)), GH)$ with $XY$ representing the sibling pair $(X, Y)$. To evaluate whether the 26-taxa model species tree is in the anomaly zone (AZ), we used PhyloNet version 3.8.2 (Than et al. 2008; Yu et al. 2014; Wen et al. 2018) and PRANC (Kim et al. 2019). We found that a gene tree with clade $C^*$, which agrees with the true (i.e. model) species tree, is more probable than a gene tree with clade $C$, so we cannot conlude that the 26-taxa model species tree is in the AZ. That being said, the probabilities of the two gene tree topologies were quite close, demonstrating that this is indeed a challenging model condition.

We ran PhyloNet given the following NEXUS file:

```
#NEXUS

BEGIN TREES;
tree gt0 = ((((((A,B),(C,D)),(E,F)),(G,H)),((I,J),(K,L))),(((M,N),(O,P)),((Q,R),(S,T)))),((U,V),(((W,X),Y),Z)));
tree gt1 = (((((A,B),(C,D)),((E,F),(G,H))),((I,J),(K,L))),(((M,N),(O,P)),((Q,R),(S,T)))),((U,V),(((W,X),Y),Z)));
END;

BEGIN NETWORKS;
Network st = ((((((((A:0.1,B:0.1):0.2,(C:0.05,D:0.05):0.25):0.4,(E:0.4,F:0.4):0.3):0.1,(G:0.4,H:0.4):0.4):1.3,
((I:0.5,J:0.5):0.7,(K:0.7,L:0.7):0.5):0.9):3.6,(((M:0.3,N:0.3):0.6,(O:0.1,P:0.1):0.8):4.1,((Q:1.5,R:1.5):1.5,
(S:0.1,T:0.1):2.9):2.0):0.7):7.0,((U:1.9,V:1.9):6.0,(((W:2.0,X:2.0):1.9,Y:3.9):1.5,Z:5.4):2.5):4.8);
END;

BEGIN PHYLONET;
CalGTProb st (gt0);
CalGTProb st (gt1);
END;
```

This yielded the following output:

```
CalGTProb st (gt0)
Species Network:
(((Z:5.4,(Y:3.9,(X:2.0,W:2.0):1.9):1.5):2.5,(V:1.9,U:1.9):6.0):4.8,((((T:0.1,S:0.1):2.9,(R:1.5,Q:1.5):1.5):2.0,
((P:0.1,O:0.1):0.8,(N:0.3,M:0.3):0.6):4.1):0.7,(((L:0.7,K:0.7):0.5,(J:0.5,I:0.5):0.7):0.9,((H:0.4,G:0.4):0.4,
((F:0.4,E:0.4):0.3,((D:0.05,C:0.05):0.25,(B:0.1,A:0.1):0.2):0.4):0.1):1.3):3.6):7.0);
Total log probability: -8.855600603349824

CalGTProb st (gt1)
Species Network:
(((Z:5.4,(Y:3.9,(X:2.0,W:2.0):1.9):1.5):2.5,(V:1.9,U:1.9):6.0):4.8,((((T:0.1,S:0.1):2.9,(R:1.5,Q:1.5):1.5):2.0,
((P:0.1,O:0.1):0.8,(N:0.3,M:0.3):0.6):4.1):0.7,(((L:0.7,K:0.7):0.5,(J:0.5,I:0.5):0.7):0.9,((H:0.4,G:0.4):0.4,
((F:0.4,E:0.4):0.3,((D:0.05,C:0.05):0.25,(B:0.1,A:0.1):0.2):0.4):0.1):1.3):3.6):7.0);
Total log probability: -8.946879528655625
```

Therefore, using PhyloNet's method for computing the probability of rooted gene trees, the gene tree with clade $C^*$ has probability $e^{-8.8556\ldots} = 0.000143$ and the gene tree with clade $C$ has probability $e^{-8.9468\ldots} = 0.000130$. When this analysis is performed for trees restricted to taxa $A--G$, $C^*$ has probability $e^{-5.2635\ldots} = 0.00518$ and $C$ has probability $e^{-5.3354\ldots} = 0.00482$ We found similar results using PRANC by putting the gene trees and species tree in text files and running PRANC as follows:

```
./pranc -uprob st.txt gt0.txt
./pranc -uprob st.txt gt1.txt
```

produced the outputs:

```
0.0051771
0.0048178
```

The values above match the PhyloNet analysis. Below we provide the NEXUS files to run PhyloNet for the remaining model species trees.

###### 1.1.2 4-ingroup-taxa-AZ model species tree

```
#NEXUS

BEGIN TREES;
tree gt0 = ((((A,B),C),D),Out);
tree gt1 = (((A,B),(C,D)),Out);
END;

BEGIN NETWORKS;
Network st = ((((A:0.2,B:0.2):0.01,C:0.21):0.01,D:0.22):20.0,Out:20.22);
END;

BEGIN PHYLONET;
CalGTProb st (gt0);
CalGTProb st (gt1);
END;
```

###### 1.1.3 5-ingroup-taxa-AZ model species tree

```
#NEXUS

BEGIN TREES;
tree gt0 = (((((A,B),C),D),E),Out);
tree gt1 = ((((A,B),C),(D,E)),Out);
tree gt2 = ((((A,B),(C,D)),E),Out);
END;

BEGIN NETWORKS;
Network st = (((((A:0.1,B:0.1):0.1,C:0.2):0.01,D:0.21):0.01,E:0.22):20,Out:20.22);
END;

BEGIN PHYLONET;
CalGTProb st (gt0);
CalGTProb st (gt1);
CalGTProb st (gt2);
END;
```

##### 1.1.4 Palaeognathae-GT-AZ model species tree

```
#NEXUS

BEGIN TREES;
tree gt0 = (galGal,((((((aptOwe,aptHaa),aptRow),(casCas,droNov)),(rheAme,rhePen)),((notPer,eudEle),
(tinGut,cryCin))),strCam));
tree gt1 = (galGal,(((((aptOwe,aptHaa),aptRow),(casCas,droNov)),((rhePen,rheAme),((notPer,eudEle),
(tinGut,cryCin)))),strCam));
END;

BEGIN NETWORKS;
Network st = (galGal:29.0,((((((aptOwe:4.462562,aptHaa:4.462562):1.0655,aptRow:5.528062):3.01202,(casCas:6.585402,
droNov:6.585402):1.95468):0.0531672,(rheAme:4.42582,rhePen:4.42582):4.16743):0.0193593,((notPer:5.216278,
eudEle:5.216278):0.309131,(tinGut:4.373979,cryCin:4.373979):1.15143):3.0872):0.387391,strCam:9.0):20.0);
END;

BEGIN PHYLONET;
CalGTProb st (gt0);
CalGTProb st (gt1);
END;
```

#### 1.2 Commands for simulating data sets

##### 4-ingroup-taxa-AZ model species tree:

```
ms 5 4000000 -t 5.0 -I 5 1 1 1 1 1 \
    -ej 0.1 2 1 -ej 0.105 3 1 -ej .11 4 1 -ej 10.11 5 1 \
    -T -s 1
```

##### 5-ingroup-taxa-AZ model species tree:

```
ms 6 2400000 -t 5.0 -I 6 1 1 1 1 1 1 \
    -ej 0.05 2 1 -ej 0.1 3 1 -ej 0.105 4 1 -ej 0.11 5 1 -ej 10.11 6 1 \
    -T -s 1
```

##### Palaeognathae-GT-AZ model species tree:

```
ms 13 1000000 -t 5.0 -I 13 1 1 1 1 1 1 1 1 1 1 1 1 1 \
    -ej 2.231281 2 1 -ej 2.764031 3 1 -ej 3.292701 5 4 -ej 4.27005 4 1 -ej 2.21291 7 6 \
    -ej 4.296625 6 1 -ej 2.608139 9 8 -ej 2.18699 11 10 -ej 2.762705 10 8 -ej 4.306305 8 1 \
    -ej 4.5 12 1 -ej 14.5 13 1 \
    -T -s 1
```

##### 26-taxa model species tree:

```
ms 26 200000 -t 5.0 -I 26 1 1 1 1 1 1 1 1 1 1 1 1 1 1 1 1 1 1 1 1 1 1 1 1 1 1 \
    -ej 0.05 2 1 -ej 0.025 4 3 -ej 0.15 3 1 -ej 0.2 6 5 -ej 0.35 5 1 \
    -ej 0.2 8 7 -ej 0.4 7 1 -ej 0.25 10 9 -ej 0.35 12 11 -ej 0.6 11 9 \
    -ej 1.05 9 1 -ej 0.15 14 13 -ej 0.05 16 15 -ej 0.45 15 13 -ej 0.75 18 17 \
    -ej 0.05 20 19 -ej 1.5 19 17 -ej 2.5 17 13 -ej 2.85 13 1 -ej 0.95 22 21 \
    -ej 1.0 24 23 -ej 1.95 25 23 -ej 2.7 26 23 -ej 3.95 23 21 -ej 6.35 21 1 \
    -T -s 1
```

#### 1.3 Commands for estimating species trees

##### 1.3.1 Parsimony Methods for 0/1 character states

Below we provide an example NEXUS file for running unordered parsimony using PAUP* version 4.0a168 (Swofford 2002).

```
#NEXUS
BEGIN PAUP
set autoclose=yes warntree=no warnreset=no;
execute <input.nex>
outgroup <name_of_outgroup_name>;
bandb;
contree all/strict=yes treefile=<output_sc.nex> format=newick;
savetrees File=<output_all.nex> root=yes trees=all format=newick;
END;
```

Note that

- To use Dollo parsimony (instead of unordered parsimony), we added the line `ctype dollo:1-100000;` after the line starting with `outgroup`. Similarly, to use Camin-Sokal parsimony, we added the line `ctype irrev:1-100000;[irrev=Camin-Sokal]`.
- To run on different numbers of RIs, we used the `exclude` command, for example the command `exclude 101-100000;` ensures that parsimony is run on the first 100 RIs (even though the input nexus file contained 100,000 RIs).

- To use heuristic search instead of branch-and-bound, we replaced the line starting with `bandb;` with `hsearch addSeq=random nreps=100 swap=TBR;`. This runs 100 heuristic searches that operates by using randomized taxon addition to build a tree on the full taxon set and then applying Tree Bisection and Reconnection (TBR) moves.

##### 1.3.2 SDPquartets

The script for running SDPquartets can be found here: `https://github.com/dbsloan/SDPquartets`.

##### 1.3.3 ASTRAL_BP

The script for running ASTRAL_BP can be found here: `https://github.com/ekmolloy/retrosim-study/run_astral_bp.py`. After running ASTRAL_BP, we ran ASTRID and MDC on the same input, referring to these methods as ASTRID_BP and MDC_BP, respectively.

##### 1.3.4 ASTRID_BP

ASTRID v2.2.1 was run using the command

```
./ASTRID-osx -i <input file> -o <output tree file> &> <output log file>
```

##### 1.3.5 MDC_BP

Below we provide an example NEXUS file for running MDC for unrooted and potentially unresolved gene trees (Yu et al. 2011) using PhyloNet version 3.8.2 (Than et al. 2008; Wen et al. 2018). Note that the "-ur" option, which allows the species tree to be unresolved, is recommended when the input gene trees are unresolved; we ran MDC with and without this option.

```
#NEXUS

BEGIN NETWORKS;
Network g1 = <newick string for RI 1>\;
Network g2 = <newick string for RI 2>\;
...
Network gM = <newick string for RI M>\;
END;

BEGIN PHYLONET;
Infer_ST_MDC_UR (gt1, gt2, ... gtM) <output file>;
END;
```

#### 2 Quartet Probabilities

Kuritzin et al. (2016) and Doronina et al. (2017) model retroelement insertion (RI)s under the MSC model with insertions following an infinite sites neutral mutation model. They use $\omega_{i,j}$ to represent the scenario where a RI in the orthologous locus is absent (0) from lineages $A_i$ and $A_j$ and present (1) in lineages $A_k$ and $A_l$, so

- $\omega_{1,2} = 0011$ and $\omega_{3,4} = 1100$ both display quartet $A_1A_2|A_3A_4$,
- $\omega_{1,3} = 0101$ and $\omega_{2,4} = 1010$ both display quartet: $A_1A_3|X_2A_4$, and
- $\omega_{1,4} = 0110$ and $\omega_{2,3} = 1001$ both display quartet: $A_1A_4|A_2A_3$.

Doronina et al. (2017) derive an approximation for the expected number $a_{i,j}$ of RIs with property $\omega_{i,j}$ for three different phylogenetic networks on four species based on the diffusion approximation of the Wright-Fisher coalescent model (Fisher 1922; Wright 1931) and the neutral mutation model (Kimura 1955a,b). Their "Hybridization model 1" is equivalent to

- a pectinate species tree $(((A_4, A_3), A_2), A_1)$ when $\gamma_1 = 0$ and $\gamma_2 = 1$ and

- a balanced model species tree $((A_4,A_3),(A_2, A_1))$ when $\gamma_1 = 1$ and $\gamma_2 = 0$

(see Figure 6 in Doronina et al. (2017) Supplemental Materials S1). We simplify the equations that they derived for "Hybridization model 1" in order to compute the probability of observing RIs corresponding to each quartet $A_iA_j|A_kA_l$:

$$p^R_{i,j|k,l} = \frac{a_{i,j} + a_{k,l}}{a_{i,j} + a_{i,k} + a_{i,l} + a_{j,k} + a_{j,l} + a_{l,k}} \tag{1}$$

We then verify that $p^R_{1,2|3,4} > p^R_{1,3|2,4} = p^R_{1,4|2,3}$ for the pectinate and balanced model species trees with unrooted topology: $A_1A_2|A_3A_4$. This is summarized in the following theorem.

**Theorem 1.** *Suppose that RIs are generated under the MSC+infinite-sites (as approximated by Doronina et al. 2017). Then, the most probable quartet agrees with the unrooted species tree, and the two alternative quartets have equal probability.*

*Proof.* For the **pectinate model species tree**, let $\tau_3$ be the length (in CUs) of the internal branch separating $A_2, A_3, A_4$ from $A_1$, and let $\tau_2$ be the length (in CUs) of the internal branch separating $A_3, A_4$ from $A_1, A_2$. Let $n_i$ be the expected number of new RIs per generation on the branch with length $\tau_i$ or on the above the root population when $i = 0$ (note that $n_i$ is the probability of a new RI occurring in an individual times the effective population size). Simplifying the equations from Doronina et al. (2017), the expected number of RIs that display the quartet topology that agrees with the unrooted model species tree (i.e. $A_1, A_2|A_3, A_4$) is

$$\begin{aligned} a_{1,2} + a_{3,4} = n_0 &\left(e^{-\tau_2} - \frac{1}{2}e^{-\tau_2-\tau_3} - \frac{1}{6}e^{-3\tau_2-\tau_3}\right) \\ &+ n_2\left(1 - e^{-\tau_2} - \frac{2}{3}e^{-\tau_3} + \frac{1}{2}e^{-\tau_2-\tau_3} + \frac{1}{6}e^{-3\tau_2-\tau_3}\right) \\ &+ n_3\big(\tau_3 - 1 + e^{-\tau_3}\big) \end{aligned} \tag{2}$$

and the expected number of RIs that display one of the two alternative quartets (i.e. $A_1, A_3|A_2, A_4$ and $A_1, A_4|A_2, A_3$) is

$$\begin{aligned} a_{1,3} + a_{2,4} = a_{1,4} + a_{2,3} = n_0 &\left(\frac{1}{2}e^{-\tau_2-\tau_3} - \frac{1}{6}e^{-3\tau_2-\tau_3}\right) \\ &+ n_2\left(\frac{1}{3}e^{-\tau_3} - \frac{1}{2}e^{-\tau_2-\tau_3} + \frac{1}{6}e^{-3\tau_2-\tau_3}\right) \end{aligned} \tag{3}$$

(see Section 2.4 below for details). Now we verify that

$$\begin{aligned} (p^R_{1,2} + p^R_{3,4}) - (p^R_{1,3} + p^R_{2,4}) &> 0 \\ (a_{1,2} + a_{3,4}) - (a_{1,3} + a_{2,4}) &> 0 \\ n_0\left(e^{-\tau_2} - \frac{1}{2}e^{-\tau_2-\tau_3} - \frac{1}{6}e^{-3\tau_2-\tau_3}\right) - n_0\left(\frac{1}{2}e^{-\tau_2-\tau_3} - \frac{1}{6}e^{-3\tau_2-\tau_3}\right) & \\ +n_2\left(1 - e^{-\tau_2} - \frac{2}{3}e^{-\tau_3} + \frac{1}{2}e^{-\tau_2-\tau_3} + \frac{1}{6}e^{-3\tau_2-\tau_3}\right) - n_2\left(\frac{1}{3}e^{-\tau_3} - \frac{1}{2}e^{-\tau_2-\tau_3} + \frac{1}{6}e^{-3\tau_2-\tau_3}\right) & \\ +n_3\big(\tau_3 - 1 + e^{-\tau_3}\big) &> 0 \\ n_0\left(e^{-\tau_2} - e^{-\tau_2-\tau_3} + \frac{1}{3}e^{-3\tau_2-\tau_3}\right) + n_2\left(1 - e^{-\tau_2} - e^{-\tau_3} + e^{-\tau_2-\tau_3}\right) + n_3\big(\tau_3 - 1 + e^{-\tau_3}\big) &> 0. \end{aligned}$$

This inequality holds for $n_0, n_1, n_2, \tau_1, \tau_2 > 0$, because the first term is positive by

$$\begin{aligned} 1 &> e^{-\tau_2} \\ 1 - e^{-\tau_2} &> 0 \\ (1 - e^{-\tau_2}) \times e^{-\tau_3} &> 0 \times e^{-\tau_3} \\ e^{-\tau_2} - e^{-\tau_2-\tau_3} &> 0, \end{aligned}$$

the second term is positive by $1 - e^{-\tau_2} - e^{-\tau_3} + e^{-\tau_2 - \tau_3} = (1 - e^{-\tau_2})(1 - e^{-\tau_3})$ and the third term is positive by $1 - \tau_2 < e^{-\tau_2}$ (Bernoulli's inequality). For the pectinate model species tree, the most probable quartet on species $A_1$, $A_2$, $A_3$, and $A_4$ agrees with the species tree on species $A_1$, $A_2$, $A_3$, and $A_4$, and the two alternative quartet trees have equal probability.

For the **balanced model species tree**, let $\tau_1$ be the length (in CUs) of the internal branch above $A_1, A_2$, and let $\tau_3$ be the length (in CUs) of the internal branch above $A_3, A_4$. Let $n_i$ be the expected number of new RIs per generation corresponding to the same branch as length $\tau_i$ or the above the root population when $i = 0$. Simplifying the equations from Doronina et al. (2017), the expected number of RIs that display the quartet topology that agrees with the species tree (i.e. $A_1, A_2|A_3, A_4$) is

$$a_{1,2} + a_{3,4} = n_0\left(2 - e^{-\tau_1} - e^{-\tau_3} + \frac{1}{3}e^{-\tau_1-\tau_3}\right) + n_1\big(\tau_1 - 1 + e^{-\tau_1}\big) \\ + n_3\big(\tau_3 - 1 + e^{-\tau_3}\big), \tag{4}$$

and the expected number of RIs that display one of the two alternative quartets (i.e. $A_1, A_3|A_2, A_3$ and $A_1, A_4|A_2, A_3$ ) is

$$a_{1,3} + a_{2,4} = a_{1,4} + a_{2,3} = n_0\frac{1}{3}e^{-\tau_1-\tau_3}. \tag{5}$$

(see Section 2.5 below for details). Now we verify that

$$\begin{aligned}(p^R_{1,2} + p^R_{3,4}) - (p^R_{1,3} + p^R_{2,4}) &> 0\\ (a_{1,2} + a_{3,4}) - (a_{1,3} + a_{2,4}) &> 0\\ n_0\left(2 - e^{-\tau_1} - e^{-\tau_3} + \frac{1}{3}e^{-\tau_1-\tau_3}\right) - n_0\frac{1}{3}e^{-\tau_1-\tau_3} + n_1\big(\tau_1 - 1 + e^{-\tau_1}\big) + n_2\big(\tau_3 - 1 + e^{-\tau_3}\big) &> 0\\ n_0\big(2 - e^{-\tau_1} - e^{-\tau_3}\big) + n_1\big(\tau_1 - 1 + e^{-\tau_1}\big) + n_2\big(\tau_3 - 1 + e^{-\tau_3}\big) &> 0\end{aligned}$$

This inequality holds for $n_0, n_1, n_2, \tau_1, \tau_3 > 0$, because the first term is positive as $1 > e^{-\tau_i}$ and the second and third terms are positive as $e^{-\tau_i} > 1 - \tau_i$ (Bernoulli's inequality). For the balanced model species tree, the most probable quartet on species $A_1$, $A_2$, $A_3$, and $A_4$ agrees with the species tree on species $A_1$, $A_2$, $A_3$, and $A_4$, and the alternative two quartet trees have equal probability. $\square$

The theorem above enables proofs of statistical consistency for two different quartet-based methods: SDPquartets and ASTRAL_BP.

#### 2.1 Quartet probabilities for short internal branches

We now consider what happens when **both** internal branches are short enough so that the small angle approximation can be applied.

For the **pectinate model species tree**, suppose both $\tau_2$ and $\tau_3$ are sufficiently short, then we can apply the small angle approximation $e^{-\tau_i} \approx 1 - \tau_i$ and drop the higher order terms (e.g. $\tau_2\tau_3$ and $\tau_2^2$). From

Equation 2, the expected number of RIs displaying the quartet that agrees with the species tree is

$$\begin{aligned}
a_{1,2}+a_{3,4} &= n_0\left(e^{-\tau_2}-\frac{1}{2}e^{-\tau_2-\tau_3}-\frac{1}{6}e^{-3\tau_2-\tau_3}\right)\\
&\quad + n_2\left(1-e^{-\tau_2}-\frac{2}{3}e^{-\tau_3}+\frac{1}{2}e^{-\tau_2-\tau_3}+\frac{1}{6}e^{-3\tau_2-\tau_3}\right)\\
&\quad + n_3\left(\tau_3-1+e^{-\tau_3}\right)\\
&\approx n_0\left((1-\tau_2)-\frac{1}{2}(1-\tau_2)(1-\tau_3)-\frac{1}{6}(1-\tau_2)^3(1-\tau_3)\right)\\
&\quad + n_2\left(1-(1-\tau_2)-\frac{2}{3}(1-\tau_3)+\frac{1}{2}(1-\tau_2)(1-\tau_3)+\frac{1}{6}(1-\tau_2)^3(1-\tau_3)\right)\\
&\quad + n_3\left(\tau_3-1+(1-\tau_3)\right)\\
&\approx n_0\left((1-\tau_2)-\frac{1}{2}(1-\tau_2-\tau_3)-\frac{1}{6}(1-3\tau_2-\tau_3)\right)\\
&\quad + n_2\left(1-(1-\tau_2)-\frac{2}{3}(1-\tau_3)+\frac{1}{2}(1-\tau_2-\tau_3)+\frac{1}{6}(1-3\tau_2-\tau_3)\right)\\
&= n_0\left(\frac{1}{3}+\frac{2}{3}\tau_3\right)
\end{aligned}$$

and from Equation 3, the expected number of RIs displaying the alternative quartets is

$$a_{1,3}+a_{2,4}=a_{1,4}+a_{2,3}\approx n_0\left(\frac{1}{3}-\frac{1}{3}\tau_3\right)$$

(see Section 2.4 below for details). Repeating this approximation for the **balanced model species tree** using Equations 4 and 5 gives

$$a_{1,2}+a_{3,4}\approx n_0\left(\frac{1}{3}+\frac{2}{3}(\tau_1+\tau_3)\right) \text{ and } a_{1,3}+a_{2,4}=a_{1,4}+a_{2,3}\approx n_0\left(\frac{1}{3}-\frac{1}{3}(\tau_1+\tau_3)\right)$$

(see Section 2.5 below for details). Plugging these formulas into Equation 1 gives

$$p^R_{1,2|3,4}\approx\frac{1}{3}+\frac{2}{3}\tau \quad\text{and}\quad p^R_{1,3|2,4}=p^R_{1,4|2,3}\approx\frac{1}{3}-\frac{1}{3}\tau \tag{6}$$

where $\tau$ is the length of the internal branch that induces quartet $A_1, A_2|A_3, A_4$. For the pectinate model species tree, $\tau=\tau_3$ (but recall that $\tau_2$ must also be short for the approximation to apply) and $\tau=\tau_1+\tau_3$ for the balanced model species tree.

#### 2.2 Quartet probabilities when the expected number of new RIs per generation is constant

We now consider what happens when the expected number of RIs per generation is constant across the species tree.

For the **pectinate model species tree**, we set $n_0=n_2=n_3$. From Equation 2, the expected number of RIs displaying the quartet that agrees with the species tree is

$$\begin{aligned}
a_{1,2} + a_{3,4} &= n_0\left(e^{-\tau_2} - \frac{1}{2}e^{-\tau_2-\tau_3} - \frac{1}{6}e^{-3\tau_2-\tau_3}\right) \\
&\quad + n_2\left(1 - e^{-\tau_2} - \frac{2}{3}e^{-\tau_3} + \frac{1}{2}e^{-\tau_2-\tau_3} + \frac{1}{6}e^{-3\tau_2-\tau_3}\right) \\
&\quad + n_3\left(\tau_3 - 1 + e^{-\tau_3}\right) \\
&= n_3\tau_3 + n_3e^{-\tau_3} - n_2\frac{2}{3}e^{-\tau_3} + (n_2 - n_3) + (n_0 - n_2)\left(e^{-\tau_2} - \frac{1}{2}e^{-\tau_2-\tau_3} - \frac{1}{6}e^{-3\tau_2-\tau_3}\right) \\
&= n_0\left(\tau_3 + \frac{1}{3}e^{-\tau_3}\right)
\end{aligned}$$

and from Equation 3, the expected number of RIs displaying the alternative quartets is

$$\begin{aligned}
a_{1,3} + a_{2,4} = a_{1,4} + a_{2,3} &= n_0\left(\frac{1}{2}e^{-\tau_2-\tau_3} - \frac{1}{6}e^{-3\tau_2-\tau_3}\right) + n_2\left(\frac{1}{3}e^{-\tau_3} - \frac{1}{2}e^{-\tau_2-\tau_3} + \frac{1}{6}e^{-3\tau_2-\tau_3}\right) \\
&= n_2\frac{1}{3}e^{-\tau_3} + (n_0 - n_2)\left(\frac{1}{2}e^{-\tau_2-\tau_3} - \frac{1}{6}e^{-3\tau_2-\tau_3}\right) \\
&= n_0\frac{1}{3}e^{-\tau_3}
\end{aligned}$$

(see Section 2.4 below for details). Repeating this simplification (i.e. $n_0 = n_1 = n_3$) for the **balanced model species tree** using Equations 4 and 5 gives

$$a_{1,2} + a_{3,4} = n_0\left((\tau_1 + \tau_3) + \frac{1}{3}e^{-(\tau_1+\tau_3)}\right) \text{ and } a_{1,3} + a_{2,4} = a_{1,4} + a_{2,3} = n_0\frac{1}{3}e^{-(\tau_1+\tau_3)}$$

(see Section 2.5 below for details). Plugging these formulas into Equation 1 gives

$$p^R_{1,2|3,4} = \frac{\frac{1}{3}e^{-\tau} + \tau}{e^{-\tau} + \tau} \quad \text{and} \quad p^R_{1,3|2,4} = p^R_{1,4|2,3} = \frac{\frac{1}{3}e^{-\tau}}{e^{-\tau} + \tau} \tag{7}$$

where $\tau$ is the length of the internal branch that induces quartet $A_1, A_2|A_3, A_4$. For the pectinate model species tree, $\tau = \tau_3$ and $\tau = \tau_1 + \tau_3$ for the balanced model species tree.

#### 2.3 Simplifying Equations from Doronina et al. (2017) – Pectinate species tree

By setting $\gamma_1 = 0$ and $\gamma_2 = 1$, we obtain the pectinate species tree with topology $(((A_3,A_4){:}T_3,A_2){:}T_2,A_1)$, where $T_3 = t_3 - t_2$ and $T_2 = t_2 - t_0$ generations (see Figure 6 in Doronina et al. (2017) Supplemental Materials S1). Let $\tau_2$ and $\tau_3$ be the branch lengths in CUs, and let $n_i$ denote the expected number of insertions per generation on the branch with length $T_i$, with $i = 0$ indicating the population above the root.

##### 2.3.1 Ancestral population for $A_1$, $A_2$, $A_3$, and $A_4$

First, we consider the case where a RI originates in the ancestral population for species $A_1$, $A_2$, $A_3$, and $A_4$. Taking the equations for $a^0_{i,j}$ at the top of page 5 of Doronina et al. (2017) Supplemental Materials S1 and setting $\gamma_1 = 0$, $\gamma_2 = 1$ gives us the following set of equations:

$$a^0_{1,2} = n_0\left(\frac{2}{3}e^{-\tau_2} - \frac{1}{6}e^{-3\tau_2-\tau_3} - \frac{1}{3}e^{-\tau_2-\tau_3}\right),$$

$$a^0_{1,3} = a^0_{1,4} = n_0\left(-\frac{1}{6}e^{-3\tau_2-\tau_3} + \frac{1}{3}e^{-\tau_2-\tau_3}\right),$$

$$a^0_{2,3} = a^0_{2,4} = n_0\left(\frac{1}{6}e^{-\tau_2-\tau_3}\right),$$

and

$$a^0_{3,4} = n_0\left(\frac{1}{3}e^{-\tau_2} - \frac{1}{6}e^{-\tau_2-\tau_3}\right).$$

###### 2.3.2 Ancestral population for $A_2$, $A_3$ and $A_4$ on branch $T_2$

Second, we consider the case where a RI originates in the ancestral population for species $A_2$, $A_3$, and $A_4$. Taking the equations for $a^2_{i,j}$ at bottom of page 7 of Doronina et al. (2017) Supplemental Materials S1 and setting $\gamma_1 = 0$, $\gamma_2 = 1$, gives us the following set of equations:

$$\begin{aligned}
a^2_{1,2} &= n_2\big(1-e^{-\tau_2}\big)\left(1-\frac{1}{6}\big(e^{-2\tau_2}+e^{-\tau_2}+4\big)e^{-\tau_3}\right)\\
&= n_2\big(1-e^{-\tau_2}\big)\left(1-\frac{1}{6}\big(e^{-2\tau_2-\tau_3}+e^{-\tau_2-\tau_3}+4e^{-\tau_3}\big)\right)\\
&= n_2\big(1-e^{-\tau_2}\big)\left(1-\frac{1}{6}e^{-2\tau_2-\tau_3}-\frac{1}{6}e^{-\tau_2-\tau_3}-\frac{4}{6}e^{-\tau_3}\right)\\
&= n_2\left(1-\frac{1}{6}e^{-2\tau_2-\tau_3}-\frac{1}{6}e^{-\tau_2-\tau_3}-\frac{4}{6}e^{-\tau_3}-e^{-\tau_2}+\frac{1}{6}e^{-3\tau_2-\tau_3}+\frac{1}{6}e^{-2\tau_2-\tau_3}+\frac{4}{6}e^{-\tau_2-\tau_3}\right)\\
&= n_2\left(1-e^{-\tau_2}-\frac{2}{3}e^{-\tau_3}+\frac{1}{2}e^{-\tau_2-\tau_3}+\frac{1}{6}e^{-3\tau_2-\tau_3}\right),
\end{aligned}$$

$$\begin{aligned}
a^2_{1,3} = a^2_{1,4} &= n_2\frac{1}{6}e^{-\tau_3}(1-e^{-\tau_2})(1-e^{-\tau_2})(2+e^{-\tau_2})\\
&= n_2\frac{1}{6}e^{-\tau_3}(1-2e^{-\tau_2}+e^{-2\tau_2})(2+e^{-\tau_2})\\
&= n_2\frac{1}{6}e^{-\tau_3}(2+e^{-\tau_2}-4e^{-\tau_2}-2e^{-2\tau_2}+2e^{-2\tau_2}+e^{-3\tau_2})\\
&= n_2\frac{1}{6}e^{-\tau_3}(2-3e^{-\tau_2}+e^{-3\tau_2})\\
&= n_2\left(\frac{1}{3}e^{-\tau_3}-\frac{1}{2}e^{-\tau_2-\tau_3}+\frac{1}{6}e^{-3\tau_2-\tau_3}\right),
\end{aligned}$$

and

$$a^2_{2,3} = a^2_{2,4} = 0.$$

###### 2.3.3 Ancestral population for $A_3$ and $A_4$ on branch $T_3$

Third, we consider the case where a RI originates in the ancestral population for species $A_3$ and $A_4$. Taking the equation for $a^3_{i,j}$ on page 8 Doronina et al. (2017) Supplemental Materials S1 and setting $\gamma_1 = 0$, $\gamma_2 = 1$, gives us the following set of equations:

$$a^3_{1,2} = n_3\big(\tau_3 - 1 + e^{-\tau_3}\big)$$

and

$$a^3_{1,3} = a^3_{1,4} = a^3_{2,3} = a^3_{2,4} = a^3_{3,4} = 0.$$

###### 2.3.4 Putting it all together

Now, we can sum the expected number of RIs that display pattern $\omega_{i,j}$ across all branches and above the root (i.e. we can compute $a_{i,j} = a^0_{i,j} + a^2_{i,j} + a^3_{i,j}$) and combine these values based on whether the RIs patterns correspond to the same quartet topology. This gives us the following set of equations:

$$\begin{aligned}
a_{1,2} + a_{3,4} &= n_0\left(e^{-\tau_2} - \frac{1}{2}e^{-\tau_2-\tau_3} - \frac{1}{6}e^{-3\tau_2-\tau_3}\right) \\
&\quad + n_2\left(1 - e^{-\tau_2} - \frac{2}{3}e^{-\tau_3} + \frac{1}{2}e^{-\tau_2-\tau_3} + \frac{1}{6}e^{-3\tau_2-\tau_3}\right) \\
&\quad + n_3\left(\tau_3 - 1 + e^{-\tau_3}\right) \\
&= n_3\tau_3 + n_3e^{-\tau_3} - n_2\frac{2}{3}e^{-\tau_3} + (n_2 - n_3) + (n_0 - n_2)\left(e^{-\tau_2} - \frac{1}{2}e^{-\tau_2-\tau_3} - \frac{1}{6}e^{-3\tau_2-\tau_3}\right)
\end{aligned}$$

and

$$\begin{aligned}
a_{1,3} + a_{2,4} = a_{1,4} + a_{2,3} &= n_0\left(\frac{1}{2}e^{-\tau_2-\tau_3} - \frac{1}{6}e^{-3\tau_2-\tau_3}\right) + n_2\left(\frac{1}{3}e^{-\tau_3} - \frac{1}{2}e^{-\tau_2-\tau_3} + \frac{1}{6}e^{-3\tau_2-\tau_3}\right) \\
&= n_2\frac{1}{3}e^{-\tau_3} + (n_0 - n_2)\left(\frac{1}{2}e^{-\tau_2-\tau_3} - \frac{1}{6}e^{-3\tau_2-\tau_3}\right)
\end{aligned}$$

###### 2.3.5 Short internal branches

If **both** $\tau_2$ and $\tau_3$ are sufficiently short, then we can apply the small angle approximation $e^{-x} \approx 1 - x$. Then, we can also drop the higher order terms: $\tau_2\tau_3$ and $\tau_2^2$.

$$\begin{aligned}
a_{1,2} + a_{3,4} &= n_0\left(e^{-\tau_2} - \frac{1}{2}e^{-\tau_2-\tau_3} - \frac{1}{6}e^{-3\tau_2-\tau_3}\right) \\
&\quad + n_2\left(1 - e^{-\tau_2} - \frac{2}{3}e^{-\tau_3} + \frac{1}{2}e^{-\tau_2-\tau_3} + \frac{1}{6}e^{-3\tau_2-\tau_3}\right) \\
&\quad + n_3\left(\tau_3 - 1 + e^{-\tau_3}\right) \\
&\approx n_0\left((1-\tau_2) - \frac{1}{2}(1-\tau_2)(1-\tau_3) - \frac{1}{6}(1-\tau_2)^3(1-\tau_3)\right) \\
&\quad + n_2\left(1 - (1-\tau_2) - \frac{2}{3}(1-\tau_3) + \frac{1}{2}(1-\tau_2)(1-\tau_3) + \frac{1}{6}(1-\tau_2)^3(1-\tau_3)\right) \\
&\quad + n_3\left(\tau_3 - 1 + (1-\tau_3)\right) \\
&\approx n_0\left((1-\tau_2) - \frac{1}{2}(1-\tau_2-\tau_3+\tau_2\tau_3) - \frac{1}{6}(1-3\tau_2-\tau_3+3\tau_2^2+3\tau_2\tau_3+\dots)\right) \\
&\quad + n_2\left(1 - (1-\tau_2) - \frac{2}{3}(1-\tau_3) + \frac{1}{2}(1-\tau_2-\tau_3+\tau_2\tau_3) + \frac{1}{6}(1-3\tau_2-\tau_3+3\tau_2^2+3\tau_2\tau_3+\dots)\right) \\
&\approx n_0\left(\frac{6}{6} - \frac{6}{6}\tau_2 - \frac{3}{6} + \frac{3}{6}\tau_2 + \frac{3}{6}\tau_3 - \frac{3}{6}\tau_2\tau_3 - \frac{1}{6} + \frac{3}{6}\tau_2 + \frac{1}{6}\tau_3 - \frac{3}{6}\tau_2^2 - \frac{3}{6}\tau_2\tau_3\right) \\
&\quad + n_2\left(\frac{6}{6}\tau_2 - \frac{4}{6} + \frac{4}{6}\tau_3 + \frac{3}{6} - \frac{3}{6}\tau_2 - \frac{3}{6}\tau_3 + \frac{3}{6}\tau_2\tau_3 + \frac{1}{6} - \frac{3}{6}\tau_2 - \frac{1}{6}\tau_3 + \frac{3}{6}\tau_2^2 + \frac{3}{6}\tau_2\tau_3\right) \\
&\approx n_0\left(\frac{1}{3} + \frac{2}{3}\tau_3\right)
\end{aligned}$$

$$
\begin{aligned}
&a_{1,3} + a_{2,4} = a_{1,4} + a_{2,3} \\
&= n_0\left(\frac{1}{2}e^{-\tau_2-\tau_3} - \frac{1}{6}e^{-3\tau_2-\tau_3}\right) + n_2\left(\frac{1}{3}e^{-\tau_3} - \frac{1}{2}e^{-\tau_2-\tau_3} + \frac{1}{6}e^{-3\tau_2-\tau_3}\right) \\
&\approx n_0\left(\frac{1}{2}(1-\tau_2)(1-\tau_3) - \frac{1}{6}(1-\tau_2)^3(1-\tau_3)\right) + n_2\left(\frac{1}{3}(1-\tau_3) - \frac{1}{2}(1-\tau_2)(1-\tau_3) + \frac{1}{6}(1-\tau_2)^3(1-\tau_3)\right) \\
&\approx n_0\left(\frac{1}{2}(1-\tau_2-\tau_3+\tau_2\tau_3) - \frac{1}{6}(1-3\tau_2-\tau_3+3\tau_2^2+3\tau_2\tau_3+\dots)\right) \\
&\quad + n_2\left(\frac{1}{3}(1-\tau_3) - \frac{1}{2}(1-\tau_2-\tau_3+\tau_2\tau_3) + \frac{1}{6}(1-3\tau_2-\tau_3+3\tau_2^2+3\tau_2\tau_3+\dots)\right) \\
&= n_0\left(\frac{3}{6} - \frac{3}{6}\tau_2 - \frac{3}{6}\tau_3 + \frac{3}{6}\tau_2\tau_3 - \frac{1}{6} + \frac{3}{6}\tau_2 + \frac{1}{6}\tau_3 - \frac{3}{6}\tau_2^2 - \frac{3}{6}\tau_2\tau_3)\right) \\
&\quad + n_2\left(\frac{2}{6} - \frac{2}{6}\tau_3 - \frac{3}{6} + \frac{3}{6}\tau_2 + \frac{3}{6}\tau_3 - \frac{3}{6}\tau_2\tau_3 + \frac{1}{6} - \frac{3}{6}\tau_2 - \frac{1}{6}\tau_3 + \frac{3}{6}\tau_2^2 + \frac{3}{6}\tau_2\tau_3\right) \\
&\approx n_0\left(\frac{1}{3} - \frac{1}{3}\tau_3\right)
\end{aligned}
$$

##### 2.4 Simplifying Equations from Doronina et al. (2017) – Balanced species tree

By setting $\gamma_1 = 1$ and $\gamma_2 = 0$, we obtain the pectinate species tree with topology $(((A_3,A_4){:}T_3,(A_1,A_2){:}T_1)$, where $T_1 = t_1 - t_0$ and $T_3 = t_3 - t_0$ generations (see Figure 6 in Doronina et al. (2017) Supplemental Materials S1). Let $\tau_1$ and $\tau_3$ denote the lengths of branch $T_1$ and branch $T_3$ in CUs, and let $n_i$ denote the expected number of insertions per generation on the branch with length $T_i$, with $i = 0$ indicating the population above the root.

###### 2.4.1 Ancestral population for $A_1$, $A_2$, $A_3$, and $A_4$

First, we consider the case where a RI originates in the ancestral population above species $A_1$, $A_2$, $A_3$, and $A_4$. Taking the equations for $a^0_{i,j}$ at the top of page 5 of Doronina et al. (2017) Supplemental Materials S1, setting $\gamma_1 = 1$, $\gamma_2 = 0$, and removing $\tau_2$ gives us the following set of equations:

$$a^0_{1,2} = n_0\left(1 - \frac{1}{3}e^{-\tau_1} - \frac{2}{3}e^{-\tau_3} + \frac{1}{6}e^{-\tau_1-\tau_3}\right),$$

$$a^0_{1,3} = a^0_{1,4} = n_0\left(\frac{1}{6}e^{-\tau_1-\tau_3}\right),$$

$$a^0_{2,3} = a^0_{2,4} = n_0\left(\frac{1}{6}e^{-\tau_1-\tau_3}\right),$$

and

$$a^0_{3,4} = n_0\left(1 - \frac{2}{3}e^{-\tau_1} - \frac{1}{3}e^{-\tau_3} + \frac{1}{6}e^{-\tau_1-\tau_3}\right).$$

###### 2.4.2 Ancestral population for $A_1$ and $A_2$

Second, we consider the case where a RI originates in the ancestral population for species $A_1$ and $A_2$. Taking Equation 5 (for $a^1_{i,j}$) at bottom of page 6 of Doronina et al. (2017) Supplemental Materials S1 and setting $\gamma_1 = 1$ gives us the following set of equations:

$$a^1_{1,2} = a^1_{1,3} = a^1_{1,4} = a^1_{2,3} = a^1_{2,4} = 0$$

and

$$a^1_{3,4} = n_1(\tau_1 - 1 + e^{-\tau_1}).$$

##### 2.4.3 Ancestral population for $A_3$ and $A_4$

Taking the equation for $a^3_{i,j}$ on page 8 Doronina et al. (2017) Supplemental Materials S1 gives us the following set of equations:

$$a^3_{1,2} = n_3\big(\tau_3 - 1 + e^{-\tau_3}\big)$$

and

$$a^3_{1,3} = a^2_{1,4} = a^4_{2,3} = a^4_{2,4} = a^4_{3,4} = 0.$$

##### 2.4.4 Putting it all together

Now, we can sum the expected number of RIs that display pattern $\omega_{i,j}$ across all branches and above the root (i.e. we can compute $a_{i,j} = a^0_{i,j} + a^1_{i,j} + a^3_{i,j}$) and combine these values based on the whether the RIs patterns correspond to the same quartet topology. This gives us the following set of equations:

$$\begin{aligned} a_{1,2} + a_{3,4} &= n_0\left(2 - e^{-\tau_1} - e^{-\tau_3} + \frac{1}{3}e^{-\tau_1-\tau_3}\right) + n_1\big(\tau_1 - 1 + e^{-\tau_1}\big) + n_3\big(\tau_3 - 1 + e^{-\tau_3}\big) \\ &= n_0\frac{1}{3}e^{-\tau_1-\tau_2} + n_1\tau_1 + n_3\tau_3 + (n_0 - n_1)(1 - e^{-\tau_1}) + (n_0 - n_3)(1 - e^{-\tau_3}) \end{aligned}$$

and

$$a_{1,3} + a_{2,4} = a_{1,4} + a_{2,3} = n_0\frac{1}{3}e^{-\tau_1-\tau_3}.$$

##### 2.4.5 Short internal branch

If $\tau_1 + \tau_3$ is sufficiently short, then we can apply the small angle approximation $e^{-x} \approx 1 - x$. Because $\tau_1 > 0$ and $\tau_3 > 0$, we can apply the small angle approximation to $\tau_1$ and $\tau_3$ individually and drop the higher order term: $\tau_1\tau_3$. Then,

$$\begin{aligned} a_{1,2} + a_{3,4} &= n_0\left(2 - e^{-\tau_1} - e^{-\tau_3} + \frac{1}{3}e^{-\tau_1-\tau_3}\right) + n_1\big(\tau_1 - 1 + e^{-\tau_1}\big) + n_3\big(\tau_3 - 1 + e^{-\tau_3}\big) \\ &\approx n_0\left(2 - (1 - \tau_1) - (1 - \tau_3) + \frac{1}{3}(1 - \tau_1)(1 - \tau_3)\right) + n_1\big(\tau_1 - 1 + 1 - \tau_1\big) + n_3\big(\tau_3 - 1 + 1 - \tau_3\big) \\ &= n_0\left(\tau_1 + \tau_3 + \frac{1}{3}(1 - \tau_1)(1 - \tau_3)\right) \\ &= n_0\left(\tau_1 + \tau_3 + \frac{1}{3}(1 - \tau_1 - \tau_3 + \tau_1\tau_3)\right) \\ &\approx n_0\left(\frac{1}{3} + \frac{2}{3}(\tau_1 + \tau_3)\right) \end{aligned}$$

and

$$\begin{aligned} a_{1,3} + a_{2,4} = a_{1,4} + a_{2,3} &= n_0\frac{1}{3}e^{-\tau_1-\tau_3} \approx n_0\frac{1}{3}(1 - \tau_1)(1 - \tau_3) \\ &= n_0\left(\frac{1}{3} - \frac{1}{3}(\tau_1 + \tau_3 + \tau_1\tau_3)\right) \\ &\approx n_0\left(\frac{1}{3} - \frac{1}{3}(\tau_1 + \tau_3)\right). \end{aligned}$$

### 3 Supplementary Results

### 4 Comparing of Branch Length Estimates

Table S1: **Branch lengths for simulated data sets with 100,000 retroelement insertions (RIs).** Branch lengths were estimated using three different techniques (MAP-GT, MLE-GT, and MLE-RI) described in the main text. Percent error is computed as $(abs(\tau^* - \hat{\tau})/\tau^*) \times 100$, where $\tau^*$ is the true branch length and $\hat{\tau}$ is the estimated branch length. In parentheses, we note the number of replicates (out of 25) where the estimated branch length $\hat{\tau}$ was longer than the true branch length $\tau^*$; importantly, the MAP-GT and MLE-GT branch lengths were consistently biased upward (25/25 replicates) whenever the true branch length was greater than 0.25 CUs. $EN$ is the effective number of RIs for that branch; this value can be smaller than the total number of insertions, because an RI represents a single bipartition rather than a fully resolved gene tree. All values not in parentheses are the result of averaging across the 25 replicate data sets. Note that the data used to create this table are available in CSV format in the Supplementary Materials available on Dryad.

| True Branch Length | Estimated Branch Length MAP-GT / MLE-GT / MLE-RI | Percent Error MAP-GT / MLE-GT / MLE-RI | EN |
|---|---|---|---|
| *5-taxon-AZ-Sim* | | | |
| 0.0100 | 0.0102 / 0.0102 / 0.0102 | 25% (15) / 25% (15) / 25% (15) | 60042.94 |
| 0.0100 | 0.0096 / 0.0096 / 0.0096 | 21% (10) / 21% (10) / 21% (10) | 59979.64 |
| *6-taxon-AZ-Sim* | | | |
| 0.0100 | 0.0103 / 0.0103 / 0.0102 | 18% (15) / 18% (15) / 18% (15) | 46017.67 |
| 0.0100 | 0.0103 / 0.0104 / 0.0103 | 18% (12) / 18% (12) / 18% (12) | 46022.51 |
| 0.1000 | 0.1059 / 0.1060 / 0.1010 | 6% (24) / 6% (24) / **2%** (16) | 46261.95 |
| *Palaeognathae-GT-AZ* | | | |
| 0.0194 | 0.0189 / 0.0190 / 0.0188 | 23% (11) / 23% (11) / 23% (11) | 6371.20 |
| 0.0532 | 0.0563 / 0.0564 / 0.0549 | 15% (15) / 15% (15) / **14%** (14) | 6376.60 |
| 0.3091 | 0.3533 / 0.3535 / 0.3108 | 14% (25) / 14% (25) / **2%** (13) | 6656.30 |
| 0.3874 | 0.4521 / 0.4523 / 0.3880 | 17% (25) / 17% (25) / **2%** (14) | 6796.89 |
| 1.0655 | 1.4150 / 1.4156 / 1.0699 | 33% (25) / 33% (25) / **1%** (18) | 9013.62 |
| 1.1514 | 1.5455 / 1.5462 / 1.1590 | 34% (25) / 34% (25) / **1%** (18) | 9357.42 |
| 1.9547 | 2.7077 / 2.7093 / 1.9650 | 38% (25) / 39% (25) / **1%** (18) | 13403.91 |
| 3.0120 | 4.1590 / 4.1639 / 3.0373 | 38% (25) / 38% (25) / **1%** (17) | 19641.24 |
| 3.0872 | 4.2476 / 4.2528 / 3.1054 | 38% (25) / 38% (25) / **1%** (17) | 20061.70 |
| 4.1674 | 5.6047 / 5.6200 / 4.1850 | 34% (25) / 35% (25) / **1%** (15) | 26803.29 |
| *26-taxon-Sim* | | | |
| 0.1000 | 0.1048 / 0.1051 / 0.1002 | 9% (14) / 9% (14) / **8%** (11) | 2117.93 |
| 0.2000 | 0.2225 / 0.2229 / 0.2037 | 12% (24) / 12% (24) / **5%** (16) | 2161.30 |
| 0.2500 | 0.2831 / 0.2835 / 0.2542 | 13% (25) / 13% (25) / **4%** (16) | 2170.60 |
| 0.3000 | 0.3372 / 0.3377 / 0.2981 | 13% (23) / 13% (23) / **5%** (13) | 2197.92 |
| 0.4000 | 0.4657 / 0.4663 / 0.3987 | 16% (25) / 17% (25) / **2%** (13) | 2259.40 |
| 0.4000 | 0.4723 / 0.4729 / 0.4037 | 18% (25) / 18% (25) / **3%** (14) | 2248.28 |
| 0.5000 | 0.6049 / 0.6056 / 0.5032 | 21% (25) / 21% (25) / **3%** (16) | 2323.75 |
| 0.6000 | 0.7414 / 0.7423 / 0.6025 | 24% (25) / 24% (25) / **3%** (14) | 2421.02 |
| 0.7000 | 0.8916 / 0.8927 / 0.7093 | 27% (25) / 28% (25) / **3%** (17) | 2516.98 |
| 0.7000 | 0.8896 / 0.8907 / 0.7079 | 27% (25) / 27% (25) / **2%** (14) | 2538.10 |
| 0.8000 | 1.0226 / 1.0238 / 0.8008 | 28% (25) / 28% (25) / **2%** (13) | 2635.46 |
| 0.9000 | 1.1786 / 1.1800 / 0.9087 | 31% (25) / 31% (25) / **2%** (16) | 2752.36 |
| 1.3000 | 1.7555 / 1.7578 / 1.3035 | 35% (25) / 35% (25) / **1%** (13) | 3317.51 |
| 1.5000 | 2.0670 / 2.0699 / 1.5178 | 38% (25) / 38% (25) / **2%** (17) | 3650.17 |
| 1.5000 | 2.0717 / 2.0747 / 1.5211 | 38% (25) / 38% (25) / **2%** (17) | 3659.76 |
| 1.9000 | 2.6322 / 2.6368 / 1.9136 | 38% (25) / 39% (25) / **2%** (17) | 4335.98 |
| 2.0000 | 2.7602 / 2.7652 / 2.0048 | 38% (25) / 38% (25) / **2%** (11) | 4510.40 |
| 2.5000 | 3.4627 / 3.4713 / 2.5169 | 38% (25) / 39% (25) / **2%** (17) | 5469.20 |
| 2.9000 | 3.9971 / 4.0101 / 2.9205 | 38% (25) / 38% (25) / **3%** (13) | 6268.18 |
| 3.6000 | 4.9248 / 4.9520 / 3.6504 | 37% (25) / 38% (25) / **3%** (17) | 7726.47 |
| 4.1000 | 5.6207 / 5.6704 / 4.2264 | 37% (25) / 38% (25) / **4%** (19) | 8738.56 |
| 6.0000 | 7.6885 / 8.0690 / 6.2425 | 28% (25) / 34% (25) / **8%** (12) | 12757.00 |
| 7.0000 | 9.6941 / $\infty$ / $\infty$ | 38% (25) / NAN (25) / NAN (25) | 25016.96 |

#### 5 Conditioning of Branch Length Estimation

We now examine the conditioning of ML branch length estimation. Note that the MLE for retroelement insertions (MLE-RI) is given by Equation 15 in the Appendix; this is equivalent to using Newton's method to solve

$$g^{RI}(\tau) = \frac{\frac{1}{3}e^{-\tau} + \tau}{e^{-\tau} + \tau} - \frac{z_1}{n} \tag{8}$$

for $\tau$. Similarly, the MLE for gene trees (MLE-GT) can be found by solving

$$g^{GT}(\tau) = 1 - \frac{2}{3}e^{-\tau} - \frac{z_1}{n} \tag{9}$$

for $\tau$ (Theorem 2 in Sayyari and Mirarab 2016). The absolute condition number for a root finding problem can be found by evaluating

$$\left(\left|\frac{\mathrm{d}g(\tau)}{\mathrm{d}\tau}\right|\right)^{-1}$$

at the root $\tau^*$. Therefore, we can use the derivatives of Equation 1

$$\begin{aligned}
\frac{\mathrm{d}g^{RI}(\tau)}{\mathrm{d}\tau} &= \frac{(1 - \frac{1}{3}e^{-\tau})(e^{-\tau} + \tau) - (\frac{1}{3}e^{-\tau} + \tau)(1 - e^{-\tau})}{(e^{-\tau} + \tau)^2} \\
&= \frac{e^{-\tau} + \tau - \frac{1}{3}e^{-2\tau} - \frac{1}{3}\tau e^{-\tau} - \frac{1}{3}e^{-\tau} + \frac{1}{3}e^{-2\tau} - \tau + \tau e^{-\tau}}{(e^{-\tau} + \tau)^2} \\
&= \frac{\frac{2}{3}e^{-\tau}(\tau + 1)}{(e^{-\tau} + \tau)^2}
\end{aligned}$$

and Equation 2

$$\frac{\mathrm{d}g^{GT}(\tau)}{\mathrm{d}\tau} = \frac{2}{3}e^{-\tau}$$

to examine the conditioning of ML branch length estimation for RI and gene tree data sets, respectively. In the table below, we show the absolute condition number of some values of $\tau^*$. Clearly, the conditioning of $\tau^*$ increases, indicating the problem is increasingly ill conditioned as $\tau^*$ increases. This is expected looking at Figure 2 in the main text, because for large values of $\tau^*$, a small perturbation in the quartet probability (estimated as $\frac{z_1}{n}$) results in a large change in $\tau^*$.

Table S2: **Condition number for ML branch length estimation.**

| $\tau^*$ | RIs | Gene Trees |
|---|---|---|
| 0.01 | 1.50 | 1.515 |
| 0.1 | 1.522 | 1.658 |
| 0.25 | 1.631 | 1.926 |
| 0.5 | 2.019 | 2.473 |
| 0.75 | 2.711 | 3.176 |
| 1 | 3.815 | 4.077 |
| 1.5 | 7.984 | 6.723 |
| 1.88 | 14.102 | 9.830 |
| 2 | 16.846 | 11.084 |
| 2.5 | 34.810 | 18.274 |
| 3 | 70.057 | 30.128 |
| 3.5 | 137.565 | 49.673 |

#### 6 Using local PP to control false positive branches

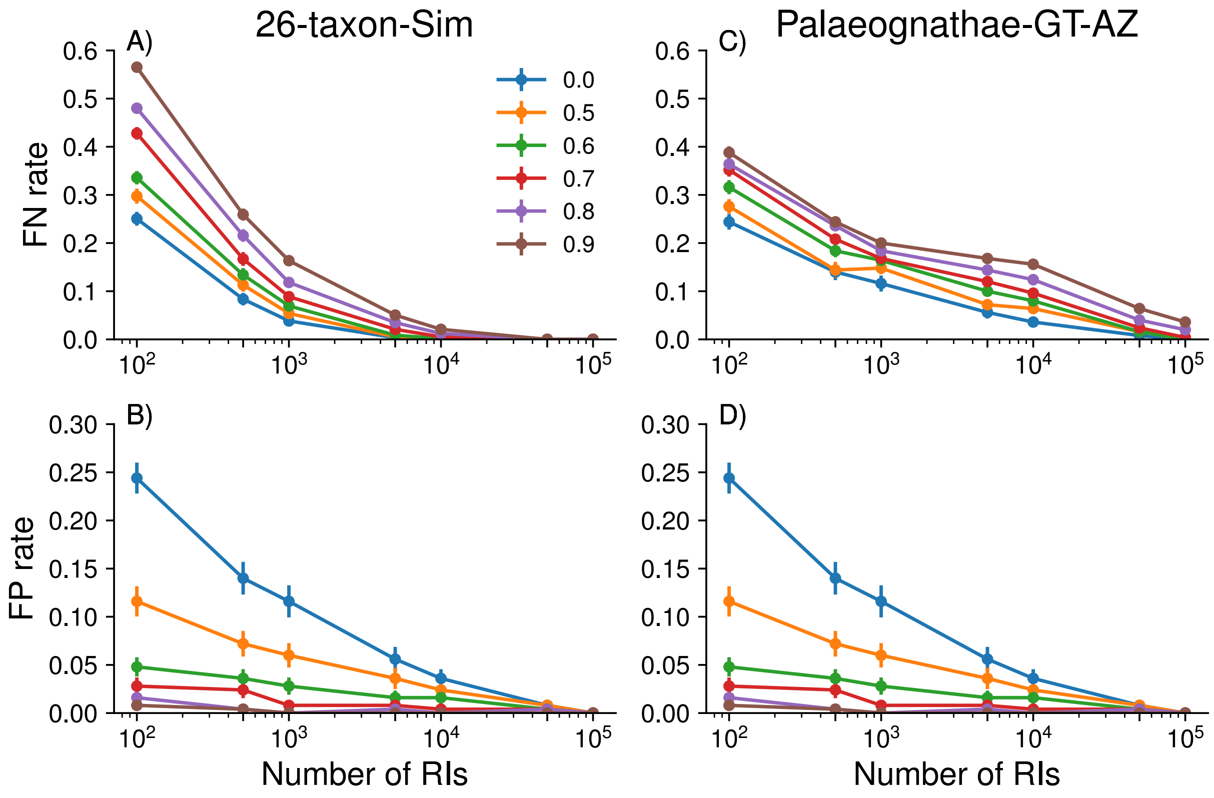

Figure S1: **Impact of collapsing branches with low support on species tree error.** Branches in the ASTRAL_BP tree are collapsed if they branch support (local posterior probability) is less than some threshold (indicated in the legend by color). Species tree error is measured as the false negative (FN) rate (i.e. the number of branches in the true species tree that are missing from the estimated tree, divided by the number of branches in the true species tree) and the false positive rate (i.e. the number of branches in the estimated species tree that are missing from the true tree, divided by the number of branches in the true tree). The top two subplots (A and C) show FN rate, and the bottom two subplots (B and D) show FP rate. These values are shown over 25 replicate data sets; dots are means, and bars are standard errors. The number of parsimony informative retroelement insertions (RIs) in these simulated data sets varies from 100 to 100,000 as shown on the $x$-axis. Subplots A–B and C–D show results for RIs simulated from the 26-taxa and Palaeognathae-GT-AZ model species trees, respectively.

### 7 ASTRAL_BP Analyses for Palaeognathae

Table S3: **Branch lengths for Palaeognathae-GT-AZ and Palaeognathae-RI species trees.** The Palaeognathae-GT-AZ species tree was estimated by Cloutier et al. (2019) from the set of 20 850 DNA-sequence-based gene trees. The Palaeognathae-RI species tree was estimated by Springer et al. (2020) from the 4 301 parsimony-informative RIs assembled by Cloutier et al. (2019). All branches in the Palaeognathae-RI tree had support (local PP) of 1.0, with the exception of Kiwi, Cassowary, & Emu, which had a local PP of 0.89. However, when the the effective number ($EN$) of RIs around a branch is low, the estimated length and local PP should be interpreted cautiously.

| Clade | Palaeognathae-GT-AZ | Palaeognathae-RI | | |
|---|---|---|---|---|
| | Branch Length MAP-GT | Branch Length MLE-RI | Quartet Support | EN |
| Spotted kiwi | 1.0655 | 2.8358 | 0.9865 / 0.0000 / 0.0135 | 148.00 |
| All kiwi | 3.0120 | $\infty$ | 1.0000 / 0.0000 / 0.0000 | 819.54 |
| Cassowary & emu | 1.9547 | $\infty$ | 1.0000 / 0.0000 / 0.0000 | 59.46 |
| Kiwi, cassowary, emu | 0.0532 | 0.2587 | 0.5006 / 0.2402 / 0.2592 | 26.23 |
| All rhea | 4.1674 | $\infty$ | 1.0000 / 0.0000 / 0.0000 | 2325.47 |
| Kiwi, cassowary, emu, rhea | 0.0194 | 0.8938 | 0.7820 / 0.1429 / 0.0752 | 13.30 |
| Chilean & elegant crested tinamou | 0.3091 | 1.3408 | 0.8912 / 0.0272 / 0.0816 | 73.50 |
| Thicket & white-throated tinamou | 1.1514 | 4.0119 | 0.9970 / 0.0030 / 0.0000 | 334.00 |
| All tinamou | 3.0872 | 5.4162 | 0.9994 / 0.0000 / 0.0006 | 522.79 |
| All but chicken & ostrich | 0.3874 | $\infty$ | 1.0000 / 0.0000 / 0.0000 | 18.00 |

Table S4: **Branch information for species trees estimated from 1 000 or 5 000 parsimony-informative RIs simulated from Palaeognathae-GT-AZ model species tree.** We report the estimated (i.e. MLE-RI) branch length, the local PP, and the $EN$, averaged over all replicates that recovered the branch. The number of replicates recovering the branch is also indicated.

| Clade | True Length | Estimated (i.e. MLE-RI) Length | local PP | $EN$ | # Replicates |
|---|---|---|---|---|---|
| *Data sets with 1 000 RIs* | | | | | |
| Spotted kiwi | 1.0655 | $1.0879 \pm 0.1438$ | $1.00 \pm 0.00$ | $89.84 \pm 8.33$ | 25 |
| All kiwi | 3.0120 | $3.4374 \pm 0.6732$ | $1.00 \pm 0.00$ | $197.27 \pm 12.00$ | 25 |
| Emu & cassowary | 1.9547 | $1.9612 \pm 0.1890$ | $1.00 \pm 0.00$ | $134.20 \pm 10.51$ | 25 |
| Kiwi, emu, cassowary | 0.0532 | $0.0880 \pm 0.0459$ | $0.62 \pm 0.16$ | $63.63 \pm 4.27$ | 13 |
| All rhea | 4.1674 | $5.2487 \pm 2.1562$ | $1.00 \pm 0.00$ | $262.06 \pm 12.08$ | 25 |
| Kiwi, emu, cassowary, rhea | 0.0194 | $0.0741 \pm 0.0485$ | $0.58 \pm 0.18$ | $65.10 \pm 5.45$ | 8 |
| Chilean & elegant crested tinamou | 0.3091 | $0.3551 \pm 0.0886$ | $1.00 \pm 0.01$ | $69.88 \pm 6.92$ | 25 |
| Thicket & white-throated tinamou | 1.1514 | $1.1556 \pm 0.2020$ | $1.00 \pm 0.00$ | $92.54 \pm 7.90$ | 25 |
| All tinamou | 3.0872 | $3.3275 \pm 0.6096$ | $1.00 \pm 0.00$ | $199.28 \pm 11.32$ | 25 |
| All but ostrich and chicken | 0.3874 | $0.3565 \pm 0.1264$ | $0.98 \pm 0.03$ | $68.49 \pm 6.02$ | 25 |
| *Data sets with 5 000 RIs* | | | | | |
| Spotted kiwi | 1.0655 | $1.0710 \pm 0.0811$ | $1.00 \pm 0.00$ | $447.04 \pm 15.55$ | 25 |
| All kiwi | 3.0120 | $3.0928 \pm 0.2305$ | $1.00 \pm 0.00$ | $977.75 \pm 23.41$ | 25 |
| Emu & cassowary | 1.9547 | $1.9708 \pm 0.1071$ | $1.00 \pm 0.00$ | $672.95 \pm 20.33$ | 25 |
| Kiwi, emu, cassowary | 0.0532 | $0.0625 \pm 0.0228$ | $0.80 \pm 0.13$ | $322.49 \pm 11.39$ | 23 |
| All rhea | 4.1674 | $4.2161 \pm 0.2535$ | $1.00 \pm 0.00$ | $1337.77 \pm 32.06$ | 25 |
| Kiwi, emu, cassowary, rhea | 0.0194 | $0.0372 \pm 0.0198$ | $0.60 \pm 0.15$ | $313.97 \pm 14.17$ | 13 |
| Chilean & elegant crested tinamou | 0.3091 | $0.3217 \pm 0.0338$ | $1.00 \pm 0.00$ | $339.69 \pm 15.81$ | 25 |
| Thicket & white-throated tinamou | 1.1514 | $1.1531 \pm 0.0588$ | $1.00 \pm 0.00$ | $470.89 \pm 23.10$ | 25 |
| All tinamou | 3.0872 | $3.1928 \pm 0.2265$ | $1.00 \pm 0.00$ | $998.28 \pm 27.17$ | 25 |
| All but ostrich & chicken | 0.3874 | $0.3785 \pm 0.0492$ | $1.00 \pm 0.00$ | $338.72 \pm 14.26$ | 25 |

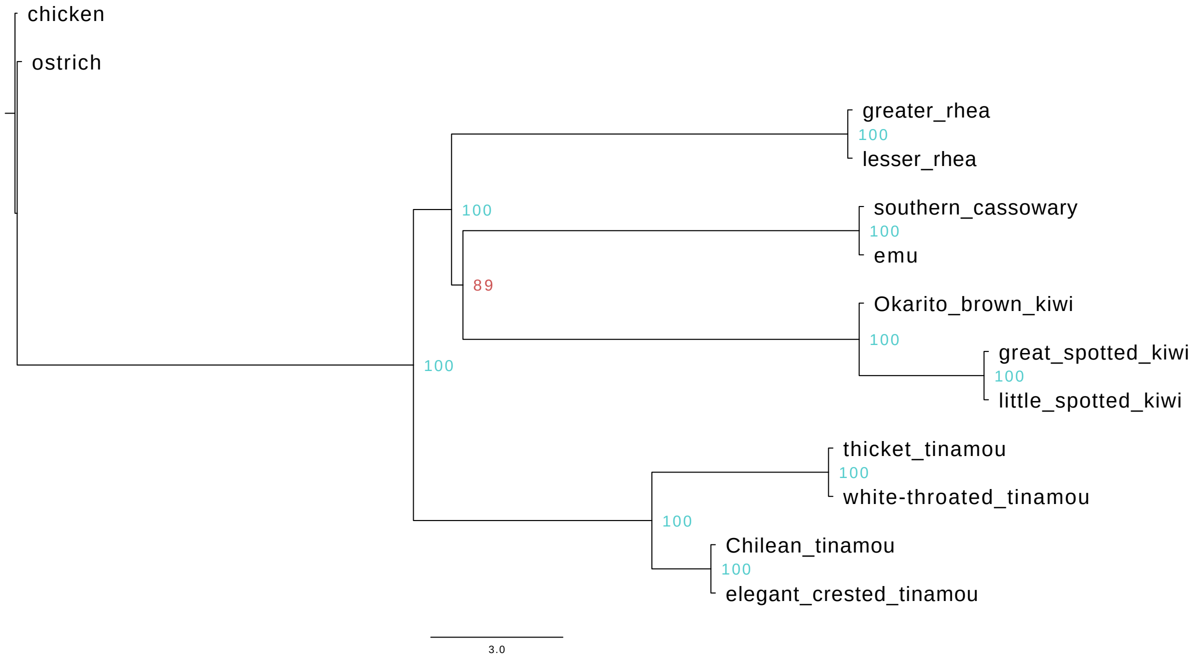

Figure S2: **Palaeognathae-RI tree.** For Palaeognathae data set with 4 301 RIs, ASTRAL_BP recovered the tree shown above. The *branch separating kiwi and emu+cassowary* from the remainder of the taxa has support (local PP) of 0.8914 and length (MLE-RI) of 0.2587 CUs. The *EN* around this branch is 26.2333, and the three quartet topologies around this branch have (normalized) frequencies of 0.5006, 0.2402, and 0.2592. The *branch separating kiwi, emu+cassowary, and rhea* from the remainder of the taxa has support (local PP) of 0.9983 and length (MLE-RI) of 0.8657 CUs. The *EN* around this branch is 13.3000, and the three quartet topologies around this branch have (normalized) frequencies of 0.7820, 0.0752, and 0.1429. Importantly, ASTRAL gave warnings for trusting the local PP due to the low EN around these branches.

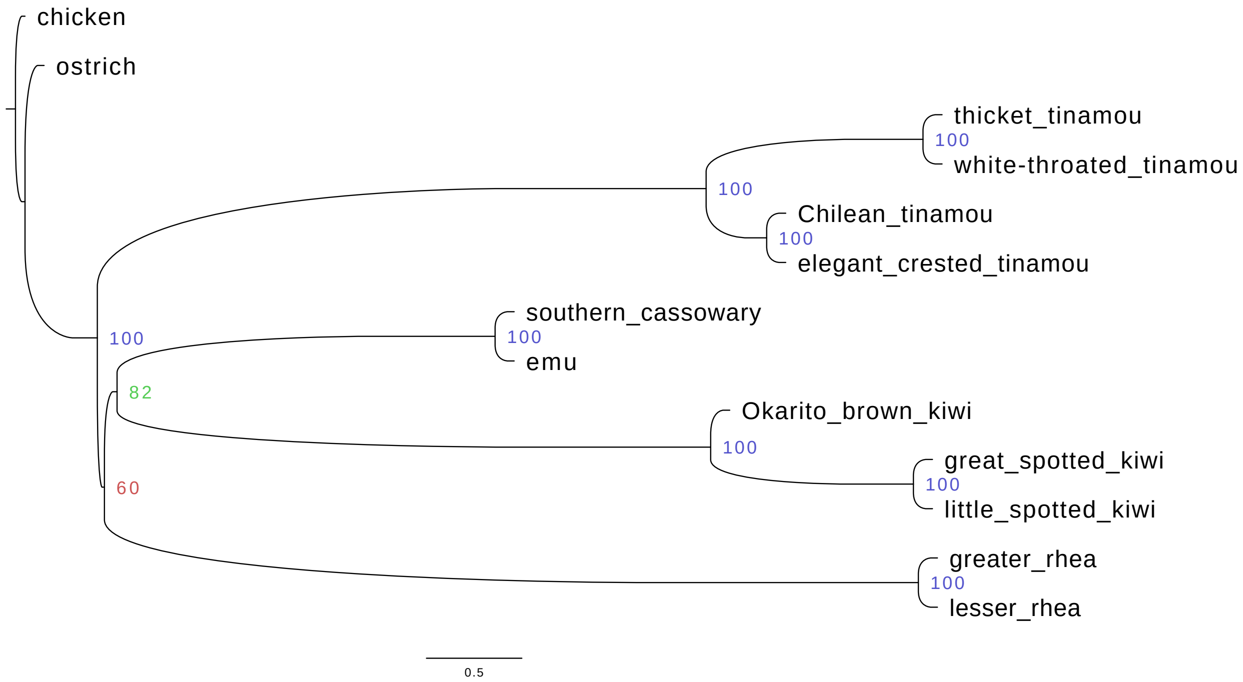

Figure S3: **ASTRAL_BP tree # 1 for Palaeognathae-GT-AZ data set with 5 000 RIs.** For the Palaeognathae-GT-AZ simulated data sets with 5 000 RIs, ASTRAL_BP recovered the correct topology for 13/25 replicates (# 3, 5, 6, 7, 9, 10, 11, 13, 14, 15, 16, 21, 23). The tree (shown above) is drawn with branch lengths (CUs) and support values (local PP multiplied by 100), averaged across the 13 replicates; similarly, the following numbers are mean ± standard deviation. The *(correct) branch separating kiwi and emu+cassowary* from the remainder of the taxa has support (local PP) of $0.8159 \pm 0.1262$ and length (MLE-RI) of $0.0662 \pm 0.0232$ CUs. The $EN$ around this branch is $322.0566 \pm 11.9794$, and the three quartet topologies around this branch have (normalized) frequencies of $0.3773 \pm 0.0153$, $0.3144 \pm 0.0166$, and $0.3082 \pm 0.0200$. The *(correct) branch separating kiwi, emu+cassowary, and rhea* from the remainder of the taxa has support (local PP) of $0.6015 \pm 0.1474$ and length (MLE-RI) of $0.0372 \pm 0.0198$ CUs. The $EN$ around this branch is $313.9673 \pm 14.1682$, and the three quartet topologies around this branch have (normalized) frequencies of $0.3581 \pm 0.0132$, $0.3127 \pm 0.0180$, and $0.3292 \pm 0.0145$. Note that all 13 of the estimated species trees are in the anomaly zone.

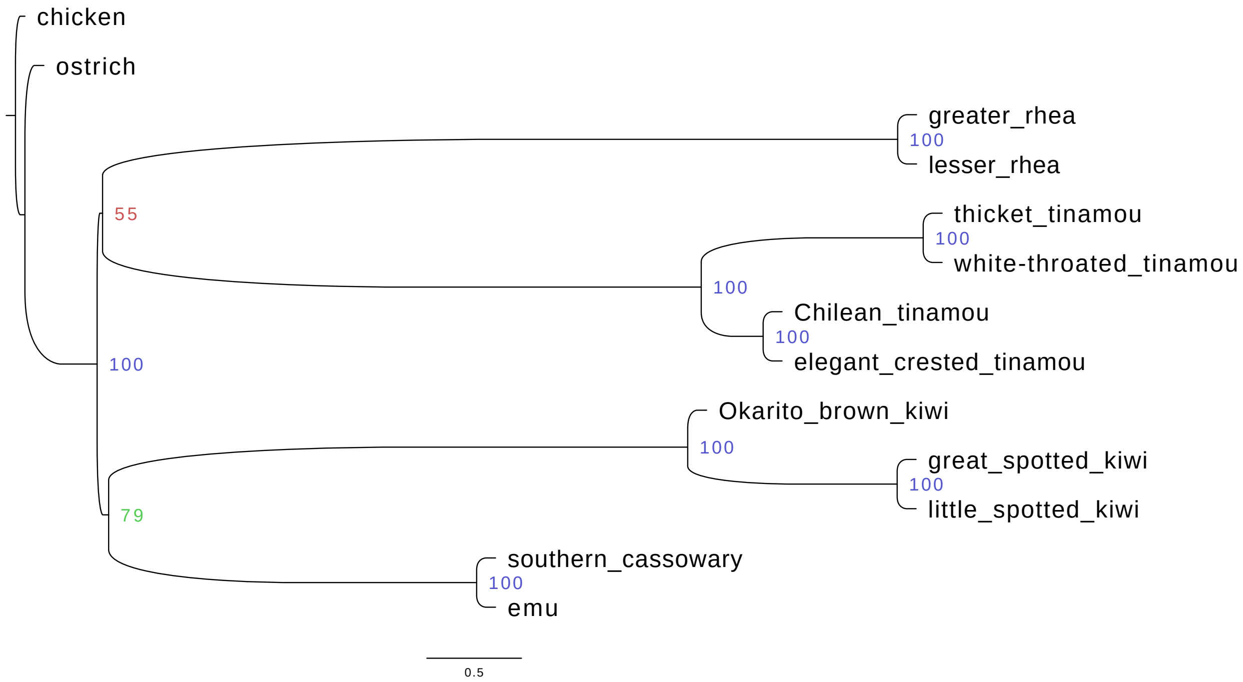

Figure S4: **ASTRAL_BP tree # 2 for Palaeognathae-GT-AZ data set with 5 000 RIs.** For the Palaeognathae-GT-AZ simulated data sets with 5 000 RIs, ASTRAL_BP recovered the same incorrect topology for 8/25 replicates (# 1, 4, 8, 12, 18, 20, 22, 24). The tree (shown above) is drawn with branch lengths (CUs) and support values (local PP multiplied by 100) averaged across the 8 replicates; similarly, the following numbers are mean ± standard deviation. The *(correct) branch separating kiwi and emu+cassowary* from the remainder of the taxa has support (local PP) of $0.7925 \pm 0.1496$ and length (MLE-RI) of $0.0610 \pm 0.0228$ CUs. The $EN$ around this branch is $322.7014 \pm 10.7719$, and the three quartet topologies around this branch have (normalized) frequencies of $0.3739 \pm 0.0151$, $0.3152 \pm 0.0167$, and $0.3108 \pm 0.0118$. The *(incorrect) branch separating rhea and tinamou* from the remainder of the taxa has support (local PP) of $0.5526 \pm 0.1558$ and length (MLE-RI) of $0.0288 \pm 0.0184$ CUs. The $EN$ around this branch is $325.5531 \pm 10.9030$, and the three quartet topologies around this branch have (normalized) frequencies of $0.3525 \pm 0.0122$, $0.3207 \pm 0.0120$, and $0.3268 \pm 0.0122$.

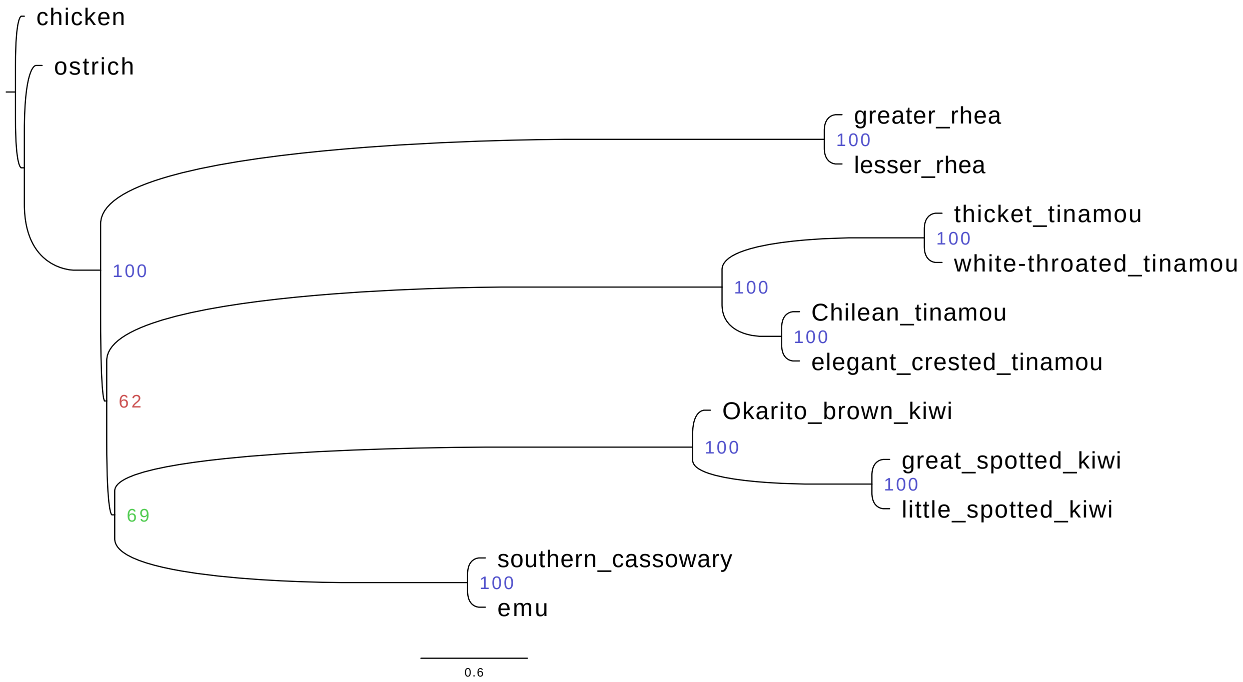

Figure S5: **ASTRAL_BP tree # 3 for Palaeognathae-GT-AZ data set with 5 000 RIs.** For the Palaeognathae-GT-AZ simulated data sets with 5 000 RIs, ASTRAL_BP recovered the same incorrect topology for 2/25 replicates (# 19 and 25). The tree (shown above) is drawn with branch lengths (CUs) and support values (local PP multiplied by 100) averaged across the 2 replicates; similarly, the following numbers are mean ± standard deviation. The *(correct) branch separating kiwi and emu+cassowary* from the remainder of the taxa has support (local PP) of $0.6905 \pm 0.0236$ and length (MLE-RI) of $0.0450 \pm 0.0006$ CUs. The $EN$ around this branch is $324.4219 \pm 9.4740$, and the three quartet topologies around this branch have (normalized) frequencies of $0.3633 \pm 0.0004$, $0.3064 \pm 0.0095$, and $0.3302 \pm 0.0099$. The *(incorrect) branch separating kiwi, emu+cassowary, and tinamou* from the remainder of the taxa has support (local PP) of $0.6182 \pm 0.0663$ and length (MLE-RI) of $0.0344 \pm 0.0073$ CUs. The $EN$ around this branch is $335.4000 \pm 0.8250$, and the three quartet topologies around this branch have (normalized) frequencies of $0.3563 \pm 0.0049$, $0.3230 \pm 0.0048$, and $0.3208 \pm 0.0097$. Note that both of these estimated species trees are in the anomaly zone.

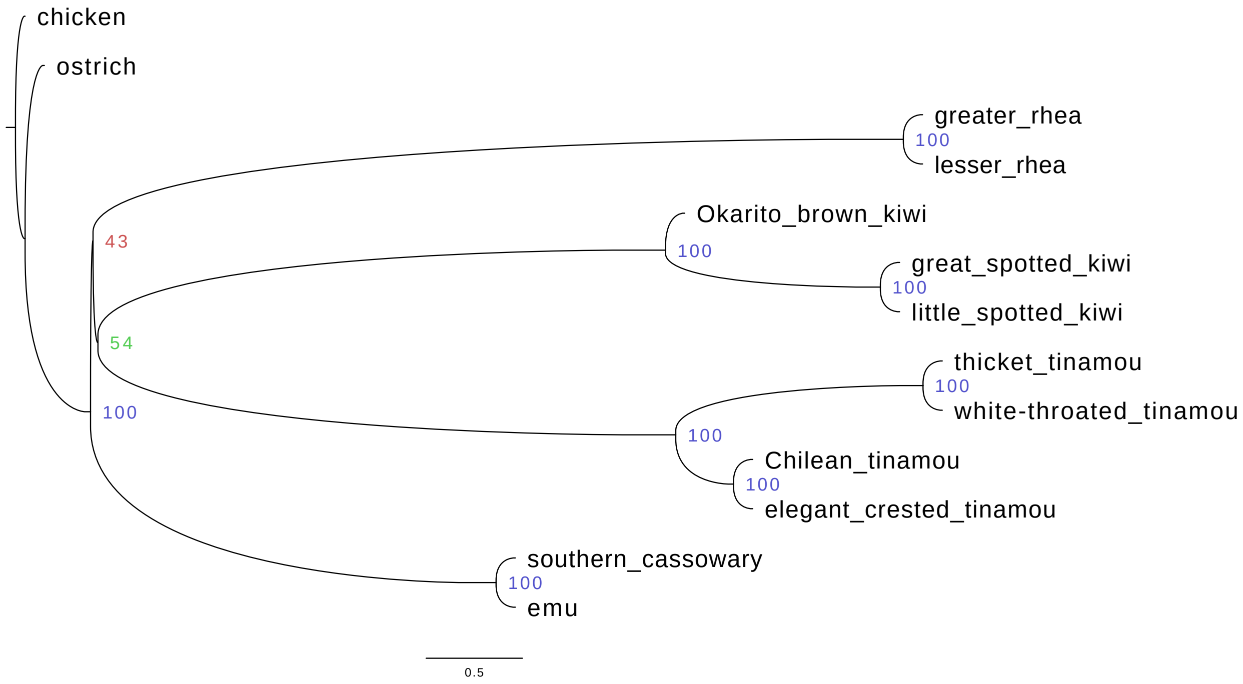

Figure S6: **ASTRAL_BP tree # 4 for Palaeognathae-GT-AZ data set with 5 000 RIs.** For the Palaeognathae-GT-AZ simulated data sets with 5 000 RIs, ASTRAL_BP recovered a unique incorrect topology for replicate # 2. The *(incorrect) branch separating kiwi and tinamou* from the remainder of the taxa has support (local PP) of 0.5446 and length (MLE-RI) of 0.0256 CUs. The $EN$ around this branch is 343.3125, and the three quartet topologies around this branch have (normalized) frequencies of 0.3504, 0.3274, and 0.3222. The *(incorrect) branch separating kiwi, tinamou, and rhea* from the remainder of the taxa has support (local PP) of 0.4320 and length (MLE-RI) of 0.0129 CUs. The $EN$ around this branch is 341.3214, and the three quartet topologies around this branch have (normalized) frequencies of 0.3419, 0.3272, and 0.3309. Note that the estimated species trees is in the anomaly zone.

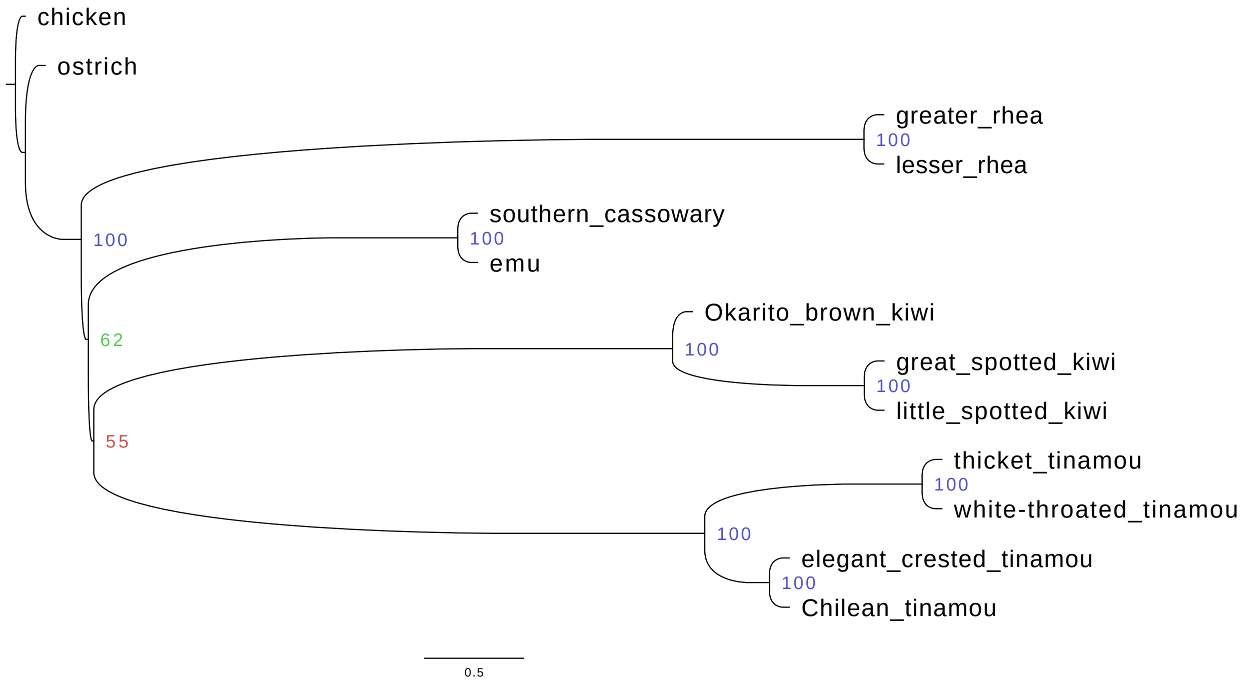

Figure S7: **ASTRAL_BP tree # 5 for Palaeognathae-GT-AZ data set with 5 000 RIs.** For the Palaeognathae-GT-AZ simulated data sets with 5 000 RIs, ASTRAL_BP recovered a unique incorrect topology for replicate # 17. The *(incorrect) branch separating kiwi and tinamou* from the remainder of the taxa has support (local PP) of 0.5455 and length (MLE-RI) of 0.0276 CUs. The $EN$ around this branch is 340.4375, and the three quartet topologies around this branch have (normalized) frequencies of 0.3517, 0.3131, and 0.3352. The *(incorrect) branch separating kiwi, tinamou, and emu+cassowary* from the remainder of the taxa has support (local PP) of 0.6205 and length (MLE-RI) of 0.0355 CUs. The $EN$ around this branch is 341.0714, and the three quartet topologies around this branch have (normalized) frequencies of 0.3570, 0.3332, and 0.3098. Note that the estimated species trees is in the anomaly zone.

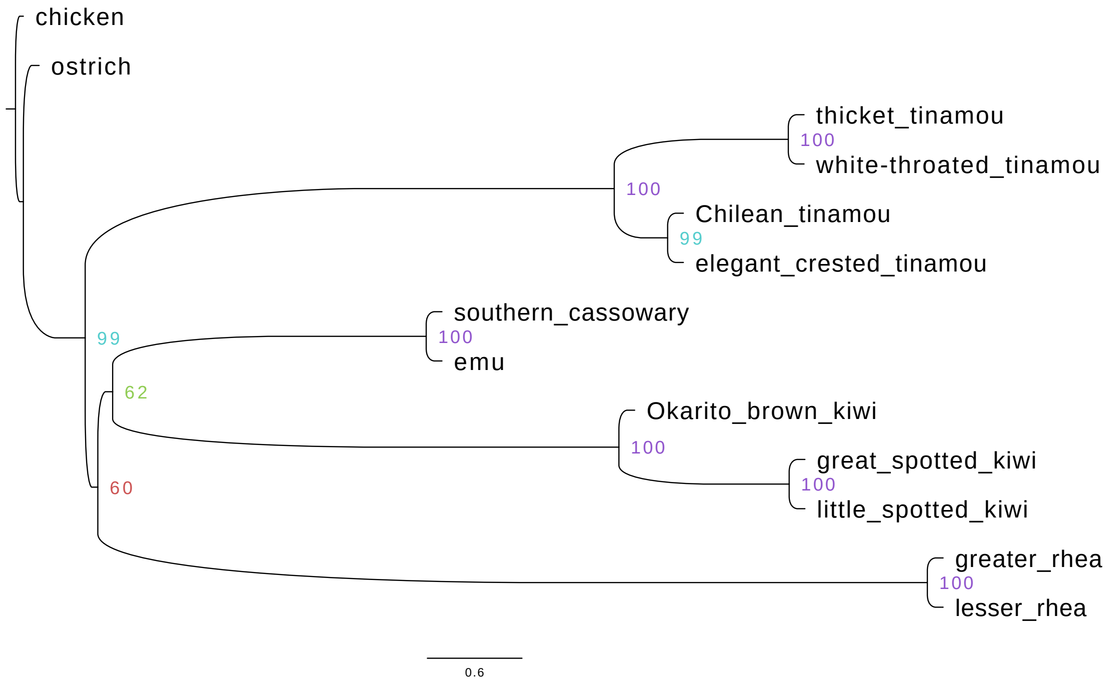

Figure S8: **ASTRAL_BP tree # 1 for Palaeognathae-GT-AZ data set with 1 000 RIs.** For the Palaeognathae-GT-AZ simulated data sets with 1 000 RIs, ASTRAL_BP recovered the correct topology for 7/25 replicates (# 3, 4, 7, 14, 16, 20, 24). The tree (shown above) is drawn with branch lengths (CUs) and support values (local PP multiplied by 100) averaged across the 7 replicates; similarly, the following numbers are mean $\pm$ standard deviation. The *(correct) branch separating kiwi and emu+cassowary* from the remainder of the taxa has support (local PP) of $0.6248 \pm 0.1850$ and length (MLE-RI) of $0.0940+/-0.0563$ CUs. The $EN$ around this branch is $62.6310 \pm 3.9461$, and the three quartet topologies around this branch have (normalized) frequencies of $0.3954 \pm 0.0366$, $0.3133 \pm 0.0395$, and $0.2913 \pm 0.0368$. The *(correct) branch separating kiwi, emu+cassowary, and rhea* from the remainder of the taxa has support (local PP) of $0.6036 \pm 0.1832$ and length (MLE-RI) of $0.0802 \pm 0.0489$ CUs. The $EN$ around this branch is $63.8429 \pm 4.6208$, and the three quartet topologies around this branch have (normalized) frequencies of $0.3865 \pm 0.0323$, $0.3014 \pm 0.0314$, and $0.3121 \pm 0.0409$. Note that 4/7 estimated species trees replicates (# 4, 7, 16, 20) are in the anomaly zone.

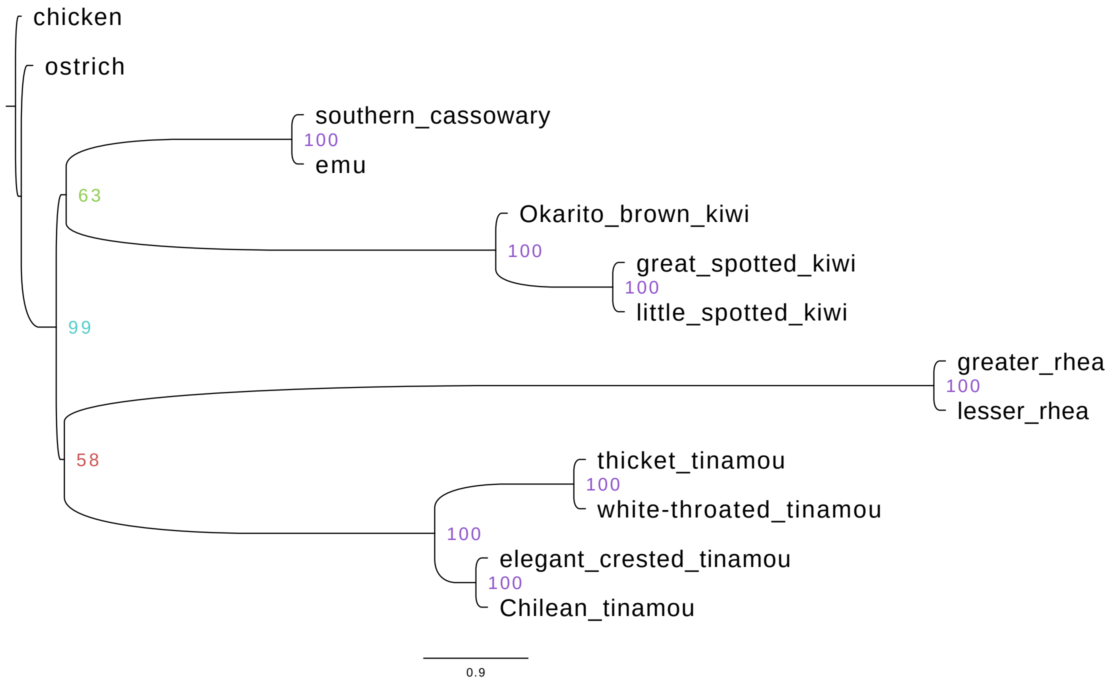

Figure S9: **ASTRAL_BP tree # 2 for Palaeognathae-GT-AZ data set with 1 000 RIs.** For the Palaeognathae-GT-AZ simulated data sets with 1 000 RIs, ASTRAL_BP recovered the same <u>incorrect</u> topology for 4/25 replicates (# 5, 11, 21, 23). The tree (shown above) is drawn with branch lengths (CUs) and support values (local PP multiplied by 100) averaged across the 4 replicates; similarly, the following numbers are mean ± standard deviation. The *(correct) branch separating kiwi and emu+cassowary* from the remainder of the taxa has support (local PP) of $0.6256 \pm 0.1355$ and length (MLE-RI) of $0.0858 \pm 0.0318$ CUs. The $EN$ around this branch is $64.9444 \pm 4.5887$, and the three quartet topologies around this branch have (normalized) frequencies of $0.3903 \pm 0.0210$, $0.3078 \pm 0.0276$, and $0.3020 \pm 0.0368$. The *(incorrect) branch separating rhea and tinamou* from the remainder of the taxa has support (local PP) of $0.5753 \pm 0.0973$ and length (MLE-RI) of $0.0702 \pm 0.0262$ CUs. The $EN$ around this branch is $64.4125 \pm 6.6018$, and the three quartet topologies around this branch have (normalized) frequencies of $0.3800 \pm 0.0174$, $0.3221 \pm 0.0186$, and $0.2979 \pm 0.0214$.

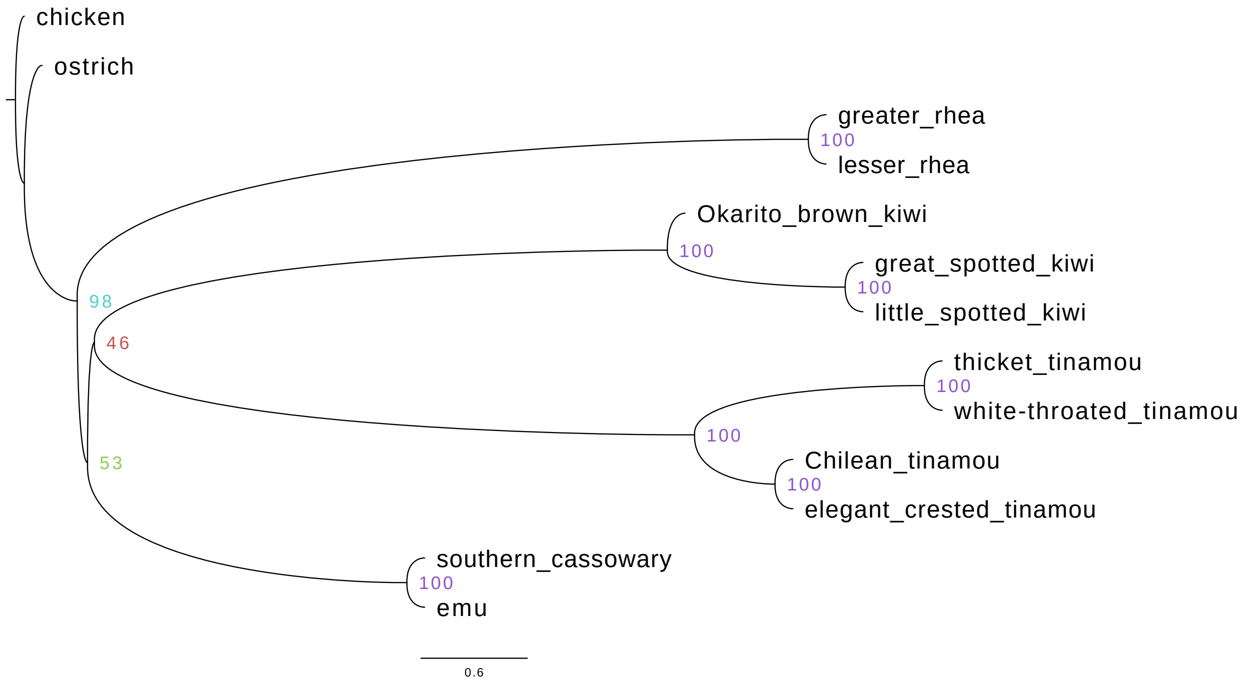

Figure S10: **ASTRAL_BP tree # 3 for Palaeognathae-GT-AZ data set with 1 000 RIs.** For the Palaeognathae-GT-AZ simulated data sets with 1 000 RIs, ASTRAL_BP recovered the same incorrect topology for 3/25 replicates (# 9, 17, 22). The tree (shown above) is drawn with branch lengths (CUs) and support values (local PP multiplied by 100) averaged across the 3 replicates; similarly, the following numbers are mean $\pm$ standard deviation. The *(incorrect) branch separating kiwi and tinamou* from the remainder of the taxa has support (local PP) of $0.4646 \pm 0.0476$ and length (MLE-RI) of $0.0390 \pm 0.0116$ CUs. The $EN$ around this branch is $62.6528 \pm 5.7339$, and the three quartet topologies around this branch have (normalized) frequencies of $0.3593 \pm 0.0077$, $0.3267 \pm 0.0030$, and $0.3140 \pm 0.0107$. The *(incorrect) branch separating kiwi, tinamou, and emu+cassowary* from the remainder of the taxa has support (local PP) of $0.5276 \pm 0.0591$ and length (MLE-RI) of $0.0583 \pm 0.0147$ CUs. The $EN$ around this branch is $61.6190 \pm 4.8926$, and the three quartet topologies around this branch have (normalized) frequencies of $0.3721 \pm 0.0097$, $0.3264 \pm 0.0165$, and $0.3014 \pm 0.0068$. Note that all 3 of the estimated species trees are in the anomaly zone.

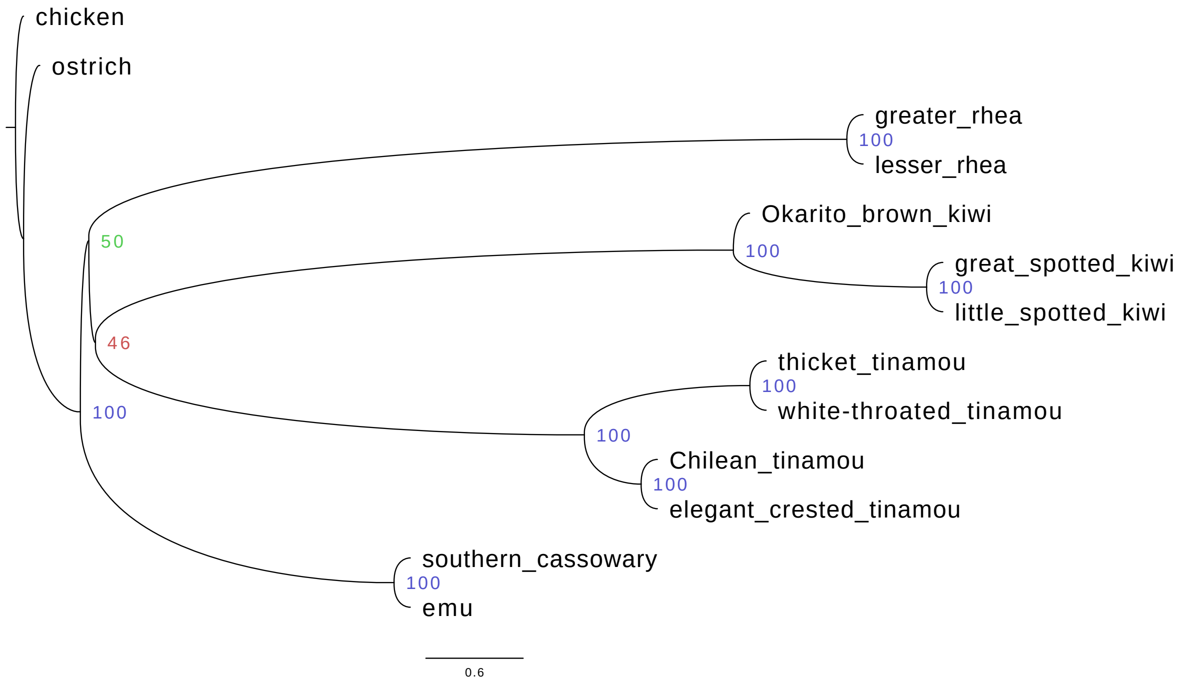

Figure S11: **ASTRAL_BP tree # 4 for Palaeognathae-GT-AZ data set with 1 000 RIs.** For the Palaeognathae-GT-AZ simulated data sets with 1 000 RIs, ASTRAL_BP recovered the same incorrect topology for 2/25 replicates (# 1 and 13). The tree (shown above) is drawn with branch lengths (CUs) and support values (local PP multiplied by 100) averaged across the 2 replicates; similarly, the following numbers are mean ± standard deviation. The *(incorrect) branch separating kiwi and tinamou* from the remainder of the taxa has support (local PP) of $0.4594 \pm 0.0229$ and length (MLE-RI) of $0.0409 \pm 0.0108$ CUs. The $EN$ around this branch is $68.3021 \pm 0.3229$, and the three quartet topologies around this branch have (normalized) frequencies of $0.3606 \pm 0.0072$, $0.3389 \pm 0.0129$, and $0.3005 \pm 0.0201$. The *(incorrect) branch separating kiwi, tinamou, and rhea* from the remainder of the taxa has support (local PP) of $0.4947 \pm 0.0436$ and length (MLE-RI) of $0.0522 \pm 0.0135$ CUs. The $EN$ around this branch is $64.8214 \pm 2.1071$, and the three quartet topologies around this branch have (normalized) frequencies of $0.3681 + / - 0.0089$, $0.2878 \pm 0.0094$, and $0.3441 \pm 0.0004$. Noth that both of these estimated species trees are in the anomaly zone.

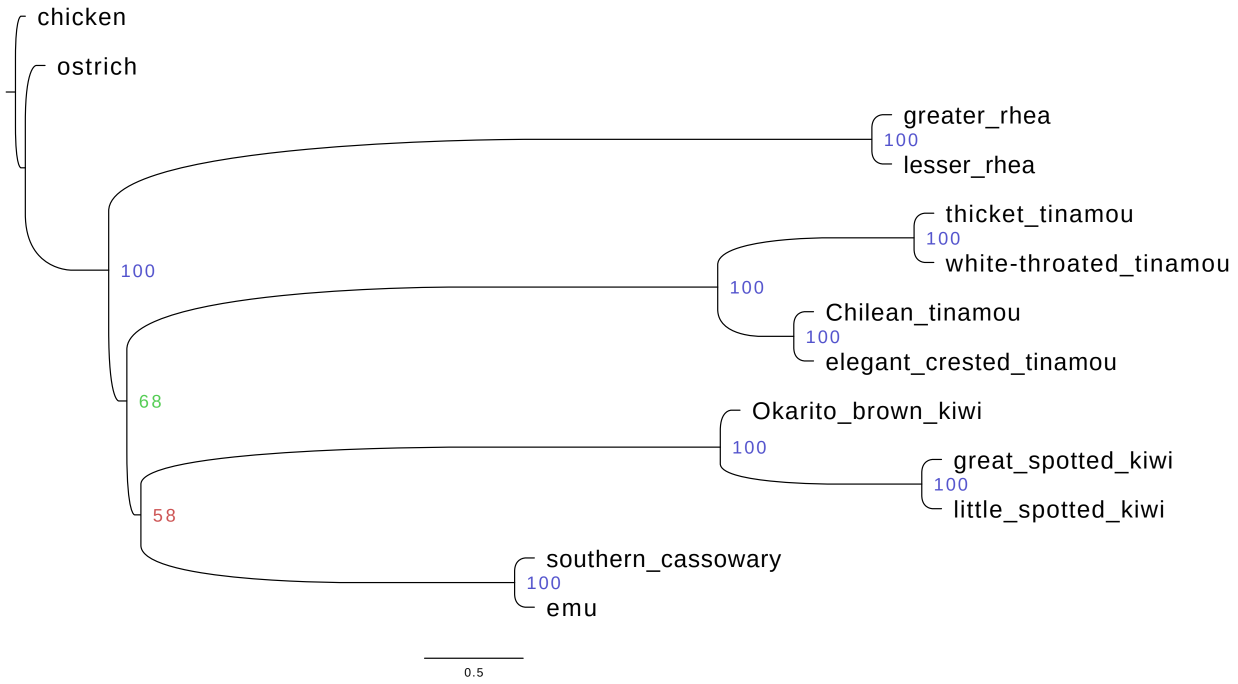

Figure S12: **ASTRAL_BP tree # 5 for Palaeognathae-GT-AZ data set with 1 000 RIs.** For the Palaeognathae-GT-AZ simulated data sets with 1 000 RIs, ASTRAL_BP recovered the same incorrect topology for 2/25 replicates (# 6 and 19). The tree (shown above) is drawn with branch lengths (CUs) and support values (local PP multiplied by 100) averaged across the 2 replicates; similarly, the following numbers are mean ± standard deviation. The *(correct) branch separating kiwi and emu+cassowary* from the remainder of the taxa has support (local PP) of $0.5761 \pm 0.0261$ and length (MLE-RI) of $0.0713 \pm 0.0120$ CUs. The $EN$ around this branch is $64.5104 \pm 3.7812$, and the three quartet topologies around this branch have (normalized) frequencies of $0.3807 \pm 0.0080$, $0.3319 \pm 0.0098$, and $0.2874 \pm 0.0177$. The *(incorrect) branch separating kiwi, emu+cassowary, and tinamou* from the remainder of the taxa has support (local PP) of $0.6753 \pm 0.0286$ and length (MLE-RI) of $0.0921 \pm 0.0122$ CUs. The $EN$ around this branch is $66.4250 + / - 7.9750$, and the three quartet topologies around this branch have (normalized) frequencies of $0.3945 \pm 0.0080$, $0.3108 \pm 0.0105$, and $0.2948 \pm 0.0025$. Note that both of the estimated species trees are in the anomaly zone.

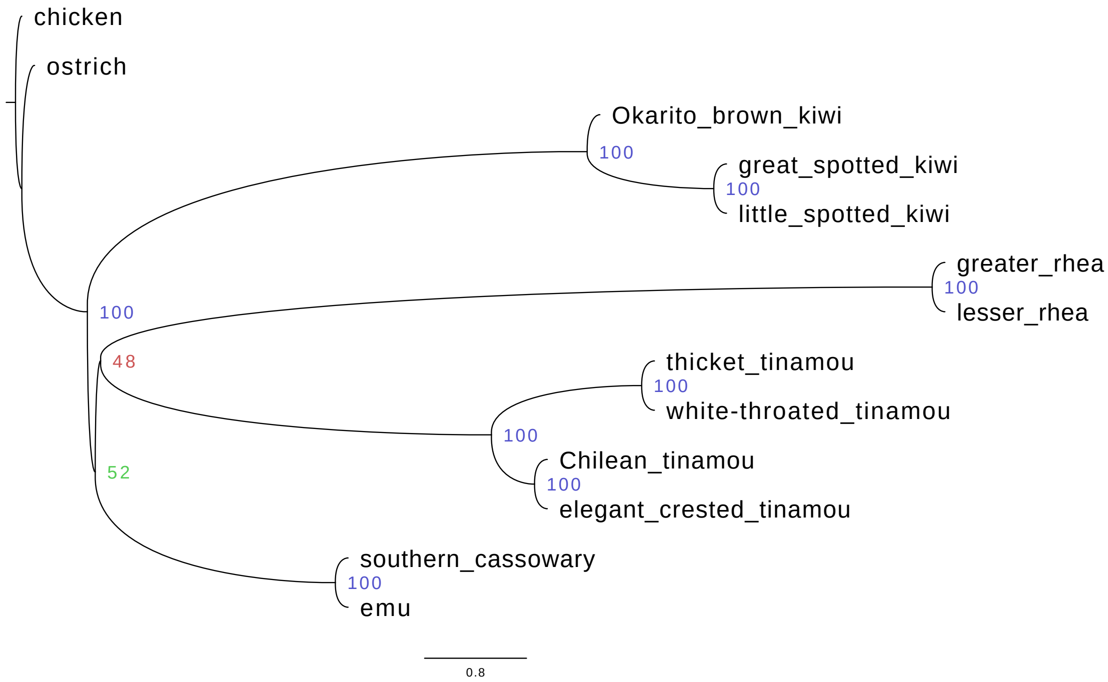

Figure S13: **ASTRAL_BP tree # 6 for Palaeognathae-GT-AZ data set with 1 000 RIs.** For the Palaeognathae-GT-AZ simulated data sets with 1 000 RIs, ASTRAL_BP recovered the same incorrect topology for 2/25 replicates (# 12 and 18). The tree (shown above) is drawn with branch lengths (CUs) and support values (local PP multiplied by 100) averaged across the 2 replicates; similarly, the following numbers are mean ± standard deviation. The *(incorrect) branch separating tinamou and rhea* from the remainder of the taxa has support (local PP) of $0.4764 \pm 0.0394$ and length (MLE-RI) of $0.0423 \pm 0.0097$ CUs. The $EN$ around this branch is $66.6250 \pm 1.3250$, and the three quartet topologies around this branch have (normalized) frequencies of $0.3615 \pm 0.0064$, $0.3073 \pm 0.0012$, and $0.3312 \pm 0.0077$. The *(incorrect) branch separating tinamou, rhea, and emu+cassowary* from the remainder of the taxa has support (local PP) of $0.5237 \pm 0.0876$ and length (MLE-RI) of $0.0618 \pm 0.0159$ CUs. The $EN$ around this branch is $69.8403 \pm 2.0625$, and the three quartet topologies around this branch have (normalized) frequencies of $0.3744 \pm 0.0105$, $0.3507 \pm 0.0120$, and $0.2749 \pm 0.0015$. Note that both of the estimated species trees are in the anomaly zone.

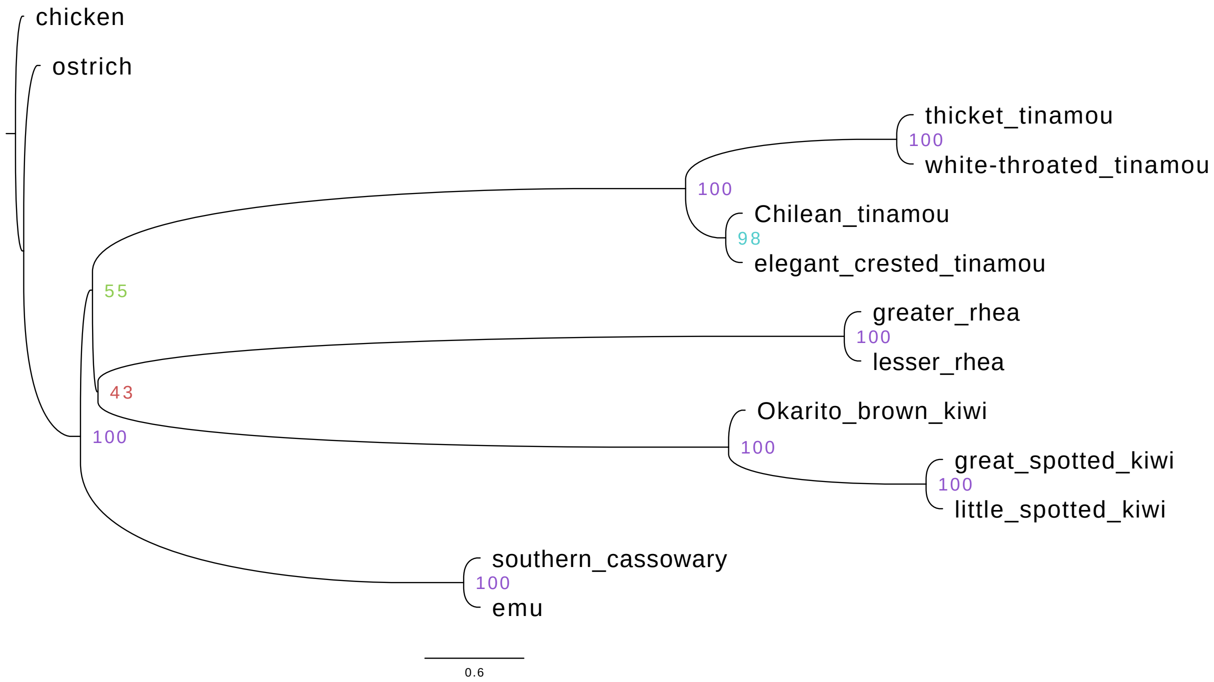

Figure S14: **ASTRAL_BP tree # 7 for Palaeognathae-GT-AZ data set with 1 000 RIs.** For the Palaeognathae-GT-AZ simulated data sets with 1 000 RIs, ASTRAL_BP recovered the same incorrect topology for 2/25 replicates (# 15 and 25). The tree (shown above) is drawn with branch lengths (CUs) and support values (local PP multiplied by 100) averaged across the 2 replicates; similarly, the following numbers are mean ± standard deviation. The *(incorrect) branch separating kiwi and rhea* from the remainder of the taxa has support (local PP) of $0.4300 \pm 0.0398$ and length (MLE-RI) of $0.0334 \pm 0.0135$ CUs. The $EN$ around this branch is $65.2708 \pm 9.2083$, and the three quartet topologies around this branch have (normalized) frequencies of $0.3556 \pm 0.0090$, $0.3013 \pm 0.0070$, and $0.3432 \pm 0.0020$. The *(incorrect) branch separating kiwi, rhea, and tinamou* from the remainder of the taxa has support (local PP) of $0.5535 \pm 0.0443$ and length (MLE-RI) of $0.0728 \pm 0.0232$ CUs. The $EN$ around this branch is $60.4750 \pm 13.5000$, and the three quartet topologies around this branch have (normalized) frequencies of $0.3817 \pm 0.0153$, $0.3228 \pm 0.0247$, and $0.2955 \pm 0.0401$. Note that both of the estimated species trees are in the anomaly zone.

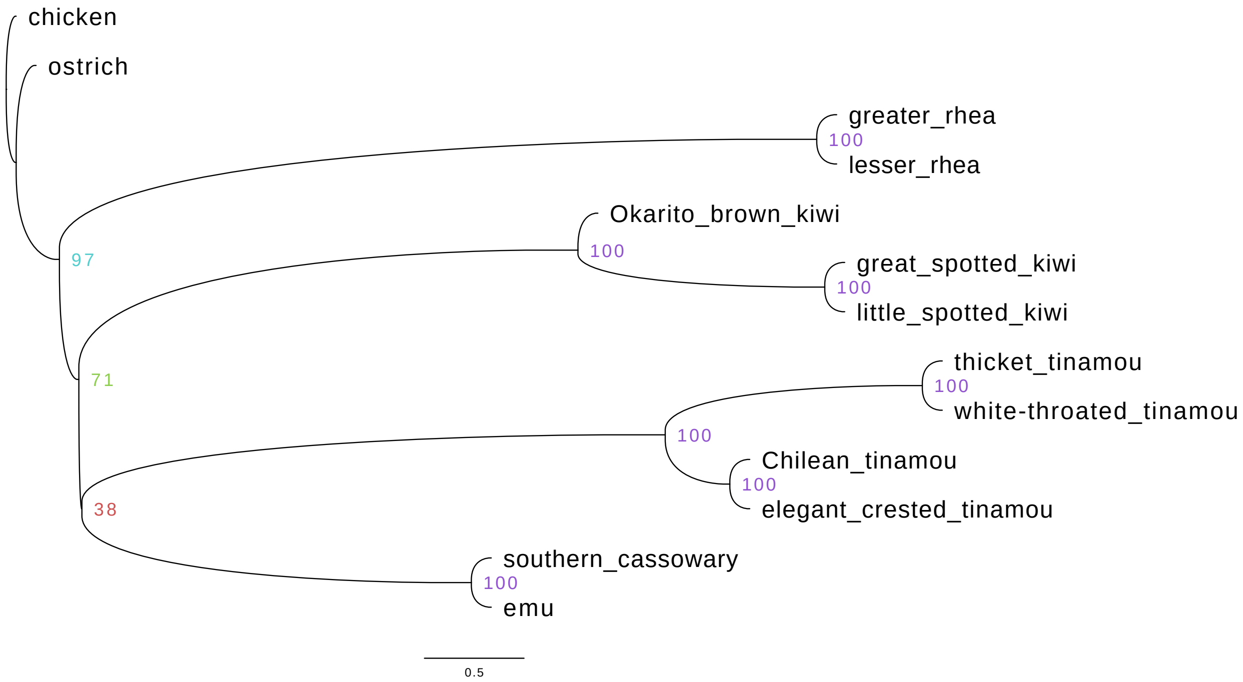

Figure S15: **ASTRAL_BP tree # 8 for Palaeognathae-GT-AZ data set with 1 000 RIs.** For the Palaeognathae-GT-AZ simulated data sets with 1 000 RIs, ASTRAL_BP recovered a unique incorrect topology (shown above) for replicate # 2. The *(incorrect) branch separating tinamou and emu+cassowary* from the remainder of the taxa has support (local PP) of 0.3767 and length (MLE-RI) of 0.0153 CUs. The $EN$ around this branch is 64.1042, and the three quartet topologies around this branch have (normalized) frequencies of 0.3435, 0.3409, and 0.3156. The *(correct) branch separating tinamou, emu+cassowary, and kiwi* from the remainder of the taxa has support (local PP) of 0.7105 and length (MLE-RI) of 0.0976 CUs. The $EN$ around this branch is 73.3333, and the three quartet topologies around this branch have (normalized) frequencies of 0.3981, 0.2765, and 0.3254. Note that the estimated species tree is in the anomaly zone.

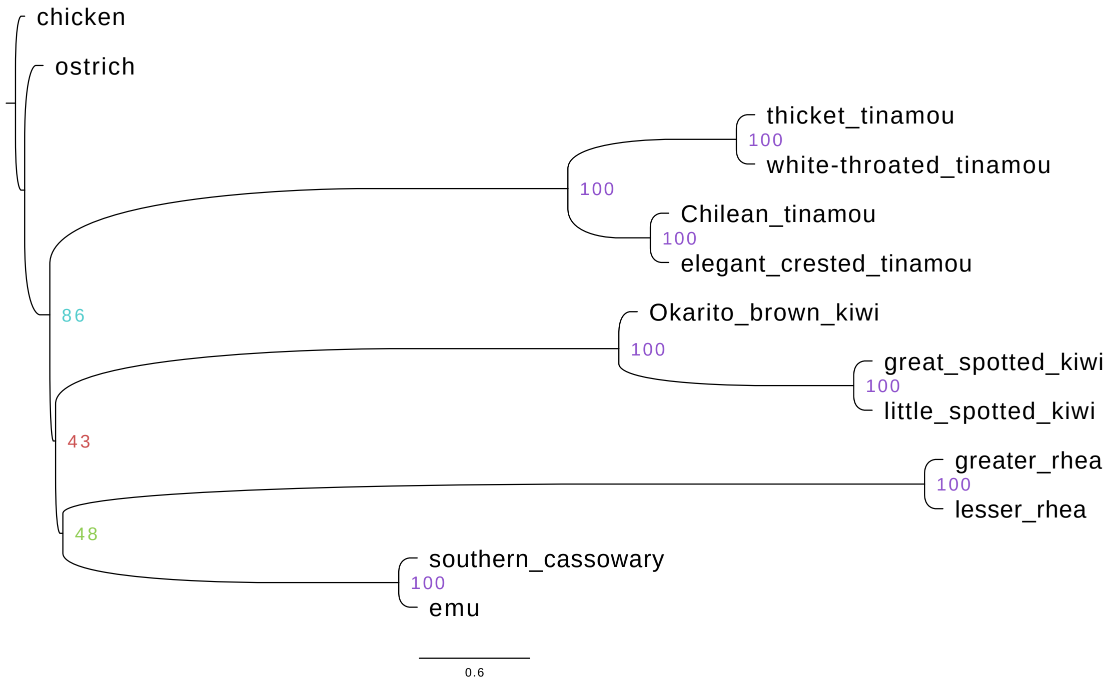

Figure S16: **ASTRAL_BP tree # 9 for Palaeognathae-GT-AZ data set with 1 000 RIs.** For the Palaeognathae-GT-AZ simulated data sets with 1 000 RIs, ASTRAL_BP recovered a unique incorrect topology (shown above) for replicate # 8. The *(incorrect) branch separating rhea and emu+cassowary* from the remainder of the taxa has support (local PP) of 0.4829 and length (MLE-RI) of 0.0397 CUs. The $EN$ around this branch is 78.1389, and the three quartet topologies around this branch have (normalized) frequencies of 0.3598, 0.3143, and 0.3260. The *(incorrect) branch separating rhea, emu+cassowary, and kiwi* from the remainder of the taxa has support (local PP) of 0.4335 and length (MLE-RI) of 0.0313 CUs. The $EN$ around this branch is 73.8750, and the three quartet topologies around this branch have (normalized) frequencies of 0.3542, 0.3418, and 0.3040. Note that the estimated species tree is in the anomaly zone.

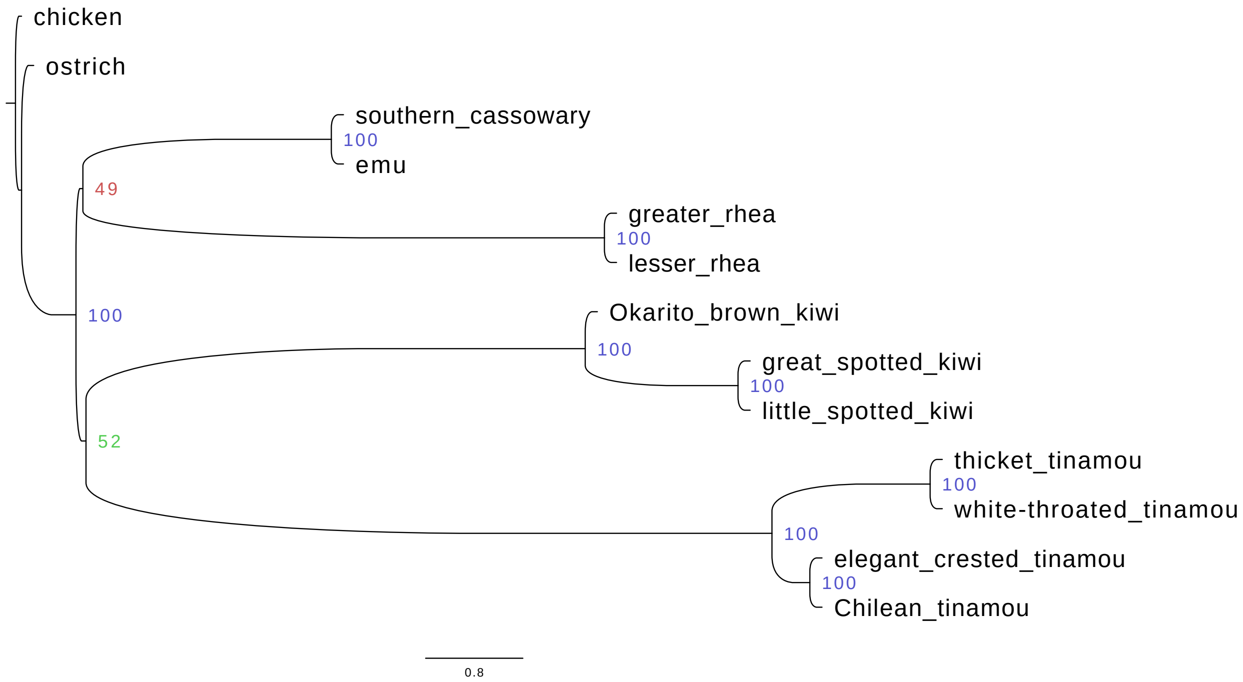

Figure S17: **ASTRAL_BP tree # 10 for Palaeognathae-GT-AZ data set with 1 000 RIs.** For the Palaeognathae-GT-AZ simulated data sets with 1 000 RIs, ASTRAL_BP recovered a unique incorrect topology (shown above) for replicate # 10. The *(incorrect) branch separating rhea and emu+cassowary* from the remainder of the taxa has support (local PP) of 0.4902 and length (MLE-RI) of 0.0569 CUs. The $EN$ around this branch is 52.8214, and the three quartet topologies around this branch have (normalized) frequencies of 0.3712, 0.2826, and 0.3462. The *(incorrect) branch separating kiwi and tinamou* from the remainder of the taxa has support (local PP) of 0.5182 and length (MLE-RI) of 0.0824 CUs. The $EN$ around this branch is 58.6250, and the three quartet topologies around this branch have (normalized) frequencies of 0.3881, 0.2388, and 0.3731.

#### 8 Non-Uniform Prior for Local Posterior Probability

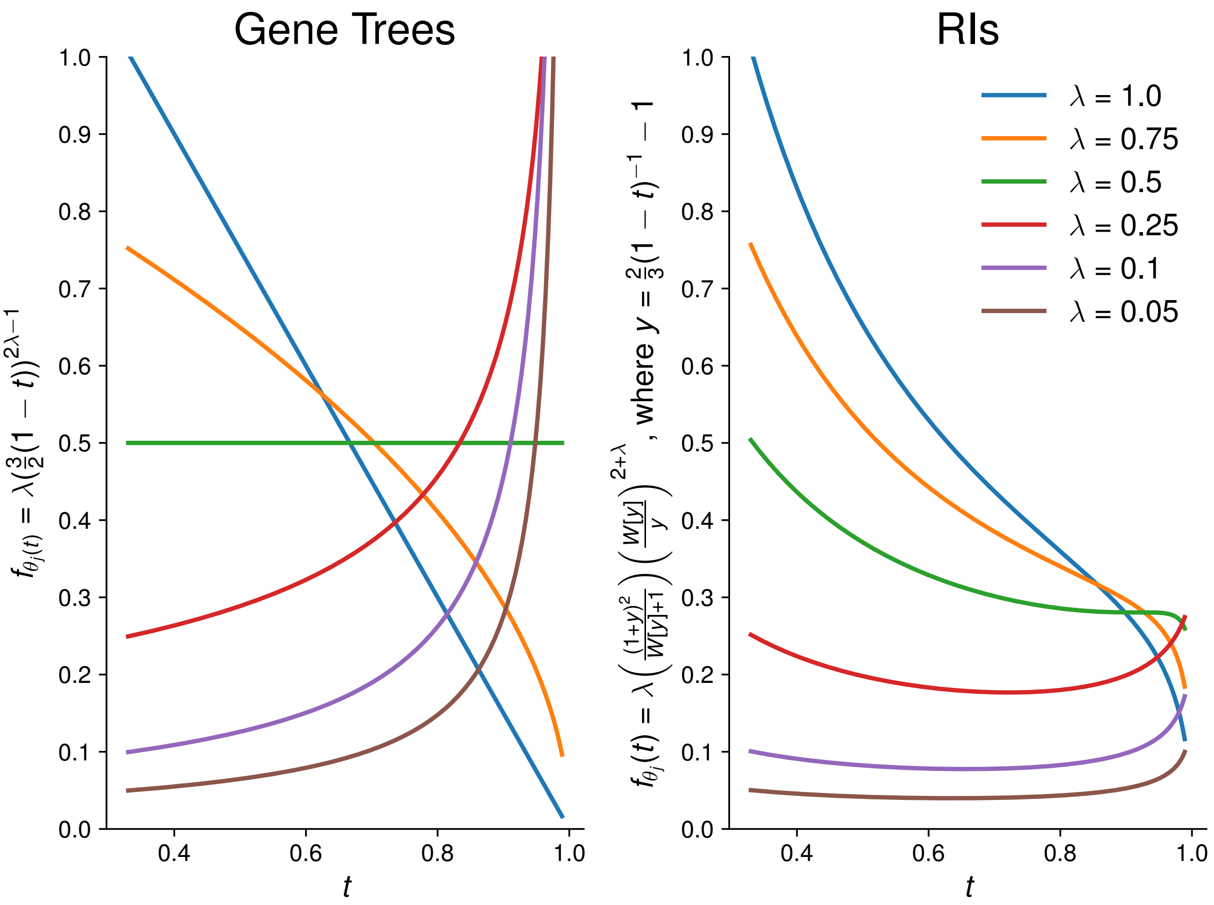

Figure S18: **Prior on probability of dominant quartet when species tree is generated under a Yule process with birth rate $\lambda$.** The equation for gene trees comes from Lemma 2 in Sayyari and Mirarab (2016), and the equation for retroelement insertions (RIs) is given in the Appendix. Note that unlike the prior for gene trees, the prior for retroelements is not uniform prior for $\lambda = 0.5$.
